# The impact of insertion bias into piRNA clusters on the invasion of transposable elements

**DOI:** 10.1101/2024.10.06.616898

**Authors:** Shashank Pritam, Almorò Scarpa, Robert Kofler, Sarah Signor

## Abstract

In our current understanding of transposable element (TE) invasions TEs move freely until they accidentally insert into a piRNA cluster. They are then silenced by the production of piRNA cognate to the TE. Under this model, one would expect that TEs might evolve to avoid piRNA clusters. Yet empirical observations show that some TEs, such as the *P* -element, insert into piRNA clusters preferentially. We were thus wondering if such a bias could be beneficial for the TE, for example by minimizing harm to the host while still being able to selfishly spread in populations. We decided to model insertion bias to determine if there was ever a situation in which insertion bias was beneficial to the TE. We performed extensive forward simulations of TE invasions with differing insertion biases into piRNA clusters. We found that insertion bias significantly altered the invasion dynamics of TEs, primarily by changing the copy number of the TE in individuals prior to silencing. Insertion into a piRNA cluster reduced the deleterious effects of TEs to the host population, but we found that TEs avoiding piRNA clusters out-compete TEs with a bias towards cluster insertions. Insertion bias was only beneficial to the TE when there was negative selection against TEs and a lack of recombination. Different TEs show different insertion biases into piRNA clusters suggesting they are an attribute of the TE not the host, yet scenarios in which this is beneficial to the TE are quite limited. This opens up an interesting area of future research into the dynamics of insertion bias during TE invasions.

**Significance Statement:** This study challenges the pre-existing understanding of the TE dynamics by investigating the potential adaptive role of insertion bias into piRNA clusters. Using extensive forward simulations, we demonstrate that while insertion bias significantly alters TE invasion dynamics, it is generally not beneficial for the TE’s spread in populations. Our results also show that transposable elements (TEs) that avoid piRNA clusters tend to do better than those that are more likely to preferencially insert into these clusters. This work provides novel insights into the complex dynamics between TEs and host genomes, showing the limited scenarios where the insertion bias could be advantageous to TEs. These results open new area for research into TE invasion dynamics and the evolution of host-TE interactions, further contributing to our understanding of genome evolution and stability.

## Introduction

Diverse transposable elements (TEs) make up a substantial fraction of eukaryotic genomes, ranging from 20% in *Drosophila* to 90% in maize [Goubert et al., 2015, Hill, 2019, Mérel et al., 2020]. These elements selfishly increase in copy number causing genomic instability in the form of double stranded DNA breaks, ectopic recombination, and disruption of coding sequences [Bourque et al., 2018]. Given that the majority of TE insertions are deleterious it was previously hypothesized that TEs maintain their copy number through a balance between transposition and negative selection [Charlesworth and Langley, 1986b, Nuzhdin and Mackay, 1995, Nuzhdin et al., 1997]. However, Brennecke (2007) found that TEs are in fact suppressed by a dedicated small RNA pathway [Brennecke et al., 2007b]. Small RNA termed *piwi* RNA (piRNA) are produced by TE-rich genomic regions and these piRNA are then bound by Argonaute class proteins which silence TEs pre and post-transcriptionally [Darricarrère et al., 2013, Gunawardane et al., 2007].

These TE-rich genomic regions which produce piRNA are discrete and are referred to as piRNA clusters. piRNA clusters are generally found in the heterchromoatin, near the euchromatic boundary [Brennecke et al., 2007a]. They make up a substantial proportion of the genome, for example in *D. melanogaster* piRNA clusters are 3.5% of the total genome. They are made up of dense TE insertions varying from recently active full length TEs to small degraded fragments of much older invasions. Several studies have found that a single insertion of a TE into a cluster region was sufficient to initiate silencing of a TE [Ronsseray et al., 1991, Josse et al., 2007, Zanni et al., 2013].

The observation that a single TE insertion into a piRNA cluster silenced the TE led to the development of the ‘trap’ model of TE suppression - under this model an invading TE jumps into a piRNA cluster, which triggers the emergence of piRNAs complementary to the TE [Bergman et al., 2006, Malone et al., 2009, Zanni et al., 2013, Goriaux et al., 2014, Yamanaka et al., 2014, Ozata et al., 2019]. This prevents the TE from further transposition. If TEs are suppressed under the trap model, several expectations should be met - piRNAs should be produced from sequences inserted in piRNA clusters, insertion into a piRNA cluster should be sufficient to suppress a TE, and TEs should not be present in many copies within piRNA clusters [Bergman et al., 2006, Malone et al., 2009, Zanni et al., 2013, Goriaux et al., 2014, Yamanaka et al., 2014, Josse et al., 2007].

Simulations of TEs invasions under the trap model have revealed additional expectations that can be empirically tested. For example, these simulations have established that TEs are initially silenced by segregating cluster insertions, and that overall around four cluster insertions in a population are necessary to stop a TE invasion [Kelleher et al., 2018, Kofler, 2019a, Scarpa et al., 2023]. TE invasions proceed through three stages - rapid, where the TE is proliferating uncontrolled in the host genome, shotgun, where cluster insertions are segregating in the population but remain unfixed, and inactive [Kofler, 2019a, Scarpa et al., 2023]. Existing work largely meets these expectations and supports the trap model of TE suppression [Muerdter et al., 2012, Luo et al., 2023, Kawaoka et al., 2012, Josse et al., 2007, Brennecke et al., 2007a, Zanni et al., 2013, Wierzbicki et al., 2023, Wierzbicki and Kofler, 2023].

However, there are some observations that do not fit with the expectations of the trap model. For example, there are fewer cluster insertions than expected [Scarpa and Kofler, 2023, Kofler et al., 2018, 2022, Selvaraju et al., 2022]. In addition, deleting the three main piRNA clusters in *D. melanogaster* did not have an effect on TE activity [Gebert et al., 2021]. The ‘trap’ model of TE suppression also supposes that TEs insert randomly within the genome, however there is some evidence that TEs insert preferentially into piRNA producing regions. For example, Kofler [2020] found that piRNA clusters must be a certain % of the genome to effectively suppress transposons. Yet, some species have piRNA clusters that do not meet this minimum size, without suffering the consequences of uncontrolled TE transposition. An insertion bias into piRNA clusters could compensate for small piRNA clusters. In fact some TEs do show evidence of insertion bias, such as the *P* -element which inserts preferentially into a piRNA cluster called *X-TAS* [Kelleher et al., 2018, Kofler, 2019a]. Investigations of novel insertions revealed that several TE families could have an insertion bias toward piRNA clusters [Khurana et al., 2011]. A high insertion rate of recently invading TEs was also observed for flamenco, i.e. the piRNA cluster of the soma [Zanni et al., 2013]. An insertion bias into piRNA cluster may be an evolutionary strategy employed by the TE. Such a bias could allow the TE to accumulate a sufficient number of TE copies in an organism to ensure efficient transmission to the next generation, yet prevent the accumulation of excessive copies that could harm the host. Previous work investigated whether TEs may evolve self-regulation through reducing the transposition rates to minimize damage to the host [Charlesworth and Langley, 1986a]. A reduced transposition rate would solely evolve under a few scenarios, such as low recombination rates. Thus, we aimed to investigate the effect of an insertion bias into piRNA clusters on the invasion dynamics of TEs and to test whether such an insertion bias could be an adaptive strategy employed by the TE.

To investigate the possibility that TEs have an insertion bias into piRNA clusters, we wanted to simulate different scenarios in which an insertion bias could potentially be beneficial to the TE. The goal of the present work is to determine with simulations whether there is a scenario in which evolving an insertion bias is beneficial to the TE in terms of copy number in the population. Using our simulator, InvadeGo, we show that an insertion bias into piRNA clusters is generally not beneficial to the TE.

## Results

### Model implementation and assumptions

If TE insertions are essentially random with regard to the whole genome, a positive insertion bias will lead to more insertions in piRNA clusters than expected by chance. For example, in the absence of insertion bias, the probability that a TE will insert into a piRNA cluster is determined by the amount of the genome that the piRNA cluster occupies - if that is 3% then that is also the probability of a cluster insertion. If a TE has an insertion bias then the probability to insert into a piRNA cluster is *>* 3%.

In our simulations, an insertion bias is a characteristic of the TE, not the host. We assume that a TE is active in all individuals that do not have an insertion into a piRNA cluster (Figure 1A). This assumption aligns with the “trap model” proposed in previous studies, where the proliferation of an active TE is halted when one copy inserts into a piRNA cluster, subsequently deactivating all TE copies in trans [Bergman et al., 2006, Malone et al., 2009, Zanni et al., 2013, Goriaux et al., 2014, Yamanaka et al., 2014, Ozata et al., 2019].

**Figure 1:**
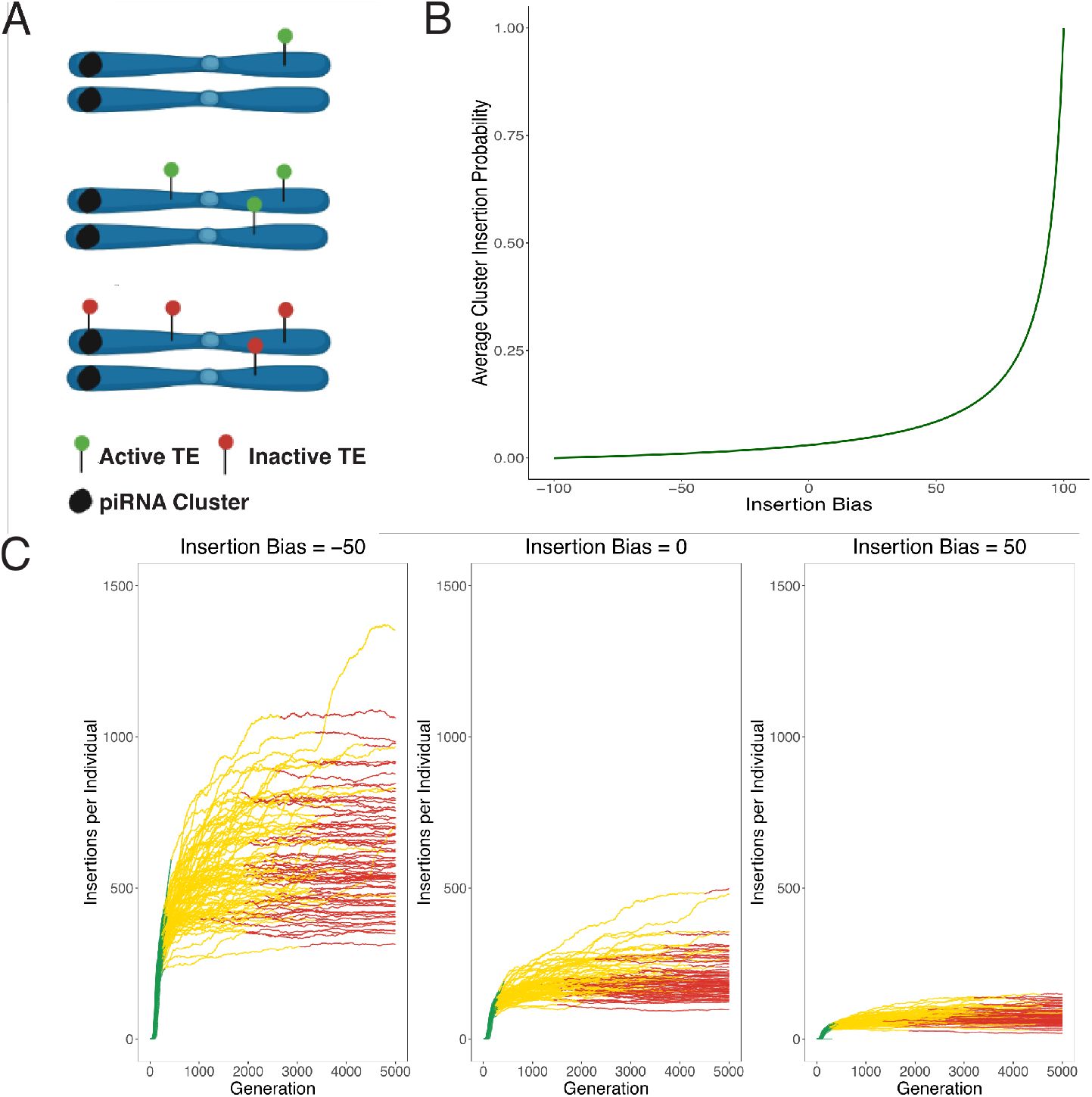
TE invasion modeling **(A)** A simple overview of our model assumptions. We begin the simulation with TE insertions in the population to avoid loss due to drift. The TE increases in copy number until the inserts into a piRNA cluster and is silenced. **(B)** Relationship between insertion bias and probability of TE integration within piRNA cluster versus other genomic regions. **(C)** Effect of insertion bias on TE abundance during invasion phases (color-coded). The three phases are rapid, shotgun, and inactive and are discussed further in the text. Higher bias into cluster regions correlates with reduced TE accumulation.

The piRNA clusters modeled here are based on dual-stranded germline clusters, where TE insertions can be in any orientation and generate piRNAs that silence TEs [Malone et al., 2009]. This model is supported by findings that individual euchromatic TE insertions can trigger the formation of novel dual-stranded piRNA clusters, which contributes to a more effective defense against TE expansion in Drosophila. The dual-stranded nature of these clusters is crucial for their function, as it allows for the production of both sense and antisense piRNAs, boosting the silencing capacity against active TEs [Shpiz et al., 2014]. It is also important to note that the size and distribution of piRNA clusters play a significant role in their effectiveness against TE invasions.

In the context of our simulations, these findings highlights the complexity of TE-piRNA cluster interactions and the importance of considering factors such as insertion bias, cluster size, and spatial organization when modeling the dynamics of TE invasions and their suppression by piRNA clusters. In the first set of simulations performed here TE insertions are assumed to be selectively neutral. There were two reasons for the approach. First, we are investigating the behavior of a complex system and the simplest possible scenario should be initially explored before adding additional complicating factors. However, the fitness effect of many TE insertions is also controversial - while it is unlikely that a system such as the piRNA pathway would have evolved without a negative fitness effect of TEs, there is ambiguous evidence that individual TE insertions are necessarily deleterious [Arkhipova, 2018, Blumenstiel et al., 2014, Mackay, 1989]. For example, we expect the X chromosome to have fewer TE insertions than the autosomes if they are negatively selected because the X chromosome is directly exposed to selection in males. However, in *Drosophila* the X chromosome does not show different patterns of TE insertions relative to the autosomes [Petrov et al., 2011, Kofler et al., 2015]. Furthermore, ectopic recombination could be the source of negative fitness effects from TE insertions, but there isn’t strong evidence of a relationship between recombination rate and TE density outside of *Drosophila* [Quadrana et al., 2016, Kent et al., 2017, Laricchia et al., 2017].

Empirical work on TE invasions supports an alternative scenario, where TE invasions are halted by many segregating cluster insertions [Kelleher et al., 2018]. Other empirical work on the *P* -element also supports this scenario, where the invasion of the TE plateaued at around 20 generations, during which all observed cluster insertions were segregating at low frequency [Kofler et al., 2018].

The following parameters were used as a default for all simulations unless specified - a transposition rate of 0.1, a population size of N=1,000, and piRNA clusters of 300 kb (3%) of the genome. We also used five chromosome arms of 10 Mbp each and a recombination rate of 4 cM/Mbp. An important base parameter is a starting population of 100 randomly inserted TEs in the population of 1,000. These insertions will have a population frequency of *f* = 1 / (2 * 1,000). Triggering a TE invasion with multiple insertions avoids early loss of TEs due to stochastic genetic drift [Scarpa and Kofler, 2023, Kofler, 2019a]. For every simulation we performed 100 replicates. We initially simulated TE insertions with no negative selection, but later incorporated scenarios with selection against TE insertions.

### Effect of insertion bias on TE invasions

Here we hypothesized that an insertion bias may be beneficial to the TE as it minimizes damage to the host while still enabling the TE to spread to appreciable copy numbers. We tested this with extensive forward simulations under the trap model, which assumes that a TE is spreading in a population until one copy jumps into a piRNA cluster 1A. An insertion in a piRNA cluster silences all copies of the TE. We modeled an insertion bias with values between -100 (complete avoidance of piRNA cluster) and +100 (all insertions in piRNA clusters). Values of 0 indicate no insertion bias. The insertion bias can be translated into the probability that a TE jumps into a piRNA cluster (see 1B). Note that in an unbiased case (bias=0) the probability of inserting into a piRNA cluster corresponds to the genomic proportion of the piRNA cluster (i.e. 0.03 in our simulations).

We first tested whether an insertion bias has an effect on the invasion dynamics of TEs. We performed 100 simulations for three values of insertion bias : -50, 0 and 50 (u=0.1; neutral insertions). Previous work established that TE invasions typically proceed through three phases - rapid, shotgun, and inactive [Kofler, 2019a]. In the rapid phase the TE spreads in the population unhindered by the host defense (Figure 1C (green)). During the shotgun phase (yellow), there are segregating piRNA cluster insertions that are controlling the spread of the TE but they have not reached fixation in the population. In the final inactive phase the population has fixed piRNA producing loci which are sufficient to entirely prevent transposition of the TE. When the first piRNA cluster with a TE insertion reaches fixation this phase begins. In the initial simulations there is no selection against transposition, so the piRNA cluster insertion reaches fixation through drift. Figure 1C illustrates the movement through these phases for three values of insertion bias. As expected we found that an insertion bias has a marked influence on invasion dynamics, where for example the number of insertions per individual decreases with the insertion bias 1C.

The first critical step after horizontal transfer of a novel TE to a naive population is establishment in the new population [Le Rouzic and Capy, 2005]. Especially at early stages of a TE invasion, when TE copy numbers are low, a newly invading TE may get easily lost by genetic drift. Since the probability of establishment decreases with the transposition rate (*p ≈* 2*u* where *u* is the transposition rate) self-regulation of TEs by limiting their activity will reduce the rate of establishment. Here we speculate that an insertion bias into piRNA clusters may be a form of self-regulation that avoids this problem, as the TE will initially (i.e in the absence of cluster insertions) have an uninhibited transposition rate. Only when the TE attains high copy numbers, i.e. is well established in the populations, cluster insertions will emerge that reduce the activity of the TE. We thus first tested whether the insertion bias affects the rate of establishment. The chances of establishment of a TE for invasions starting with a single segregating insertion are fairly low, which makes it hard to see further reduction due to an insertion bias. We thus started the simulations with 10 insertion to elevate the range of the observed values. We say that a TE is established if it persists for at least 500 generations. Interestingly we found that the insertion bias has little impact on the chances of establishment, unless the bias is very high (over 60-70, see: Figure 2). This suggests that moderate insertion biases into piRNA clusters (*<* 60) do not reduces the chances of a TE to get established in a population.

**Figure 2:**
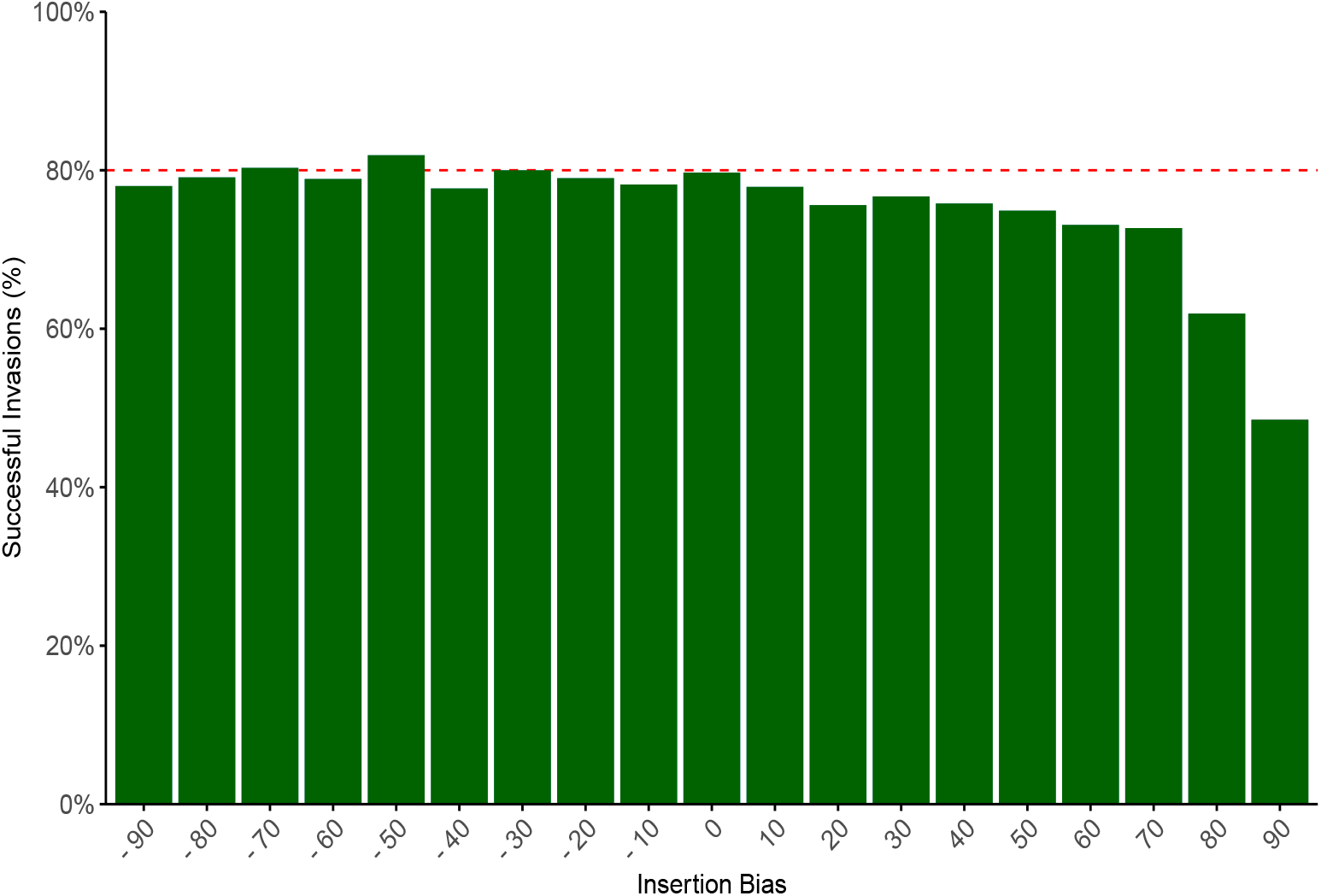
The effect of insertion bias on the probability that a TE will establish in the population. The dotted red line indicates the theoretical expectation that a TE will establish in a population across all simulations. Overall, insertion bias does not affect the likelihood of establishment unless insertion bias is quite large.

Next we examined the effect of an insertion bias on invasion dynamics in more detail (u=0.1, neutral etc, some parameters). We first noticed that the insertion bias has a substantial effect on the number of TEs accumulating during an invasion (Figure 3A). An increasing insertion bias leads to fewer TEs accumulating during the invasions (Figure 3A). Therefore the degree of bias determines the number of non-cluster insertions prior to silencing of the TE. This makes intuitive sense - as a TE is randomly inserting into the genome it will take more insertions to hit a piRNA cluster when there is negative bias towards piRNA clusters or no bias. This is not unexpected, since an insertion bias has conceptually a similar effect as increasing the size of piRNA clusters. Both increasing insertion bias and larger piRNA clusters increase the likelihood that a TE will jump into a piRNA cluster. Previous work revealed that the number of TE insertions accumulating during TE invasions depends largely on the size of piRNA clusters (where large clusters lead to few TEs accumulating during an invasion) [Kofler, 2019a]. Therefore it is expected that an an increasing insertion bias has a similar effect as larger piRNA clusters. Next we investigated the effect of insertion bias on the length of the TE invasion phases. We hypothesized that an insertion bias into piRNA clusters would lead to quicker suppression of TE transposition, because it should take fewer total insertions before one occurs in a cluster and silences transposition. This is however not what we found. Figure 3 reveals that the average duration of the rapid and shotgun phases — the periods before TE inactivation — remains relatively constant across different bias levels (Figure 3A). While there is slight variation in values and ranges, the mean phase lengths are essentially similar. This is in agreement with previous work where the length of the phases was not significantly dependent on the size of piRNA clusters [Kofler, 2019a] (which is conceptually similar to an insertion bias). This can be explained by the fact that TE copy numbers at early stages of an invasion increase exponentially, such that cluster insertions will rapidly emerge in all simulated scenarios.

**Figure 3:**
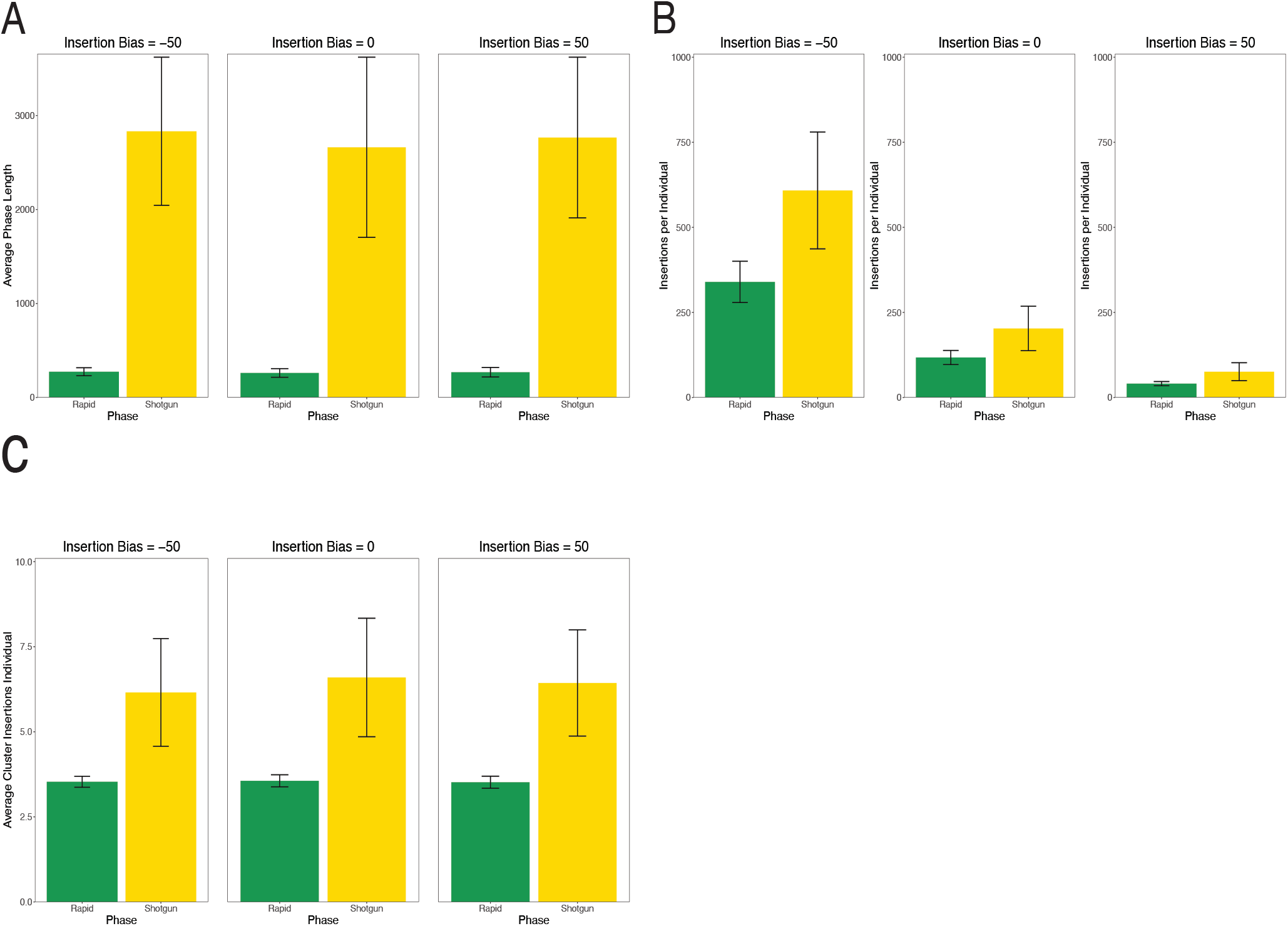
**(A)** The average length of the rapid and shotgun phases of TE invasion with different insertion biases. Note that we do not include the inactive phase as it has no clear termination point. **(B)** The average number of insertions per individual in the different phases of a TE invasion with different insertion biases. **(C)** The average number of cluster insertions per individual in the rapid and shotgun invasion phases under different insertion biases.

A similar observation (Figure 3C) can be made regarding the average number of cluster insertions per diploid individual; there is slight variance, but it does not vary significantly with bias. This is perhaps counter-intuitive as one might expect more cluster insertions with increasing insertion bias. But it needs to be considered that in our model the TE activity stops in individuals with one (or more) cluster insertions, thus preventing further accumulation of TE copy numbers. The number of cluster insertions necessary to stop an invasion remains about four, consistent with all previous simulations of TE invasions [Kofler, 2019a, Scarpa et al., 2023]. This is true regardless of the fact that a single insertion is sufficient to silence TE transposition. Recombination among cluster insertions results in a fraction of individuals that do not carry a cluster insertion and thus the TE is able to maintain low levels of activity in the population. This will increase the average number of cluster insertions until most individuals carry about four insertions. Changing the insertion bias of the TEs did not have an effect on the average number of cluster insertions necessary to halt a TE invasion. This is again consistent with previous work where the size of piRNA clusters did not have an effect on the number of cluster insertion [Kofler, 2019a].

To summarize, in neutral simulations an insertion bias decreases the number of TEs accumulating during an invasion but has little effect on the length of the phases or the number of cluster insertions (at later generations, when the TE is silenced by piRNAs).

### Insertion bias affects the fitness of the population

Under neutral conditions insertion bias reduces the total number of insertions per individual in a population. However, most TE insertions are likely either neutral or negatively selected, so we next asked how insertion bias might effect the fitness burden of TE invasions. In particular, we speculate that an insertion bias may reduce the fitness burden that TEs pose to hosts, which could then indirectly benefit the TE.

Classic literature on TE invasions published prior to the discovery of piRNA defense were able to show that negative selection has a substantial impact on the dynamics of TE invasions, controlling the invasion of the TE regardless of host silencing [Charlesworth and Charlesworth, 1983]. However, it has not been explored how negative selection will impact TE invasions in the presence of host silencing and insertion bias [Kofler, 2019b].

To explore this question, we introduced deleterious effects of TE insertions into our simulations. We simulated a linear fitness cost of TE insertions *w* = 1 *− x ∗ n* where *w* is the individual fitness, *n* is the number of insertions, and *x* is the fitness cost of individual insertions. Negative selection alters the invasion dynamics of TEs considerably compared to a neutral scenario [Kofler, 2019a]. Under neutrality, TE copy numbers increase rapidly in the population early in the invasion, followed by a plateau as more piRNA cluster insertions are introduced. Dependent on the extent of negative effects (*x*) three principal outcomes are feasible. First if negative effects are strong (*x > u*), a TE may not be able to invade since all copies are quickly purged from the population. Second if negative effects are small (*Ne ∗ x <* 1) than the invasion will resemble a neutral scenario. Third for intermediate values a TE will be able to spread in a population until the host defence and negative selection controls the TE (TSC-balance [Kofler, 2019a]). Figure 4A presents average fitness across generations, with standard deviations shown in lighter colors. Initially a TE will quickly multiply in a population lowering the average fitness. As piRNA cluster insertions arise and the TE is silenced, negative selection purges the TE insertions from the population and fitness recovers. Please note that piRNA cluster insertions were still subject to negative selection in this scenario, thus fitness does not recover to 1.

**Figure 4:**
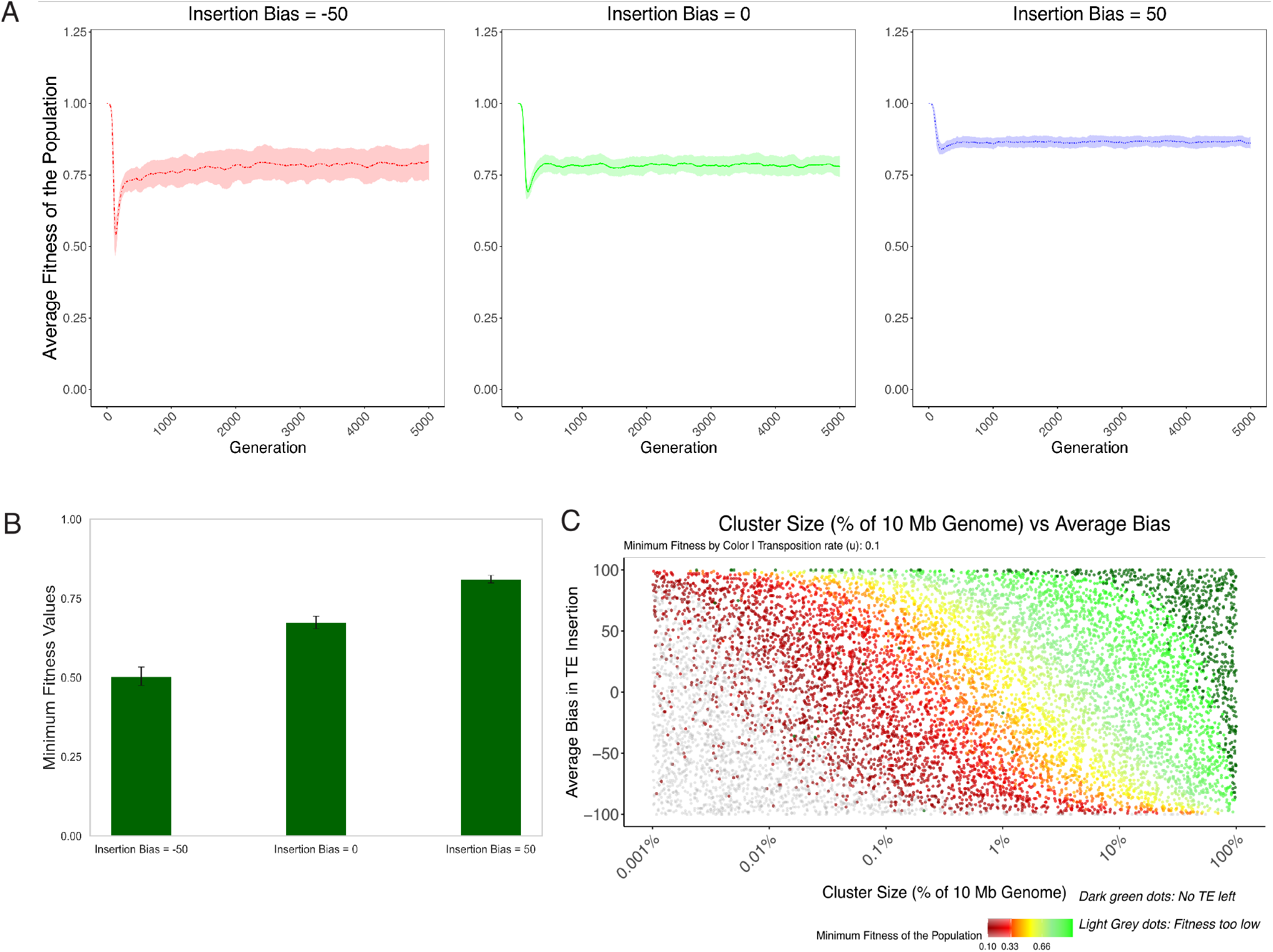
Fitness dynamics during TE invasions with varying insertion biases **(A)** Average population fitness over generations for different TE insertion biases. Lines show mean fitness; shaded areas represent standard deviations. **(B)** Minimum population fitness (*min w*) achieved during invasions for three TE bias levels. **(C)** Population fitness mapped against piRNA cluster size (x-axis) and average TE insertion bias (y-axis). Color indicates fitness value, ranging from dark red (lowest fitness, *min w <* 0.01) through red (*min w <* 0.1), yellow (*min w <* 0.33), to green (highest fitness, *min w* = 1). Dark green points indicate populations where no TEs are left (*fail −* 0), light grey points represent populations with fitness too low (*fail − w*, extinction).

In this context we refer to minimum fitness as the lowest fitness of individuals during the TE invasion. This can also be thought of as the maximum fitness burden of the TE during an invasion. This is an important parameter, given that it has been hypothesized that TE invasions could drive local population extinctions [Munasinghe et al., 2023b, Studer et al., 2011]. Our observations indicate that the insertion bias indeed reduces the maximum fitness burdens of TEs (Figure 4A). This is intuitive given our previous results which found that higher bias results in a lower total number of insertions in each individual. Without selection against TE insertions, higher bias results in fewer TE insertions per individual prior to silencing of the TE. When these TEs have a negative fitness cost, it results in a higher overall fitness and thus a lower fitness burden.

Figure 4B further illustrates this maximum TE burden by showing minimum fitness during invasions of TEs with three different biases. The lowest value corresponds to a -50 bias, increasing as bias increases. This demonstrates that lower bias is more costly for the population. A key finding, depicted in Figure 4C, explores population fitness in a 2D parameter space of piRNA cluster size (x-axis) and average TE insertion bias (y-axis), with fitness values color-coded. We found that for small piRNA clusters that the minimum fitness can drop below 0.1. We assume that these populations cannot persist and thus will go extinct. Confirming Kofler [2020] observation that piRNA clusters require a minimum size to control TE invasions. We note that increased insertion bias into piRNA clusters may compensate for smaller cluster sizes. Population fitness increases with cluster size and average bias, while negative bias leads to extinction even with large cluster sizes. For successful invasion, TEs must find a strategy above this “sweet spot” where the population survives.

In summary, we found that an insertion bias reduces the fitness burden of TEs to hosts. Furthermore a strong bias against piRNA clusters could lead to extinction of populations.

### Invasion dynamics of multiple TEs with different insertion bias

To increase in copy numbers TEs may employ two, mutually not exclusive, strategies. First, they may selfishly proliferate even if this reduces the fitness of the host. Second, they may impose some sort of self-regulation, thus reducing damage to the host. Hosts with higher fitness (i.e. less damage due to TEs) will rise in frequency and thus the TE will hitchhike with the host to higher copy numbers. Since both strategies have their pros and cons it is not intuitively clear which one will be best for proliferation of TEs. Here we speculate that an insertion bias into piRNA clusters may be beneficial for a TE, as it allows TEs to spread rapidly in a population (as long as copy numbers are low) but then when copy numbers are increasing cluster insertions will emerge that limit damage to the host.

To test this hypothesis we performed pairwise-competitions of TEs with two different insertion biases. We asked the question, under what genomic and evolutionary conditions might TEs with higher insertion bias towards piRNA clusters stabilize at higher copy number than those with lower bias? Hence we performed simulations with two different TEs that jointly invade in a population.

To trigger the invasions we introduced 100 copies of each TE at random positions. We assumed that both TEs have identical properties (transposition rate *u* = 0.1, negative effect) except for the bias into piRNA clusters. We further used a population size of *N* = 1000 and 100 replicates for each scenario.

Importantly we also assumed that insertion in a piRNA cluster silences both TEs. This is justified as we assume that insertion bias into a piRNA cluster may gradually evolve in the TE by mutations, and a few mutations may be sufficient to alter the insertion bias but they will not allow the TE to escape the host defence (e.g. piRNAs act broadly over a wide range of the TE). It has for example been argued that up to 10% sequence divergence are tolerated between piRNAs and the silenced TE [Schwarz et al., 2021, Post et al., 2014, Kotov et al., 2019]. After 500 generations we recorded the copy numbers of both TEs and scaled the value between -1 and 1, where -1 means all TEs in the population have a negative bias, +1 all TEs have a positive bias, and 0 that the number of TEs with a negative and positive bias is identical.

As a control, we started with neutral simulations. In this scenario an insertion bias into a cluster is of no benefit to the TE, since TE insertions have no adverse effects on host fitness. We thus expect that TEs that avoid piRNA clusters outcompete TEs with a higher preference for clusters. We modeled two scenarios, one with recombination (random assortment among five chromosomes and crossovers occurring at a rate of 4cM/Mbp) and one without recombination (a single non-recombining chromosome; Table 1). As expected in both scenarios we consistently observed that TEs with lower insertion bias towards piRNA clusters obtained higher copy number than those with higher bias.

**Table 1:** Overview of Competitive TE Invasion Simulations: Figure 5.

| Scenario | Selection | Genome Structure | Premise and Rationale |
| --- | --- | --- | --- |
| A | Neutral | 5 chr, 5 clusters | Baseline for complex genome with distributed insertion targets |
| B | Neutral | 1 chr, 1 cluster, 0 RR | Simplified genome with concentrated target, no recombination |
| C | Negative | 5 chr, 5 clusters | Purifying selection in complex genomic environment |
| D | Negative | 1 chr, 1 cluster, 0 RR | Extreme case: selection pressure, simple genome, no recombination |

*Note:* All scenarios: N = 1000, initial TE = 100, 100 reps, 500 gens, u = 0.1. Negative selection: x = 0.01. chr = chromosomes, RR = recombination rate.

Next we introduced negative selection against TEs (*x* = 0.01) and again performed simulations in a scenario with and without recombination. If our hypothesis holds (an insertion bias is beneficial for the TE) we expect that TEs with a bias attain higher copy numbers than TEs with a lower bias. We solely observed this in the scenario without recombination. In the more biologically relevant scenario (several recombining chromosomes) we consistently observed that TEs with lower insertion bias towards piRNA clusters obtained higher copy number than those with higher bias. This shows that our hypothesis that an insertion bias is beneficial to the TE does not hold, except for the scenario without recombination. This is in agreement with previous works suggesting that self-regulation of TEs could evolve in the absence of recombination [Charlesworth and Langley, 1986a].

In summary, we found that our hypothesis that an insertion bias into piRNA clusters may be beneficial to the TE does not hold, except in a scenario with negatively selected TEs in organisms without recombination. Our work suggests that TEs might evolve a bias to avoid piRNA clusters.

## Discussion

In this manuscript we explored the possibility that under different conditions in a population it could be beneficial to a TE to evolve an insertion bias into piRNA clusters. Our results, as illustrated in Figures 1-5, reveal a new insight about TE invasion dynamics and their impacts on host fitness.

While many TEs do not empirically appear to have an insertion bias into piRNA clusters, some TEs such as the *P* -element show a strong insertion bias. Recent studies have revealed that the relevance of insertion bias varies among different TEs and environmental conditions. The *P* -element in Drosophila has been shown to have a stronger insertion bias into telomere-associated sequences (TAS), which are important piRNA clusters, under hot conditions compared to cold conditions [Kofler et al., 2022]. Some somatic TEs, like *gypsy* in Drosophila, may have an insertion bias into specific piRNA clusters such as the flamenco locus [Kofler, 2019a]. In certain cases, the direction of TE insertion into piRNA clusters has been found to correlate with the sense/antisense bias in piRNA production, suggesting that insertion bias can influence piRNA-mediated defense mechanisms [Hirano et al., 2014].

While higher TE insertion rates into piRNA clusters have been observed in Drosophila, similar biases have not been consistently described in mammals, indicating potential differences in TE-host dynamics across species [Ernst et al., 2017]. The important role of the insertion preference in the invasion trajectory of transposons has been further emphasized by recent studies [Munasinghe et al., 2023a], building upon earlier work on the evolution of self-regulated transposition [Charlesworth and Langley, 1986a]. These findings, in essence, highlight the complexity and variability of TE insertion biases across different TE families, host species, and environmental conditions. The observed high insertion bias might confer unexpected benefits to both TEs and hosts. While TE insertions are generally considered costly to the host, a higher bias towards piRNA clusters could mitigate this cost by concentrating insertions in genomic regions already dedicated to TE regulation. This strategy could allow TEs to persist in the genome while minimizing disruption to essential host genes.

These findings suggest that the observed bias of *P* -elements towards *X-TAS* may represent an evolutionary strategy that balances the need for TE propagation with minimizing host damage. In genomic environments where silencing is efficient and recombination is limited, targeting piRNA clusters could provide TEs with a “safe harbor” or “sweet spot” for insertion, allowing them to persist in the population while potentially contributing to the host’s defensive repertoire. The important role of insertion preference in the invasion trajectory of transposons has been further emphasized by recent studies [Munasinghe et al., 2023a], building upon earlier work on the evolution of self-regulated transposition [Charlesworth and Langley, 1986a]. These findings, in essence, highlight the complexity and variability of TE insertion biases across different TE families, host species, and environmental conditions. This study not only sheds light on the specific case of *P* -elements and *X-TAS* but also broadens our understanding of the evolutionary forces shaping TE-host interactions across diverse genomic landscapes. The interplay between insertion biases, environmental conditions, and host defense mechanisms reveals a complex evolutionary “arms-race” between TEs and their hosts, with implications for genome evolution and the maintenance of genomic stability.

An insertion bias into a piRNA clusters is a form of self-regulation that was not previously explored [Charlesworth and Langley, 1986a]. We thought that an insertion bias into piRNA clusters may be an appealing strategy for the TE as it avoids several problems of other forms of self-regulation. In particular self-regulation of TE activity will lower the chances of establishment in a novel population. Reduced rate of establishment will threaten the long-term persistence of a TE. We showed that an insertion bias into piRNA clusters does avoids this problem, as the effect on the establishment is minor (Figure 2). We also demonstrated that higher insertion bias towards piRNA clusters correlates with reduced TE accumulation. The fitness dynamics presented in Figure 4 highlight the complex relationship between TE bias, piRNA cluster size, and host fitness, with higher biases typically resulting in less fitness reduction. This supports our idea that an insertion bias may reduce harm to the host. However, contrary to expectations our competition simulations (Figure 5) revealed that under most genomic contexts, lower-bias TEs obtain higher copy numbers than their high-bias counterparts. An insertion bias into piRNA clusters was solely beneficial (in terms of final copy numbers) in a scenario without recombination. This is in agreement with previous works showing that self-regulation of TEs might typically solely evolve in the absence of recombination [Charlesworth and Langley, 1986a]. Our work implies that in most organisms (recombining) TEs will evolve to avoid piRNA clusters. However we further showed that a bias against piRNA clusters will lead to elevated rates of host extinction, where the load of deleterious TEs cannot be kept in check by negative selection anymore.

**Figure 5:**
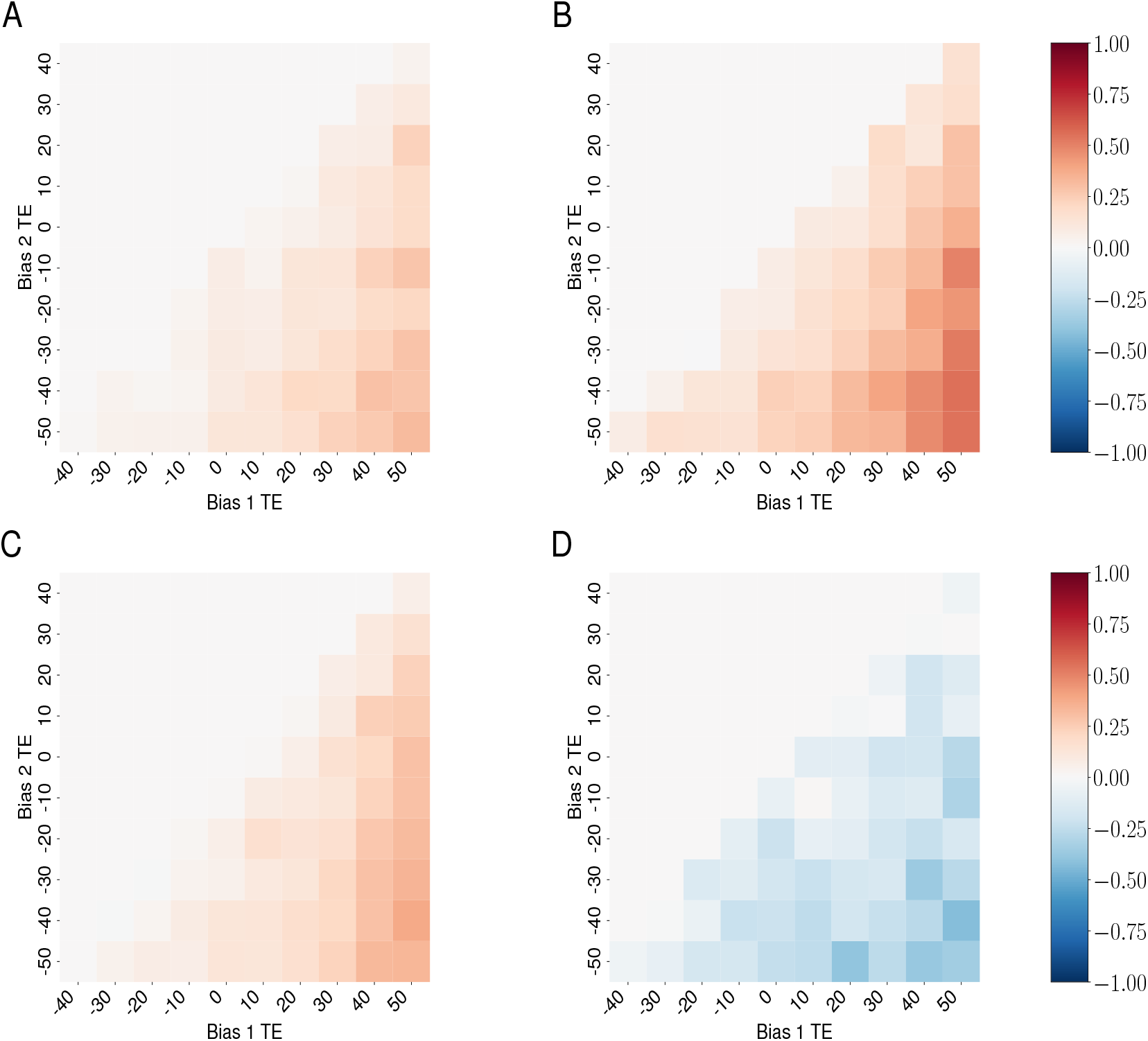
Competition dynamics between TEs with different insertion biases across varied genomic contexts. (A-D) correspond to scenarios detailed in Table 1. Color scale represents competitive outcomes: red indicates dominance of less biased TEs, blue shows dominance of more biased TEs, white represents equal competition, and grey dots indicate absence of both TE types. Scale calculated as 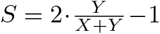, where X and Y represent average total insertions for more (x-axis) and less (y-axis) biased TEs, respectively.

This raises the question as to why more populations don’t go extinct from TE invasions. There are several possible explanations. First it is possible that an insertion into a piRNA cluster does not trigger the host defence. This hypothesis aligns with recent discoveries about the complexity and adaptability of piRNA-mediated defense systems. Studies [Gebert et al., 2021] have shown that even after the removal of three major piRNA clusters, TEs remained effectively silenced, suggesting a robust redundancy in the system. As an alternative it was suggested that siRNAs are mediating the conversion of TEs into piRNA producing loci. Also other forms of host defence may protect against extinctions such as KRAB-ZNFs or the hush silencing in humans. Second it is also possible that TEs cannot evolve to avoid piRNA clusters. Since piRNA clusters are very diverse (e.g. accounting for 3% of the genome) there may be few genomic or epigenomic cues that allows TEs to distinguish between cluster and non-cluster sites. Third, recent work by [Scarpa and Kofler, 2023] demonstrated the crucial role of paramutation, a mechanism distinct from piRNA clusters, in the dynamics of TE silencing. In the context of TEs, paramutations refers to the conversion of a regular TE insertions into piRNA producing loci. This process is typically mediated by maternally transmitted piRNAs. The emergence of abundant piRNA producing loci due to paramutations may prevent extinctions.

These insights highlight the complex co-evolutionary dynamics between TEs and their hosts, suggesting that what appears costly or parasitic at one level might confer unexpected benefits at another. Future research could focus on experimentally testing these hypotheses, perhaps by competing TEs with (*P* -element) and without insertion bias in experimental populations of model organisms and observing the long-term effects on both TE proliferation and host fitness across varying genomic architectures. Such studies could shed light on the intricate interplay between TE insertion preferences, piRNA cluster dynamics, paramutation, and the evolution of host genome defense mechanisms.

## Materials and methods

### Simulation software

To simulate TE invasions with insertion bias we developed a novel branch (“insertionbias”) for the previously developed simulation software (Invadego (v0.1.3)) [Scarpa and Kofler, 2023]. This software performs individual-based forward simulations of TE invasions in populations of diploid organism using discrete and non-overlapping generations. Every TE insertion has two properties, i) a genomic position (integer) in the half-open interval [0, *g*), where *g* is the genome size and ii) and an insertion bias (byte) into piRNA clusters. Note that it is thus possible to simulate TEs with different insertion biases in the same genome. The TE insertions in a haploid genome are represented as a dictionary where the position acts as key and the bias as value. Thus a diploid individual carries two separate dictionaries of TE insertions. Each chromosome occupies a unique non-overlapping territory in the genomic interval [0, *g*), where every TE insertion is part of exactly one chromosome. piRNA clusters occupy sub-regions of each chromosome. TE insertions may be a part of none or one piRNA clusters. We opted to model the insertion bias as a discrete integer value from -100 to +100 (represented as a single byte, to minimize memory consumption), where 0 is unbiased, -100 is a strong bias against piRNA clusters (no insertions in piRNA clusters) and +100 is a strong insertion bias into piRNA clusters (all insertions are in piRNA clusters). The probability of a novel TE inserting into a piRNA cluster (*p*_*c*_) can be computed from the bias (*b*) and the genomic proportion of piRNA clusters (*f*). For example if piRNA clusters account for 3.5% of the genome, as in Drosophila, then *f* = 0.035.

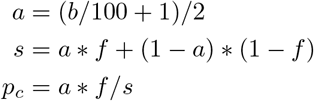

The resulting probability (*p*_*c*_) will be a value between 0 and 1. Note that in the absence of an insertion bias (*b* = 0) the probability to insert into a piRNA cluster is identical to the genomic fraction of the piRNA cluster (*p*_*c*_ = *f*).

Each individual has a fitness *w*, which solely depends on the number of TE insertions *w* = 1 *− xn*, where *x* is the negative effect of a single TE insertion and *n* is the number of TE insertions per diploid individual. Simulations with neutral TE insertions can be performed using *x* = 0. The fitness determines the mating probability (i.e. fecundity selection). We simulated hermaphrodites that may randomly mate with other hermaphrodites. Each parent generates a single gamete that is passed to the offspring. To create a gamete, first recombination and random assortment among chromosomes are simulated and then novel transposition events are introduced into the recombined gamete. We assumed that TEs multiply with a given transposition rate *u*, which is the probability that a TE insertion will generate a novel insertion in the next generation. A transposition rate of zero (*u* = 0) was used for individuals carrying an insertion in a piRNA cluster. To avoid excessive computation times, we calculated the number of novel insertion sites for each gamete based on a Poisson distributed random variable with *λ* = *u ∗ n/*2. Based on the probability of jumping into a piRNA cluster (*p*_*c*_ see above), we randomly distributed novel insertions either within or outside of piRNA clusters. If a site was already occupied, the novel insertion was ignored.

Our software allows the user to provide a wide range of different parameters such as the number of chromosomes, the size of the chromosomes, the size of the piRNA clusters, the recombination rate, the transposition rate, the population size, the number of generations, the number of TE insertions in the base population, the negative effect of TEs and a flag indicating whether or not cluster insertions are selectively neutral. For the base population it is possible to provide a file with the position and the bias of the TE insertions.

The novel tool was thoroughly tested with unit-tests. We further validated the correct implementation of our software to confirm that it correctly models population forces such as recombination, drift, and selection (Supplementary Figs. S1–S7). For example, theoretically a proportion of TE insertions should reach fixation due to genetic drift depending upon the population size. These expectations were met and all of the simulations performed to validate the model are described in the supplement. Additionally, we verified that the software accurately models insertion bias as specified, illustrated in Figure 1B. The simulations, analysis, and figures for visualization from this work have been documented and deposited on GitHub. 92–100% of the invasions were stopped after 5,000 generations and all after 10,000 generations (Supplementary Table S1; Fig. 1C)

### Simulations and data analysis

For simulations we have used several default conditions - five chromosome arms of 10 Mbp each, a recombination rate of 4cM/Mbp, piRNA clusters of 300 kb (3% of the genome), a population size of 1000, transposition rate of 0.1, and a base population with 100 randomly inserted TEs. The last parameter is to avoid losing TEs to genetic drift [Scarpa and Kofler, 2023, Kofler, 2019a, Le Rouzic and Capy, 2005]. For every simulation we performed 100 replicates. We initially simulated TE insertions with no negative selection, but later incorporated scenarios with selection against TE insertions.

The output of all of the simulations was visualized in R using ggplot2 [Wickham, 2016], Seaborn [Waskom, 2021], and matplotlib [Hunter, 2007]. Simulations output a large amount of data therefore we also used DuckDB [Raasveldt and Mühleisen, 2019] for data management.

## Data Availability

Invadego insertion module is available at GitHub: https://github.com/RobertKofler/invadego/tree/insertionbias. The population genetics validations are documented at: https://github.com/shashankpritam/Insertion-Bias-TE

## Acknowledgments

SS would like to thank F. T. and S. M. Emery for thoughtful commentary on this work. SP would like to thank his parents for their constant love, support and invaluable presence in his life.

## Author contributions

SP performed simulations and data analysis, as well as drafting and revising the manuscript. RK conceived of the manuscript, wrote InvadeGO and assisted in data analysis and writing. AS assisted with conception of the manuscript and development of the simulator. SS provided funding, assisted in interpreting the data, and drafted the manuscript.

