## Supplementary material for "The impact of insertion bias into piRNA clusters on the invasion of transposable elements": Suppliment Data

### Supplementary Data

Shashank Pritam

October 6, 2024

13 Analysis of Insertion Bias: Box plot showing the number of TE insertions within piRNA clusters for different insertion bias values. The x-axis represents the insertion bias, ranging from 0 to 90 and then from -10 to -90, with a step size of 10. The y-axis represents the number of TE insertions within piRNA clusters. . 15
14 Validation of selection effects: Relative frequency of TEs in clusters (clu) and non-clusters (noclu) at generations 0 and 50 for different selection scenarios. Left: Selection on all TEs. Right: Selection only on non-cluster TEs. . . . . . . . . . . . . . . . . . 16
15 Validation of selection effects: Distribution of TEs in clusters (red) and non-clusters (blue) across the genome at generations 0, 25, and 50 for a single replication. The y-axis represents the count of TEs per 10kb bin, while the x-axis shows the genomic position from 0 to 1Mb. The piRNA clusters are defined in the initial part of the genome (left side). . . . . . . . . . . . . . . . . 16
16 Validation of selection effects: Relative frequency of TEs in clusters (clu) and non-clusters (noclu) at generations 0 and 100 for different selection scenarios. Left: Selection on all TEs. Right: Selection only on non-cluster TEs. . . . . . . . . . . . . . . . . . 17
17 Validation of selection effects: Progression of TE population (y-axis) through generations (x-axis) for different selection coefficients (x) on the same population. Top left: x = 0, top middle: x = 0.1, top left: x = 0.01, bottom right: x = 0.001, bottom left: x = 0.0001. . . . . . . . . . . . . . . . . . . . . . . . . . . . . 18
18 Validation of selection effects: Progression of TE population frequency through generations for x = 0. The blue line represents the average TE frequency. . . . . . . . . . . . . . . . . . . . . . . 19
19 Validation of selection effects: Progression of TE population frequency through generations for x = 0.1. The blue line represents the average TE frequency. . . . . . . . . . . . . . . . . . . . . . . 19
20 Validation of selection effects: Progression of TE population frequency through generations for x = 0.01. The blue line represents the average TE frequency. . . . . . . . . . . . . . . . . . . . . . . 20
21 Validation of selection effects: Progression of TE population frequency through generations for x = 0.001. The blue line represents the average TE frequency. . . . . . . . . . . . . . . . . . . . 20
22 Validation of selection effects: Progression of TE population frequency through generations for x = 0.0001. The blue line represents the average TE frequency. . . . . . . . . . . . . . . . . . . . 21
23 Validation of selection effects: Progression of TE population frequency through generations for different selection coefficients (x) on the same population, with average TE frequency represented by blue lines. Top left: x = 0, top right: x = 0.1, middle left: x = 0.01, middle right: x = 0.001, bottom: x = 0.0001. . . . . . . . 21

24 Validation of insertion: Average cluster insertion as a function of insertion bias. The x-axis represents the insertion bias ranging from -100 to 100 with a step size of 10. The y-axis shows the average cluster insertion. The dots in the plot represent the observed values, while the orange line represents the expected values. The observed and expected values match, showing the steep exponential curve. . . . . . . . . . . . . . . . . . . . . . . . 22
25 Validation of insertion: Variability in TE insertions across different levels of insertion bias, based on data from 100 replications. The y-axis represents the relative difference between 'avcli' and 'pc' columns, expressed as a percentage. The x-axis represents the categorical 'Insertion Bias' variable. Each boxplot summarizes the distribution of 'TE_insertions' for each level of 'Insertion Bias', indicating the median, interquartile range, and potential outliers. . . . . . . . . . . . . . . . . . . . . . . . . . . . . . . . 23
26 Validation of insertion: Mean of average cluster insertion across various insertion bias levels, with error bars indicating standard deviation. The orange line represents the expected values. . . . . 24
27 Validation of insertion: The same data as Figure 26, but with a logarithmic transformation applied to the y-values. The log scale helps to better visualize the variability in the data (shown by the error bars) and the deviation of the observed values (blue points) from the expected values (orange line). . . . . . . . . . . . . . . . 25
28 Validation of insertion: Average TE insertions (y-axis) and insertion bias (x-axis) for a single replicate. The x-axis represents the insertion bias varying from -100 to 100, while the y-axis depicts the average TE insertions for each bias level. Each data point represents the average TE insertions for a specific bias level for replicate 1. . . . . . . . . . . . . . . . . . . . . . . . . . . . . . 26
29 Validation of insertion: All replicates' mean average TE insertions (y-axis) against the insertion bias (x-axis). The plot includes error bars representing the standard deviation for the average TE insertions at each bias level. The x-axis represents the insertion bias, ranging from -100 to 100, while the y-axis depicts the mean of the average TE insertions for each bias level. . . . . 27

### List of Tables

1 Across the 21 insertion bias values (sampleid 0 to 20), the mean of cluster insertion is calculated for all 100 replications. The standard deviation is also calculated. We have another column named pc which is the theoretical value of average cluster insertion for a given insertion bias with a 3% cluster size. . . . . . . . . . . . . . 28
2 Probability (p) that a TE jumps into a piRNA cluster depending on the insertion bias of the TE. The piRNA clusters accounts for 3% of the genome. Additionally, we show the number of expected insertions and observed with our software invade-insbias for 1000 new TE insertions. Observed values are averaged over 1000 replicates. This demonstrates that our software correctly implements the insertion bias. . . . . . . . . . . . . . . . . . . . . . . 29

### 1 Supplementary Figures

This section presents the supplementary figures related to the validation steps described in the manuscript.

#### Invasion

The purpose of this validation is to show that the simulator generates, on average, the number of insertions predicted by the following equation:

$$c_t = c_0(1 + \mu)^t \tag{1}$$

where:

- $c_t$: number of TE copies at generation $t$
- $c_0$: number of TE copies at generation 0
- $\mu$: transposition rate
- $t$: generation number

The initial conditions for the simulation are:

$$c_0 = 10$$
$$\mu = 0.1$$
$$t = 100$$

The simulation is performed on a chromosome of size 1 Mb without any piRNA clusters. A total of 500 replicates are used.

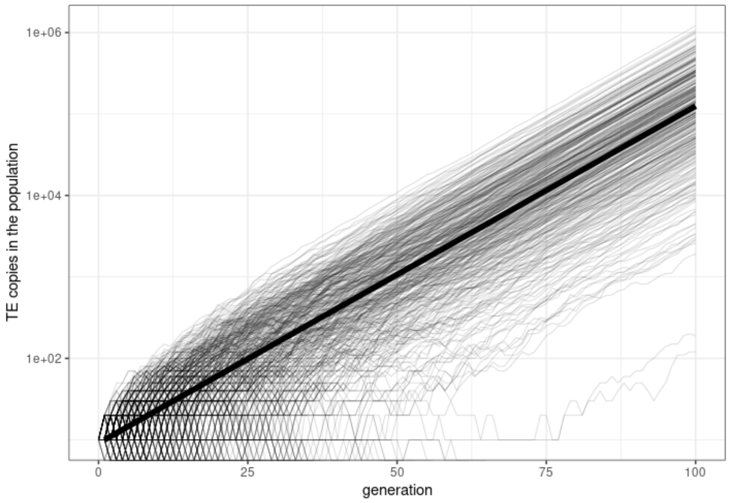

Figure 1: Validation of invasion dynamics: In this simulation, we validated that the transposable element (TE) insertion in the entire population followed the expected exponential growth based on theoretical and mathematical predictions. While most invasion replicates adhered to the expected trajectory (represented by the thick black line), some replicates also show variation due to stochastic effects.

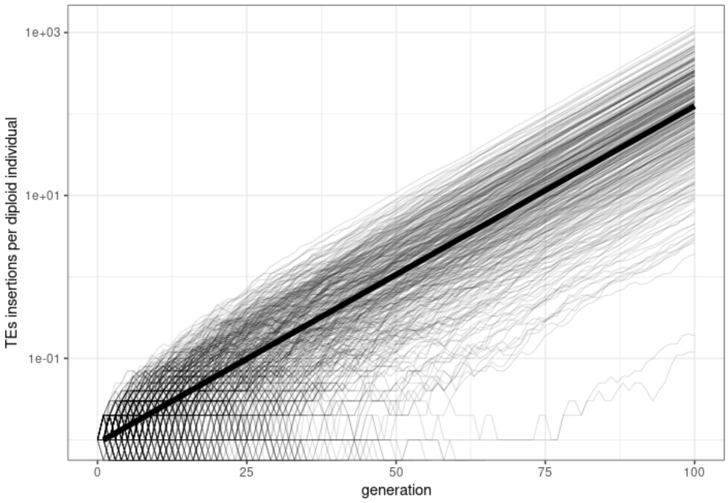

Figure 2: Validation of invasion dynamics: Similar to the previous simulation, we validated that the transposable element (TE) insertion in the individual organism followed the expected exponential growth based on theoretical and mathematical predictions. While most invasion replicates adhered to the expected trajectory (represented by the thick black line), some replicates also exhibited variation due to stochastic effects.

#### Drift

To test if genetic drift is simulated correctly, we exploit a fundamental population genetics principle: the probability of fixation of a neutral singleton in a diploid organism is given by:

$$P(\text{fix}) = \frac{1}{2N_e} \tag{2}$$

where $N_e$ is the effective population size. In the simulation results, the x-axis represents the number of fixed transposable elements (TEs) within the population, while the y-axis shows the count of replicates. We expect the distribution of fixed TEs to follow the probability of fixation given by Equation (1).

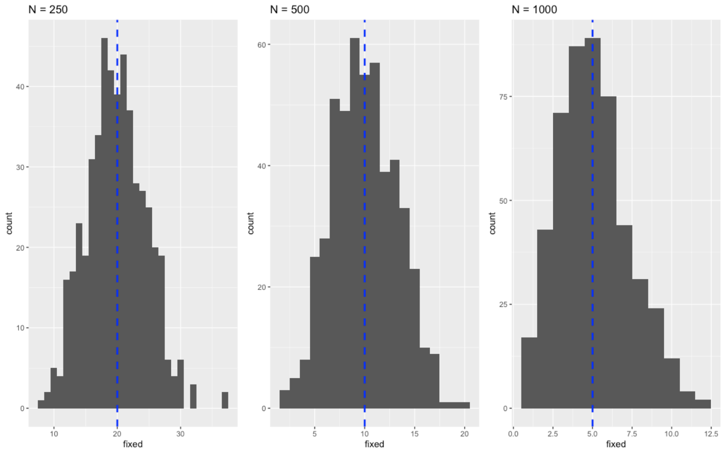

Figure 3: Validation of genetic drift: Distribution of fixed TEs in the population. The x-axis shows the number of fixed TEs, and the y-axis shows the count of replicates.

As shown in Figure 3, the observed distribution of fixed TEs in the simulated population aligns with the expected probability of fixation, confirming that genetic drift is accurately simulated in the model.

#### piRNA clusters

In this study, we consider three different scenarios based on the proportion of the genome composed of piRNA clusters:

- pc0: no piRNA clusters
- pc50: 50% of the genome is composed of piRNA clusters
- pc100: 100% of the genome is composed of piRNA clusters

##### Starting Conditions

The following starting conditions are used for all simulations:

$$c_0 = 10$$
$$\mu = 0.1$$
$$t = 100$$

The simulations are performed on a chromosome of size 1 Mb with variable piRNA cluster sizes.

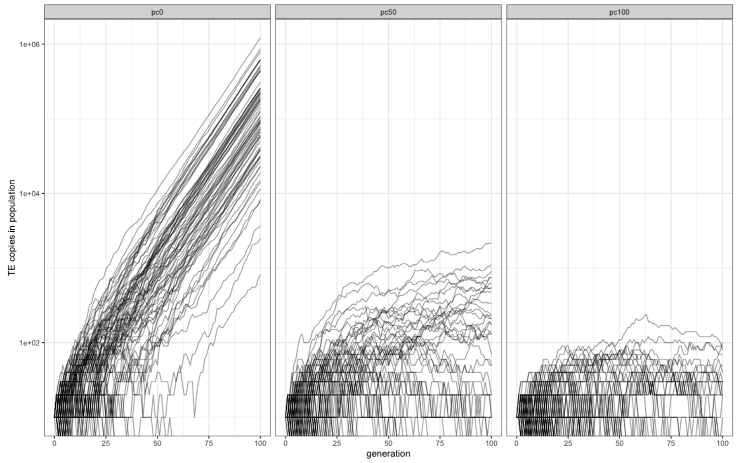

Figure 4: Validation of piRNA cluster: TE copies in the entire population with respect to generation for different piRNA cluster sizes

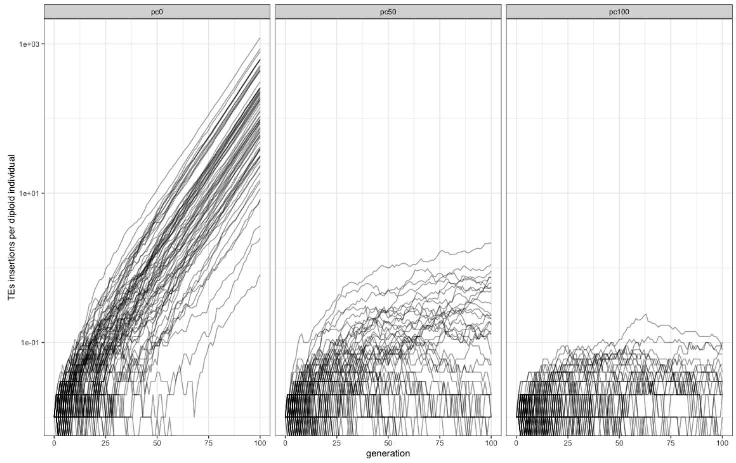

Figure 5: Validation of piRNA cluster: TE copies per diploid individual with respect to generation for different piRNA cluster sizes.

Figures 4 and 5 show the effect of piRNA clusters on the spread of transposable elements (TEs) in the population. Figure 4 depicts the total number of TE copies in the entire population, while Figure 5 shows the average number of TE copies per diploid individual. Both figures demonstrate that the introduction of piRNA clusters effectively suppresses the proliferation of TEs over generations. As the proportion of the genome composed of piRNA clusters increases (from pc0 to pc100), the spread of TEs is increasingly inhibited, showing the important role of piRNA clusters in regulating TE activity.

#### Recombination

This validation aims to test the correct implementation of recombination using linkage disequilibrium decay. Linkage disequilibrium (LD) is the non-random association of alleles at different loci in a population. The decay of LD is influenced by factors such as inbreeding and recombination rate. The number of generations needed to reach $D = 0$ is described by the equation:

$$D_n = (1 - c)^n D_0 \tag{3}$$

where $n$ is the number of generations, $D_n$ is the LD value at generation $n$, $D_0$ is the initial LD value, and $c$ is the recombination rate.

##### Scenarios

All validations use a population of $N = 10000$ and an initial TE distribution of 1000, simulated for 150 generations.

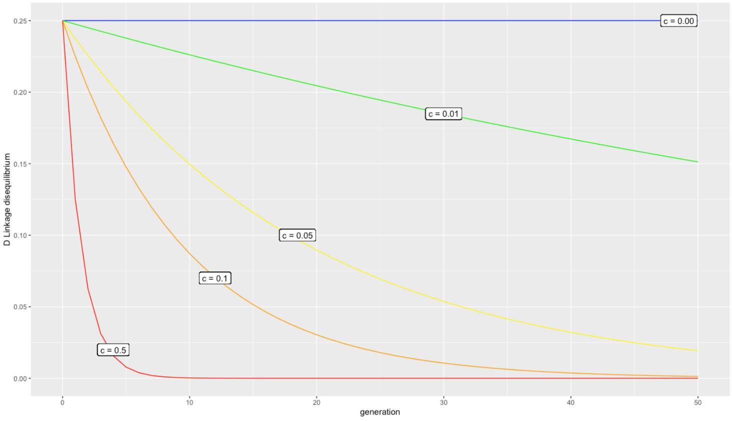

Figure 6: Validation of recombination dynamics: Expected LD decay for different recombination rates.

##### Validation Results

- Validation 7 A: Recombination rate = 0.00
- Validation 7 B: Recombination rate = 0.01
- Validation 7 C: Recombination rate = 0.05
- Validation 7 D: Recombination rate = 0.1

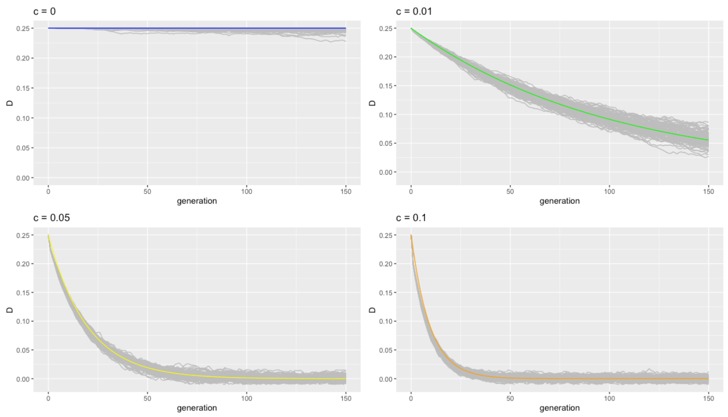

Figure 7: Validation of recombination dynamics: Observed LD decay for different recombination rates..

The observed LD decay (Figure 7) closely matches the expected decay (Figure 6) for each recombination rate, confirming the correct implementation of recombination in the simulation.

#### Insertion Bias

This simulation aims to understand the effect of insertion bias on the invasion dynamics of transposable elements (TEs).

##### Initial Conditions

The simulation uses the following initial conditions:

- Population size: 1000
- Number of chromosomes: 5
- Chromosome size: 10 Mb
- Number of piRNA clusters: 5
- piRNA cluster size: 300 Kb
- Initial number of TEs in the population: 10

For the establishment probability simulation (Part A), 1000 replicates were used.

#### Results and Observations

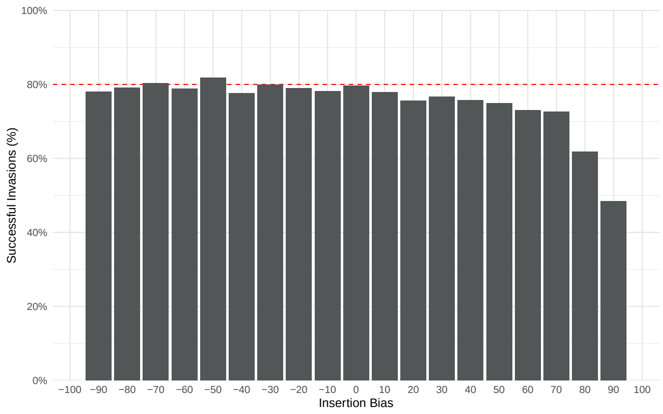

Figure 8: Analysis of Insertion Bias: Percentage of successful TE invasions as a function of insertion bias. The x-axis represents the insertion bias ranging from -90 to 90, while the y-axis shows the percentage of successful TE invasions.

Figure 8 shows the relationship between insertion bias and the percentage of successful TE invasions. The x-axis represents the insertion bias, ranging from -90 to 90, while the y-axis shows the percentage of successful TE invasions.

The main observation from this simulation is that the probability of successful TE invasion remains relatively constant (around 80%) unless the insertion bias is extremely high (over 60-70).

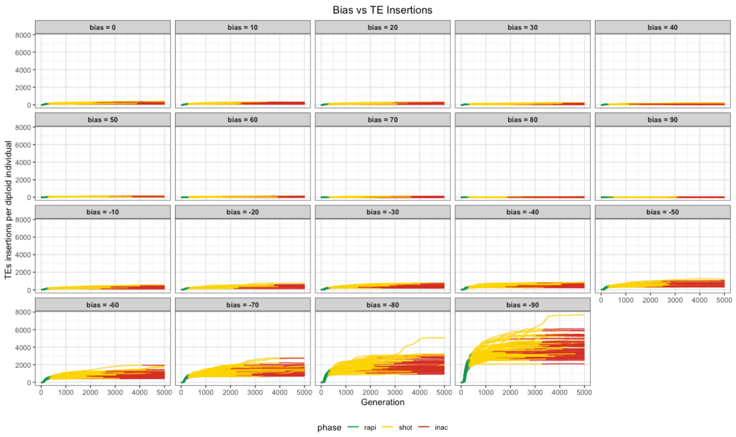

Figure 9: Analysis of Insertion Bias: TE insertions per diploid individual over generations for different insertion bias values. Each panel represents a specific insertion bias ranging from -90 to 90.

Figure 9 shows the number of TE insertions per diploid individual over generations for different insertion bias values, ranging from -90 to 90. Each panel in the figure represents a specific insertion bias value. The main observation from this simulation is that as the insertion bias moves from -90 to +90, the number of TE copies per individual decreases. This suggests that positive insertion bias (towards piRNA clusters) leads to a reduction in the overall TE load within the population.

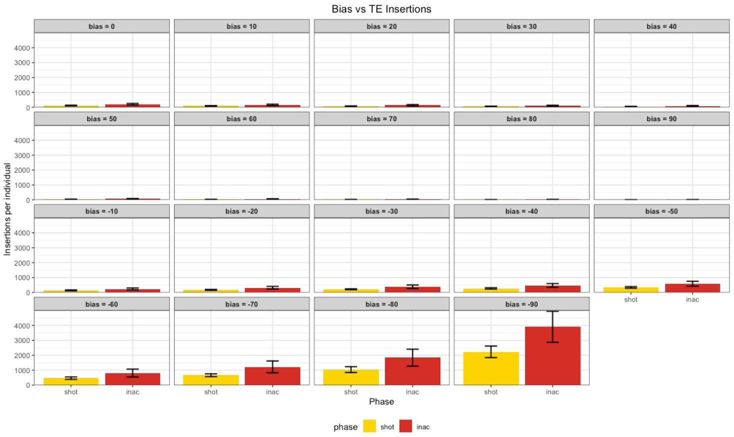

Figure 10: Analysis of Insertion Bias: TE insertions within piRNA clusters at different phases for various insertion bias values. The bar plot shows the number of TE insertions in piRNA clusters before the shotgun (rapid) and before the inactive (shotgun) phases. Each panel represents a specific insertion bias value ranging from -90 to 90.

Figure 10 presents a bar plot displaying the number of TE insertions per individual at two distinct phases: before the shotgun phase (rapid) and before the inactive phase (shotgun). The plot is divided into panels, each representing a specific insertion bias value ranging from -90 to 90.

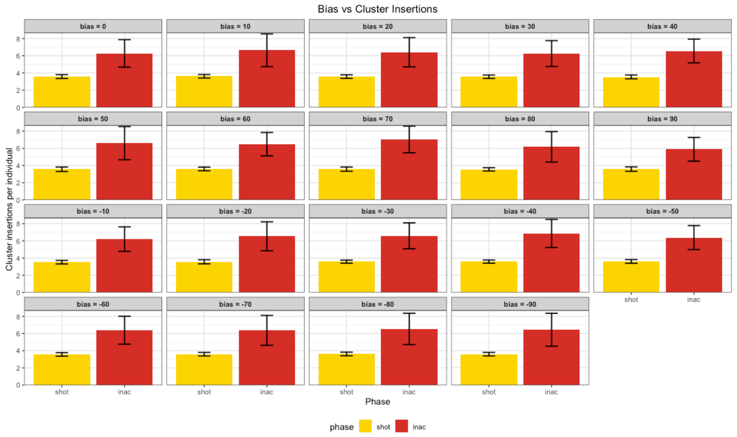

Figure 11: Analysis of Insertion Bias: TE insertions within piRNA clusters at different phases for various insertion bias values. The bar plot shows the number of TE insertions in piRNA clusters before the shotgun (rapid) and before the inactive (shotgun) phases. Each panel represents a specific insertion bias value ranging from -90 to 90.

Figure 11 presents a bar plot displaying the number of TE insertions within piRNA clusters at two distinct phases: before the shotgun phase (rapid) and before the inactive phase (shotgun). The plot is divided into panels, each representing a specific insertion bias value ranging from -90 to 90. We observe that the number of TE insertions within piRNA clusters remains consistent across different insertion bias values within each panel.

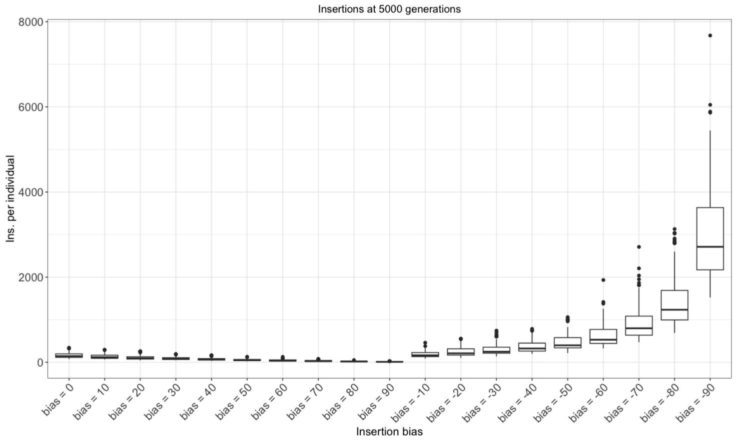

Figure 12: Analysis of Insertion Bias: Box plot showing the number of TE insertions per individual for different insertion bias values. The x-axis represents the insertion bias, ranging from 0 to 90 and then from -10 to -90, with a step size of 10. The y-axis represents the number of TE insertions per individual.

Figure 12 presents a box plot illustrating the relationship between insertion bias and the number of TE insertions per individual. The x-axis represents the insertion bias, ranging from 0 to 90 and then from -10 to -90, with a step size of 10. The y-axis represents the number of TE insertions per individual.

Starting from an insertion bias of 0, the median number of TE insertions per individual gradually decreases as the bias increases towards 90. And as the insertion bias becomes negative, starting from -10, the median number of TE insertions per individual begins to rise again. This trend continues until the insertion bias reaches -90.

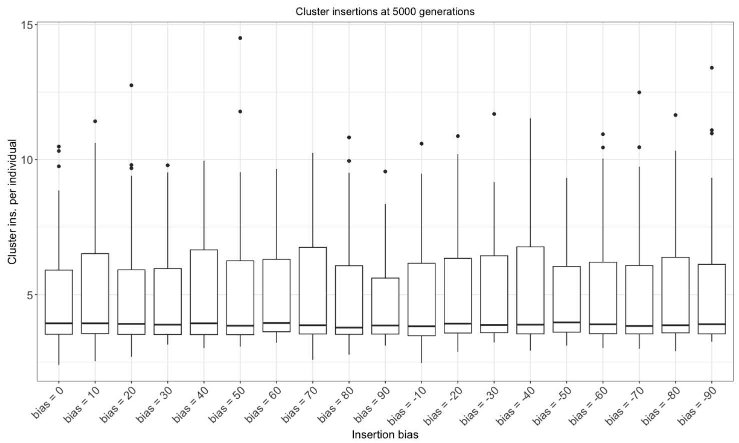

Figure 13: Analysis of Insertion Bias: Box plot showing the number of TE insertions within piRNA clusters for different insertion bias values. The x-axis represents the insertion bias, ranging from 0 to 90 and then from -10 to -90, with a step size of 10. The y-axis represents the number of TE insertions within piRNA clusters.

Figure 13 presents a box plot illustrating the relationship between insertion bias and the number of TE insertions within piRNA clusters. The x-axis represents the insertion bias, ranging from 0 to 90 and then from -10 to -90, with a step size of 10. The y-axis represents the number of TE insertions within piRNA clusters.

The box shows trend in the number of TE insertions within piRNA clusters as the insertion bias varies. Starting from an insertion bias of 0, the median number of TE insertions within piRNA clusters remains steady as the value of bias changes.

In this validation, we aimed to test if selection was correctly implemented in the simulation. To do so, we tested different scenarios: selection on all TEs versus selection on non-cluster TEs.

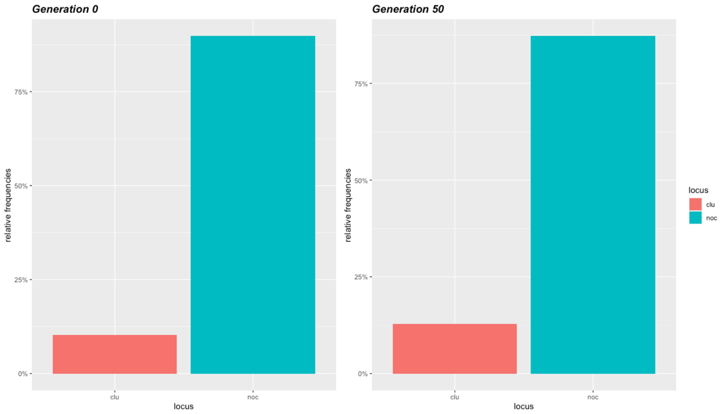

Figure 14: Validation of selection effects: Relative frequency of TEs in clusters (clu) and non-clusters (noclu) at generations 0 and 50 for different selection scenarios. Left: Selection on all TEs. Right: Selection only on non-cluster TEs.

Figure 14 shows two side-by-side bar plots comparing the relative frequency of TEs in clusters (clu) and non-clusters (noclu) at generations 0 and 50 for different selection scenarios. The plot on the left shows the results when selection is applied to all TEs, while the plot on the right displays the outcome when selection is only applied to non-cluster TEs. In both scenarios, the relative frequencies of TEs in clusters and non-clusters increase slightly from generation 0 to generation 50. Also, the overall ratio between the frequencies of TEs in clusters and non-clusters remains relatively constant across generations in both selection scenarios.

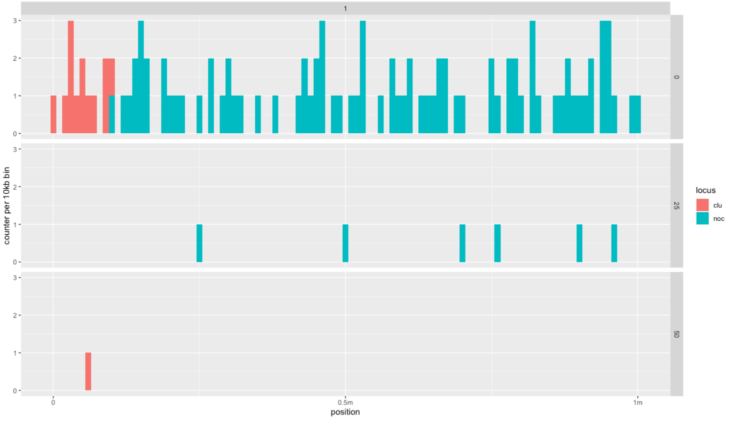

Figure 15: Validation of selection effects: Distribution of TEs in clusters (red) and non-clusters (blue) across the genome at generations 0, 25, and 50 for a single replication. The y-axis represents the count of TEs per 10kb bin, while the x-axis shows the genomic position from 0 to 1Mb. The piRNA clusters are defined in the initial part of the genome (left side).

Figure 15 show three stacked bar plots, each representing a different generation (0, 25, and 50) for a single replication of the simulation. The y-axis indicates the count of TEs per 10kb bin, while the x-axis represents the genomic position from 0 to 1Mb. The red bars denote TEs within piRNA clusters, and the blue bars represent TEs in non-cluster regions.

At generation 0, we observe a high abundance of both red and blue bars, indicating a substantial number of TEs in both cluster and non-cluster regions. The blue bars (non-cluster TEs) appear to be more numerous compared to the red bars (cluster TEs).

As we progress to generation 25, there is a noticeable change in the distribution of TEs. The blue bars are predominantly located on the right side of the plot, suggesting a shift in the distribution of non-cluster TEs towards the latter part of the genome. The red bars, representing cluster TEs, are no longer visible.

Finally, at generation 50, we observe a single red bar on the left side of the plot, corresponding to the initial part of the genome where piRNA clusters are defined in the simulation. This suggests that the cluster TEs have been largely eliminated, with only a small remnant remaining in the designated cluster region. The initial abundance of both cluster and non-cluster TEs at generation 0, followed by the shift in non-cluster TEs towards the right side of the genome at generation 25, and the eventual elimination of most cluster TEs by generation 50, shows the complex interplay between TE activity, piRNA-mediated silencing, and selection pressures.

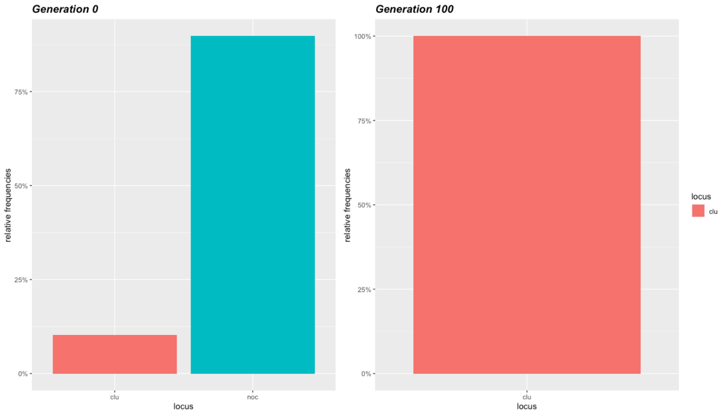

Figure 16: Validation of selection effects: Relative frequency of TEs in clusters (clu) and non-clusters (noclu) at generations 0 and 100 for different selection scenarios. Left: Selection on all TEs. Right: Selection only on non-cluster TEs.

Figure 16 presents two side-by-side bar plots comparing the relative frequency of TEs in clusters (clu) and non-clusters (noclu) at generations 0 and 100 for different selection scenarios. The plot on the left shows the results when selection is applied to all TEs, while the plot on the right displays the outcome when selection is only applied to non-cluster TEs. At generation 0, both plots exhibit a similar distribution of TEs in clusters and non-clusters, with a considerable presence of both red bars (cluster TEs) and blue bars (non-cluster TEs). This indicates that at the initial stage, TEs are distributed evenly across both cluster and non-cluster regions of the genome. However, at generation 100, a difference emerges between the two selection scenarios. In the plot, the red bars representing cluster TEs dominate the distribution, while the blue bars representing non-cluster TEs have diminished to zero.

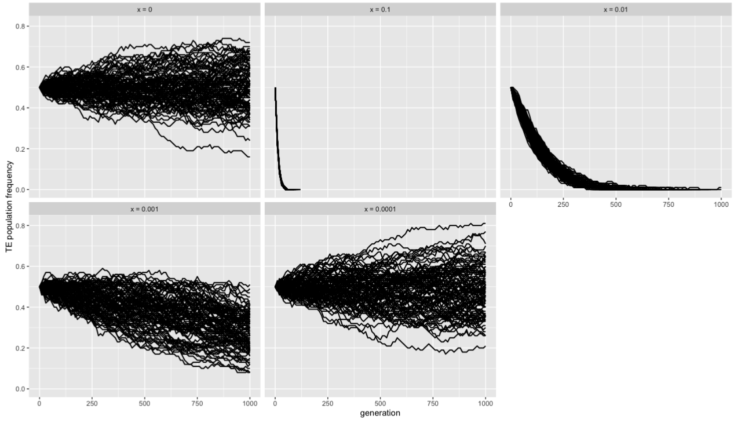

Figure 17: Validation of selection effects: Progression of TE population (y-axis) through generations (x-axis) for different selection coefficients (x) on the same population. Top left: x = 0, top middle: x = 0.1, top left: x = 0.01, bottom right: x = 0.001, bottom left: x = 0.0001.

Figure 17 presents a facet wrap plot showing the progression of TE population frequency (y-axis) through generations (x-axis) for different selection coefficients (x) on the same population. In the top left panel (x = 0), the base population is defined as 4000 TEs in the genome, and the replicates show both upward and downward trends, indicating that the TE population frequency is not biased. Moving towards the right, in the top middle panel (x = 0.1), all replicates show a decrease in TE population frequency towards 0. In the top left panel (x = 0.01), more replicates survive, but the TE count still decreases. The bottom right panel (x = 0.001) shows a similar trend, with more surviving replicates but a consistent decrease in TE frequency. The bottom left panel (x = 0.0001) resembles the top left panel, with TE frequency moving both up and down.

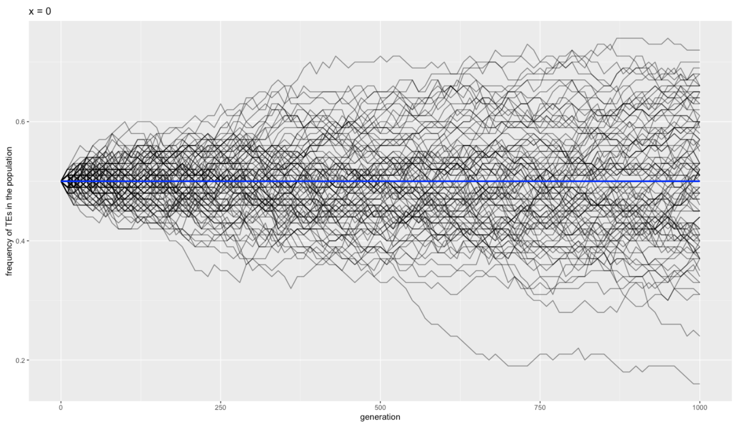

Figure 18: Validation of selection effects: Progression of TE population frequency through generations for x = 0. The blue line represents the average TE frequency.

Figure 18 focuses on the progression of TE population frequency for x = 0, with the blue line indicating the average TE frequency.

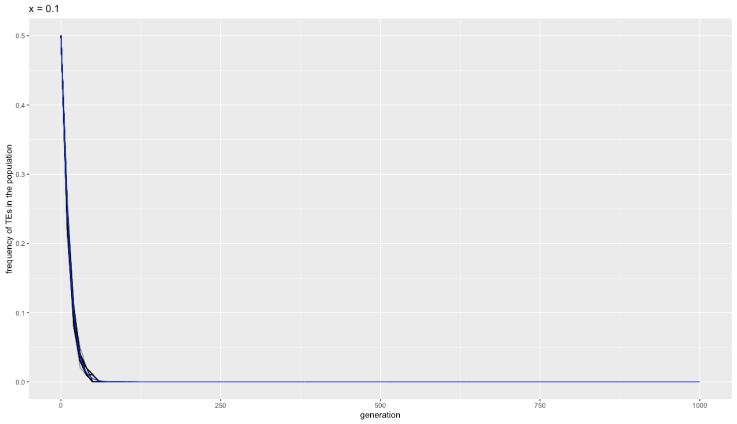

Figure 19: Validation of selection effects: Progression of TE population frequency through generations for x = 0.1. The blue line represents the average TE frequency.

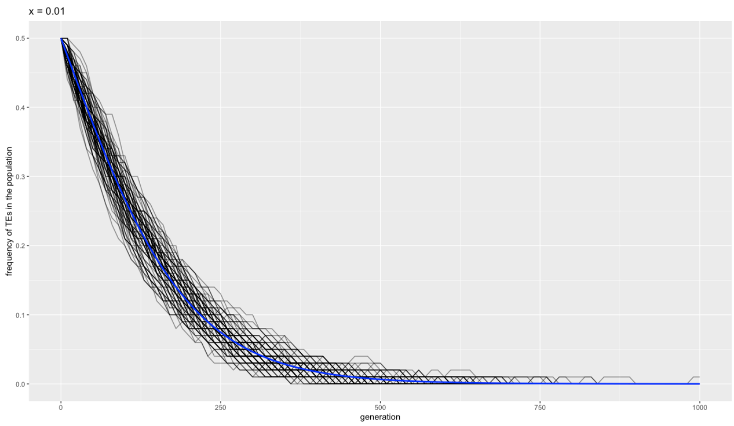

Figure 20: Validation of selection effects: Progression of TE population frequency through generations for x = 0.01. The blue line represents the average TE frequency.

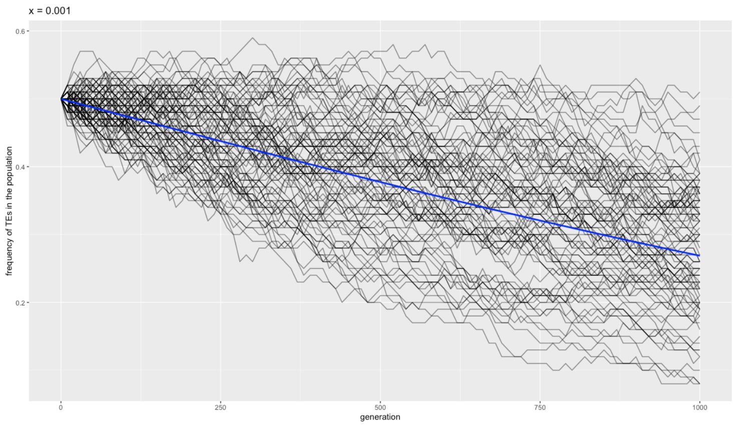

Figure 21: Validation of selection effects: Progression of TE population frequency through generations for x = 0.001. The blue line represents the average TE frequency.

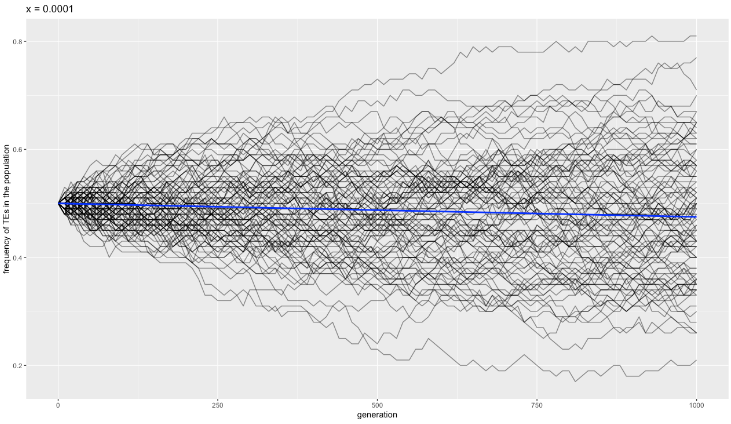

Figure 22: Validation of selection effects: Progression of TE population frequency through generations for x = 0.0001. The blue line represents the average TE frequency.

Figures 19, 20, 21, and 22 show the progression of TE population frequency through generations for selection coefficients x = 0.1, x = 0.01, x = 0.001, and x = 0.0001, respectively.

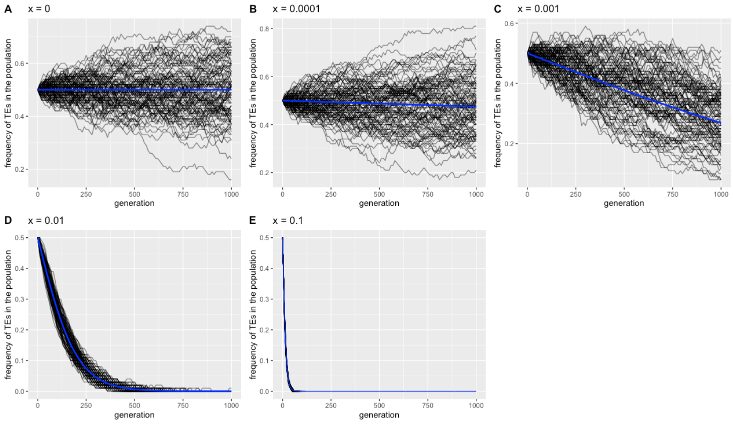

Figure 23: Validation of selection effects: Progression of TE population frequency through generations for different selection coefficients (x) on the same population, with average TE frequency represented by blue lines. Top left: x = 0, top right: x = 0.1, middle left: x = 0.01, middle right: x = 0.001, bottom: x = 0.0001.

Figure 23 is similar to Figure 17 but includes blue lines representing the average TE frequency for each selection coefficient.

#### Insertion Validation

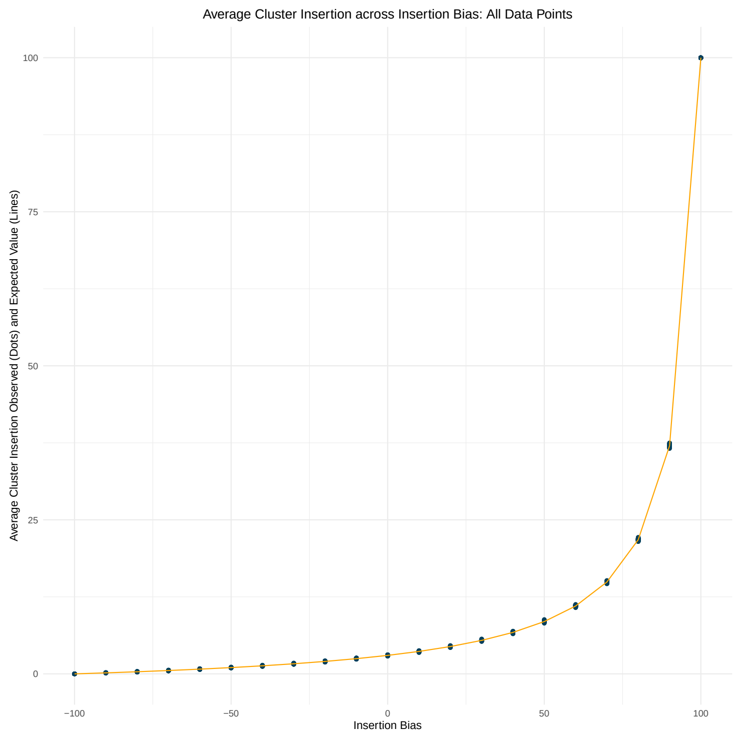

Figure 24: Validation of insertion: Average cluster insertion as a function of insertion bias. The x-axis represents the insertion bias ranging from -100 to 100 with a step size of 10. The y-axis shows the average cluster insertion. The dots in the plot represent the observed values, while the orange line represents the expected values. The observed and expected values match, showing the steep exponential curve.

This plot validates our implementation of insertion as piRNA cluster insertion is translating as expected with insertion bias.

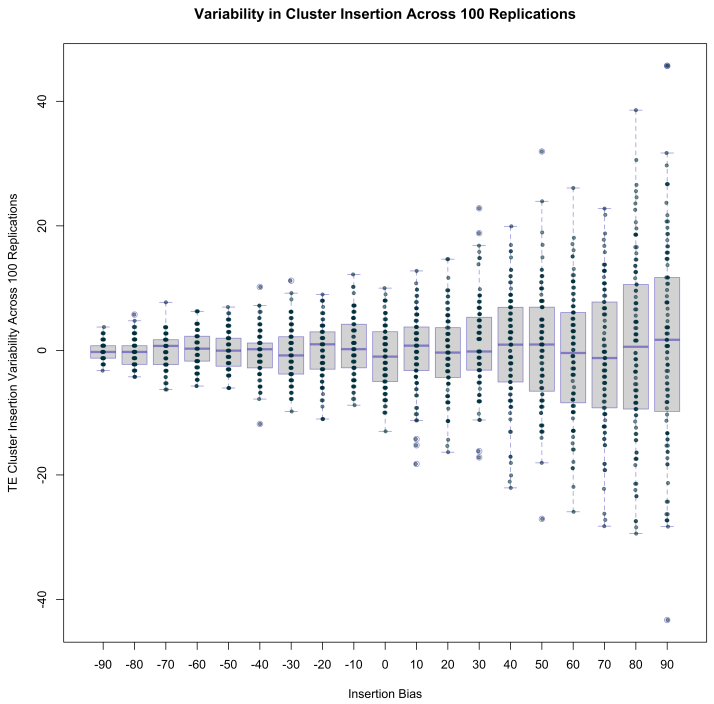

Figure 25: Validation of insertion: Variability in TE insertions across different levels of insertion bias, based on data from 100 replications. The y-axis represents the relative difference between 'avcli' and 'pc' columns, expressed as a percentage. The x-axis represents the categorical 'Insertion Bias' variable. Each boxplot summarizes the distribution of 'TE_insertions' for each level of 'Insertion Bias', indicating the median, interquartile range, and potential outliers.

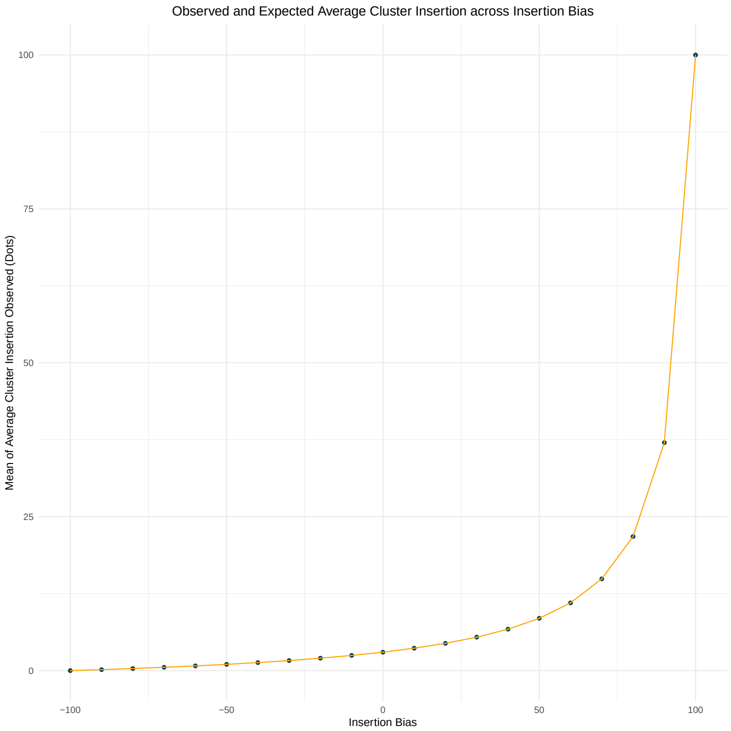

Figure 26: Validation of insertion: Mean of average cluster insertion across various insertion bias levels, with error bars indicating standard deviation. The orange line represents the expected values.

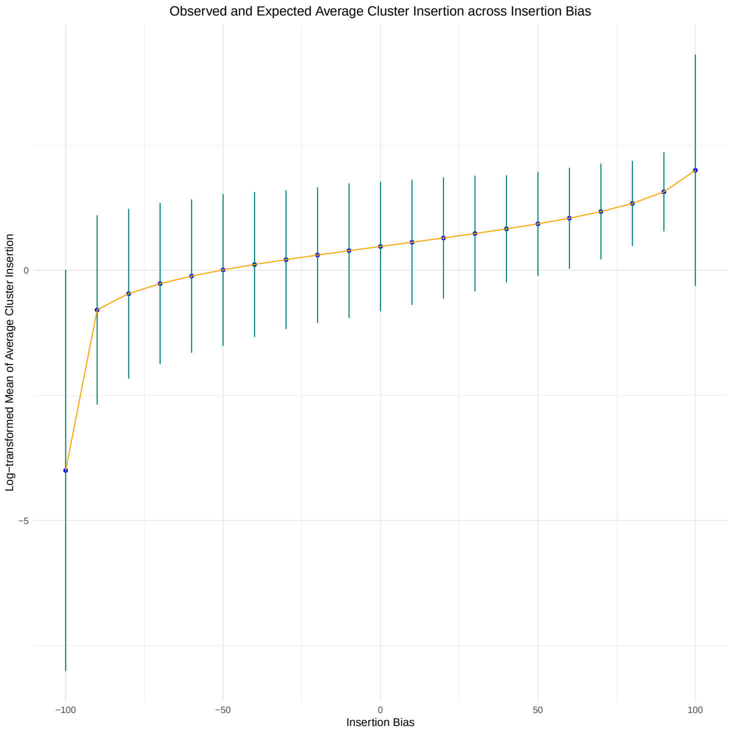

Figure 27: Validation of insertion: The same data as Figure 26, but with a logarithmic transformation applied to the y-values. The log scale helps to better visualize the variability in the data (shown by the error bars) and the deviation of the observed values (blue points) from the expected values (orange line).

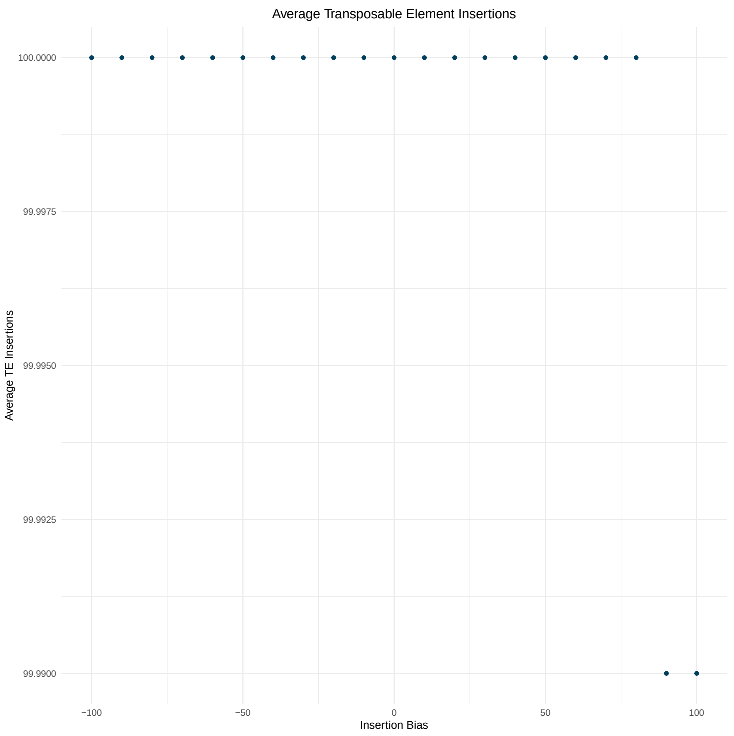

Figure 28: Validation of insertion: Average TE insertions (y-axis) and insertion bias (x-axis) for a single replicate. The x-axis represents the insertion bias varying from -100 to 100, while the y-axis depicts the average TE insertions for each bias level. Each data point represents the average TE insertions for a specific bias level for replicate 1.

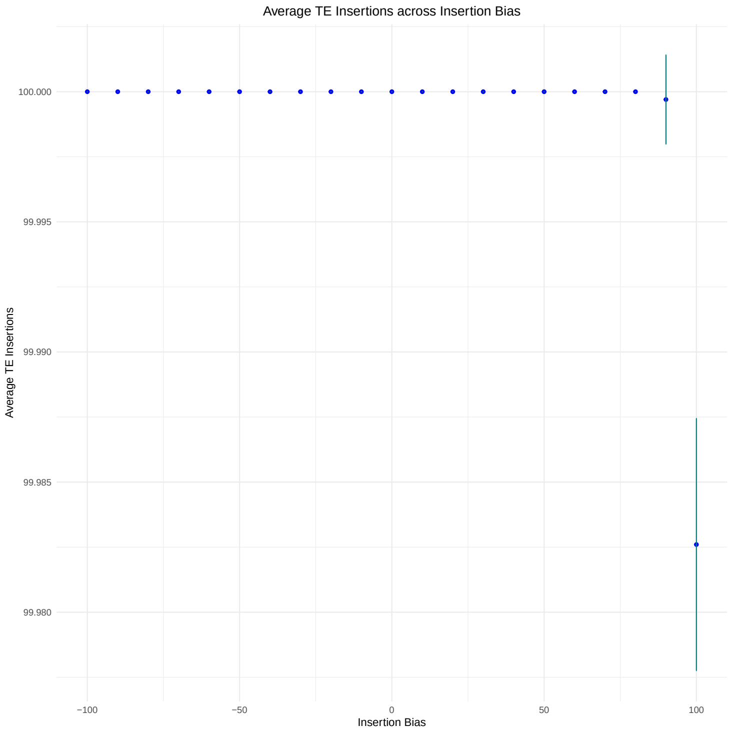

Figure 29: Validation of insertion: All replicates' mean average TE insertions (y-axis) against the insertion bias (x-axis). The plot includes error bars representing the standard deviation for the average TE insertions at each bias level. The x-axis represents the insertion bias, ranging from -100 to 100, while the y-axis depicts the mean of the average TE insertions for each bias level.

Across the 21 insertion bias values (sampleid 0 to 20), the mean of cli is calculated for all 100 replications, along with the standard deviation. The pc column represents the theoretical value of average cluster insertion for a given insertion bias with a 3% cluster size. The validation matches our expectations, confirming that the insertion is working as expected and the simulation successfully incorporates the user-defined TE insertions as specified.

### 2 Supplementary Tables

This section presents the supplementary tables related to the validation steps described in the manuscript.

Table 1: Across the 21 insertion bias values (sampleid 0 to 20), the mean of cluster insertion is calculated for all 100 replications. The standard deviation is also calculated. We have another column named pc which is the theoretical value of average cluster insertion for a given insertion bias with a 3% cluster size.

| sampleid | mean_cli | sd_meancli | pc | deviation_pc |
|---:|---:|---:|---:|---:|
| -100 | 0.000 | 0.000 | 0.000 | 0.000 |
| -90 | 0.162 | 0.0130 | 0.163 | -0.000814 |
| -80 | 0.341 | 0.0204 | 0.342 | -0.00157 |
| -70 | 0.542 | 0.0248 | 0.543 | -0.000423 |
| -60 | 0.769 | 0.0297 | 0.767 | 0.00154 |
| -50 | 1.020 | 0.0307 | 1.020 | -0.00111 |
| -40 | 1.300 | 0.0360 | 1.310 | -0.00454 |
| -30 | 1.630 | 0.0417 | 1.640 | -0.00397 |
| -20 | 2.020 | 0.0448 | 2.020 | -0.00160 |
| -10 | 2.480 | 0.0459 | 2.470 | 0.00719 |
| 0 | 2.990 | 0.0514 | 3.000 | -0.00930 |
| 10 | 3.640 | 0.0571 | 3.640 | 0.000316 |
| 20 | 4.430 | 0.0622 | 4.430 | -0.00360 |
| 30 | 5.440 | 0.0707 | 5.430 | 0.00955 |
| 40 | 6.730 | 0.0861 | 6.730 | 0.00333 |
| 50 | 8.500 | 0.0932 | 8.490 | 0.00513 |
| 60 | 11.000 | 0.0994 | 11.000 | -0.00867 |
| 70 | 14.900 | 0.112 | 14.900 | -0.0104 |
| 80 | 21.800 | 0.144 | 21.800 | 0.00271 |
| 90 | 37.000 | 0.164 | 37.000 | 0.0114 |
| 100 | 100.000 | 0.00485 | 100.000 | -0.0174 |

Table 2: Probability (p) that a TE jumps into a piRNA cluster depending on the insertion bias of the TE. The piRNA clusters accounts for 3% of the genome. Additionally, we show the number of expected insertions and observed with our software invade-insbias for 1000 new TE insertions. Observed values are averaged over 1000 replicates. This demonstrates that our software correctly implements the insertion bias.

| bias | p | expected | observed |
|---|---|---|---|
| 100 | 1.000 | 1000 | 1000 |
| 90 | 0.370 | 370 | 369.72 |
| 80 | 0.218 | 218 | 217.65 |
| 70 | 0.149 | 149 | 149.66 |
| 60 | 0.110 | 110 | 109.57 |
| 50 | 0.085 | 85 | 84.91 |
| 40 | 0.067 | 67 | 67.16 |
| 30 | 0.054 | 54 | 54.42 |
| 20 | 0.044 | 44 | 44.36 |
| 10 | 0.036 | 36 | 36.68 |
| 0 | 0.030 | 30 | 30.11 |
| -10 | 0.025 | 25 | 24.57 |
| -20 | 0.020 | 20 | 20.19 |
| -30 | 0.016 | 16 | 16.37 |
| -40 | 0.013 | 13 | 12.98 |
| -50 | 0.010 | 10 | 10.24 |
| -60 | 0.008 | 8 | 7.65 |
| -70 | 0.005 | 5 | 5.44 |
| -80 | 0.003 | 3 | 3.48 |
| -90 | 0.002 | 2 | 1.62 |
| -100 | 0.000 | 0 | 0 |
