## Supplementary figures and images for "The impact of insertion bias into piRNA clusters on the invasion of transposable elements"

### 2023_04_08_Validation_1a_Invasion.pdf

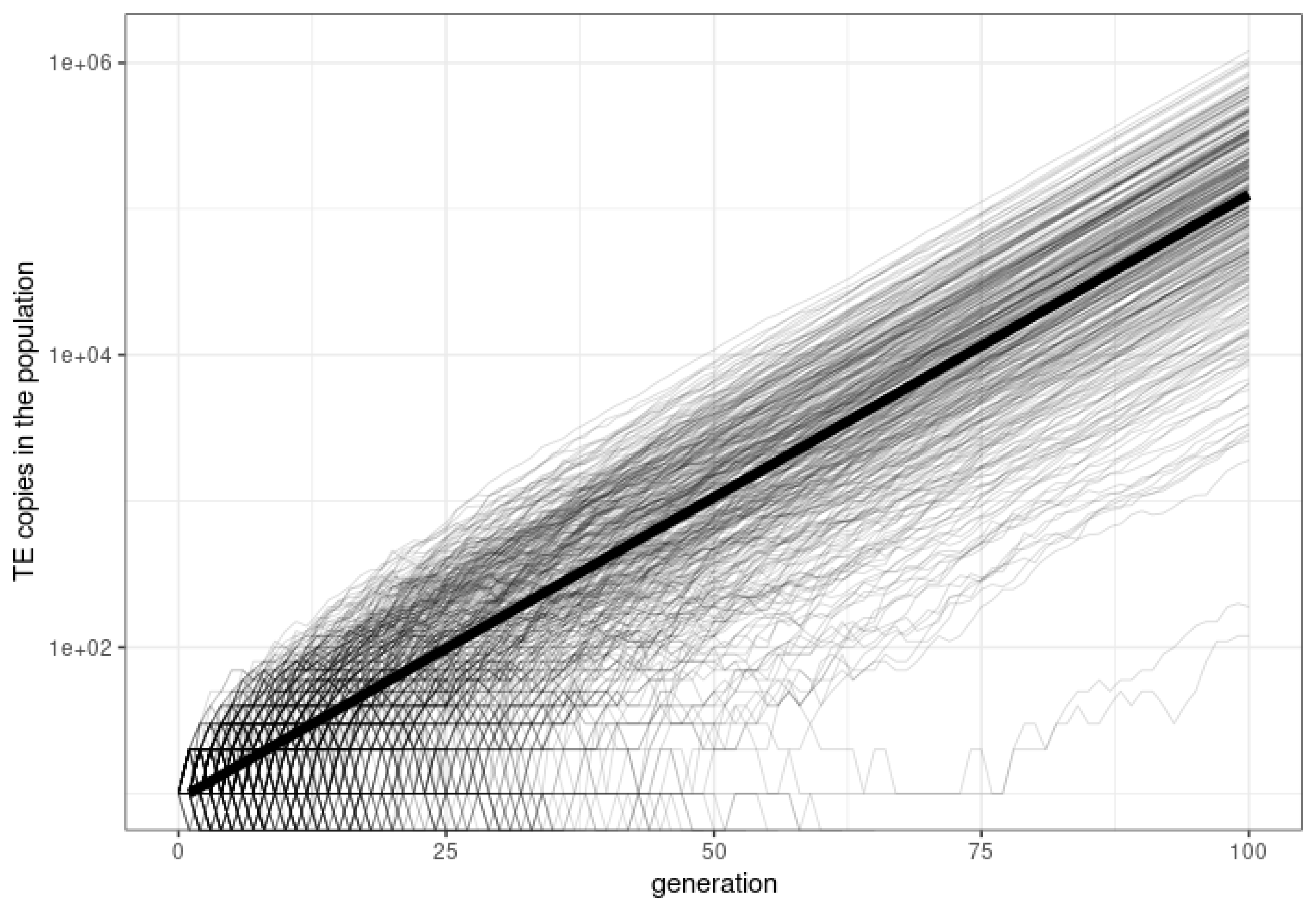

### 2023_04_08_Validation_1b_Invasion.pdf

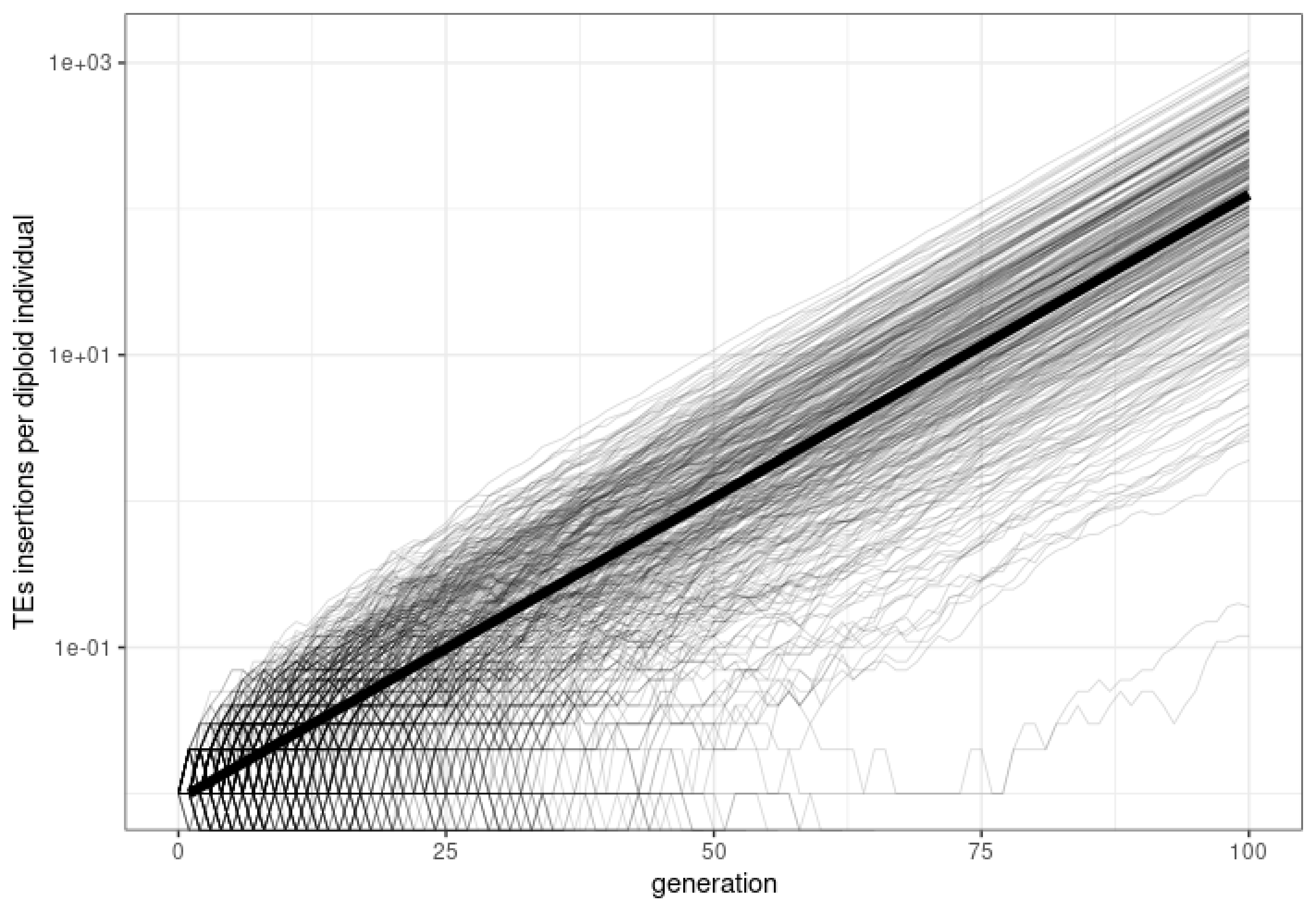

### 2023_04_11_Validation_2a_Drift.pdf

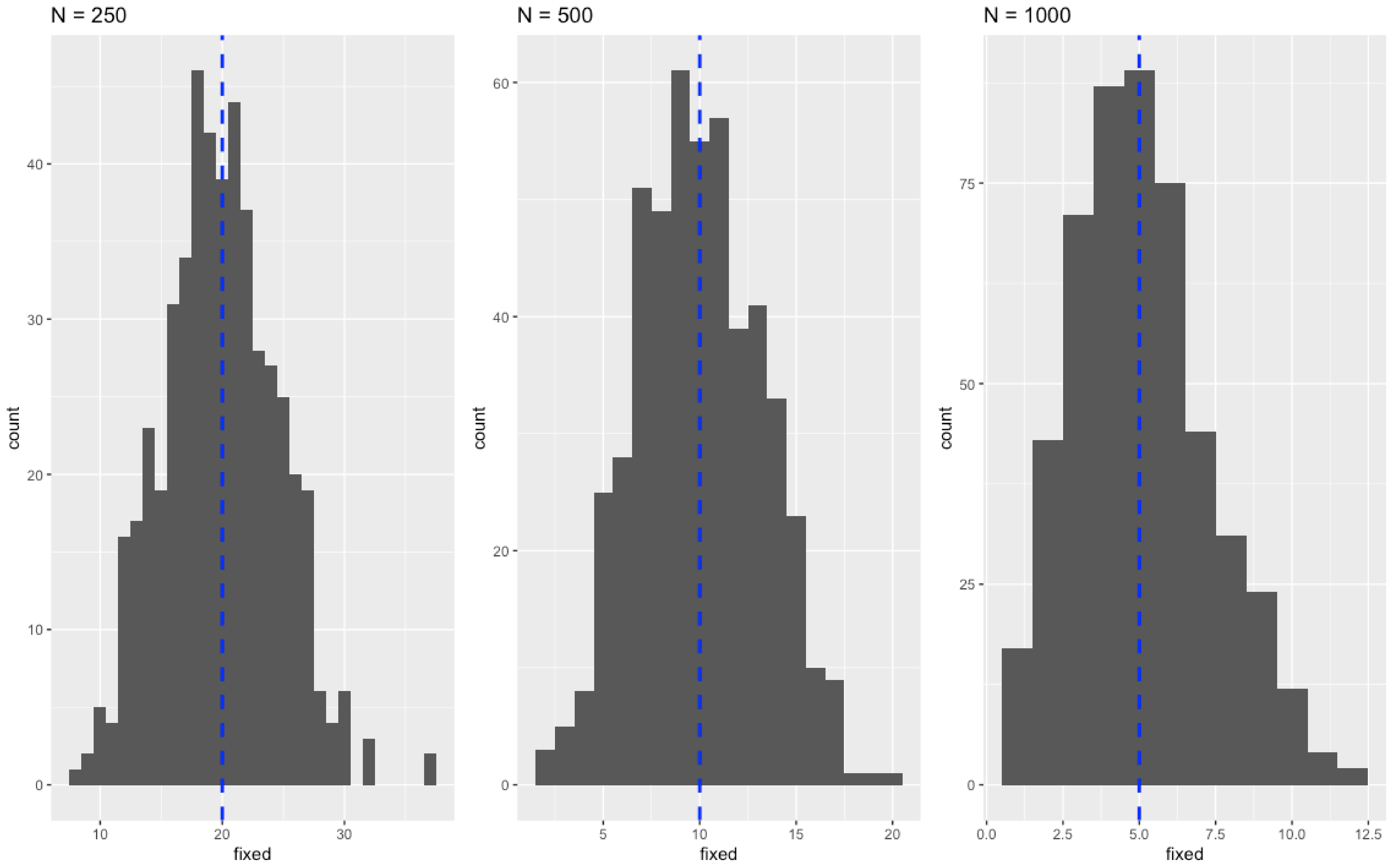

### 2023_04_11_Validation_3a_clusters.pdf

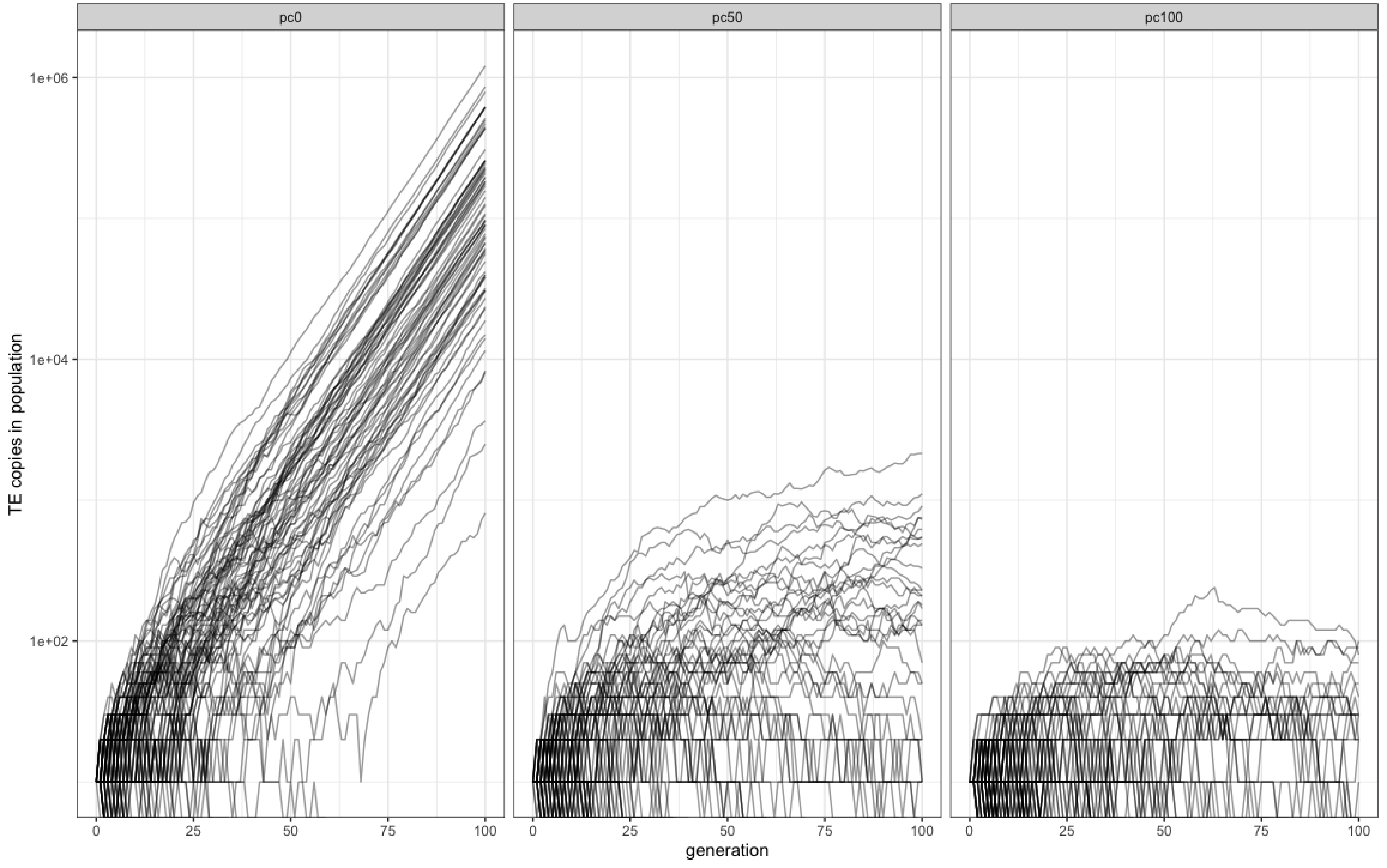

### 2023_04_11_Validation_3b_clusters.pdf

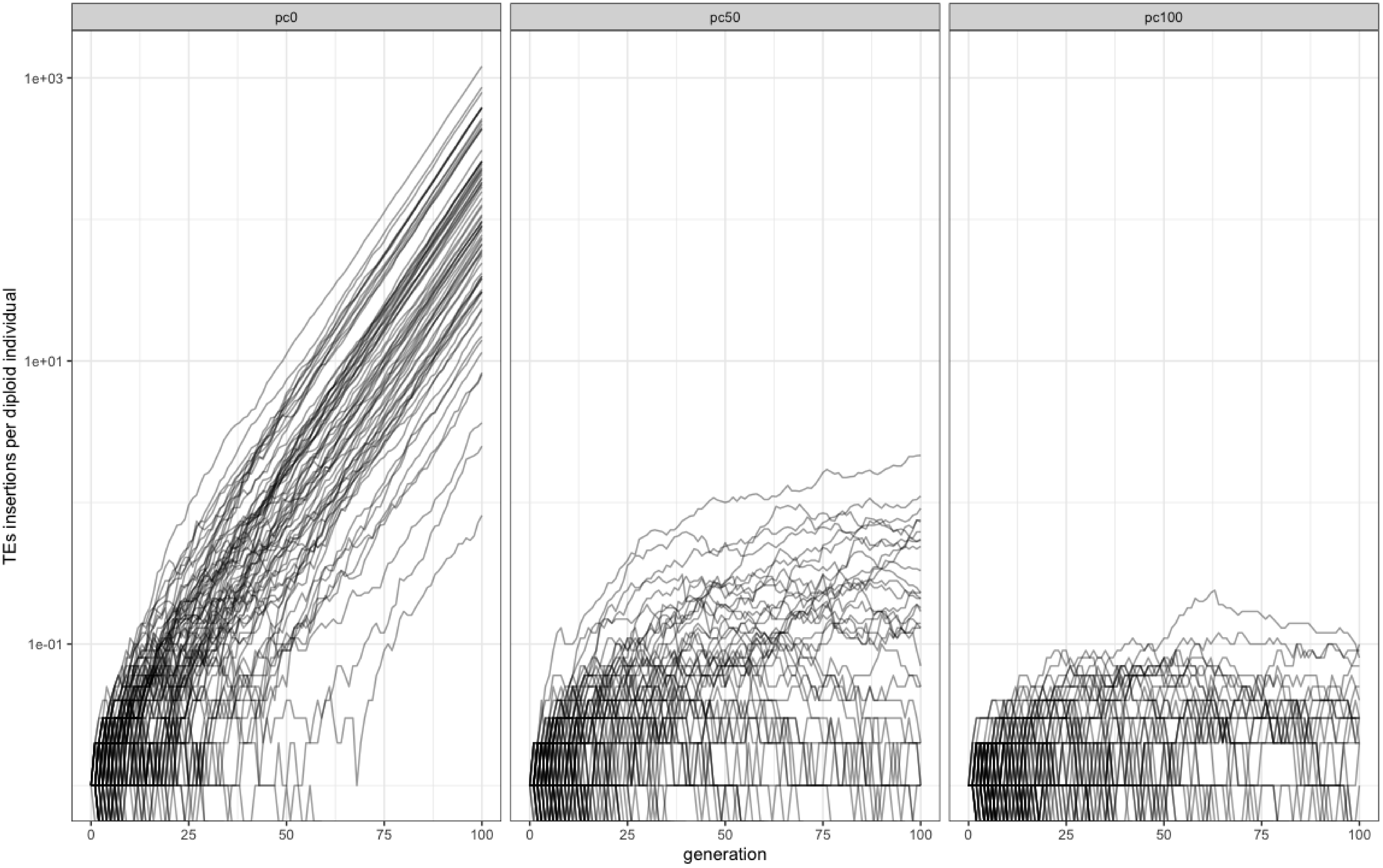

### 2023_04_17_Validation_5a_bias.pdf

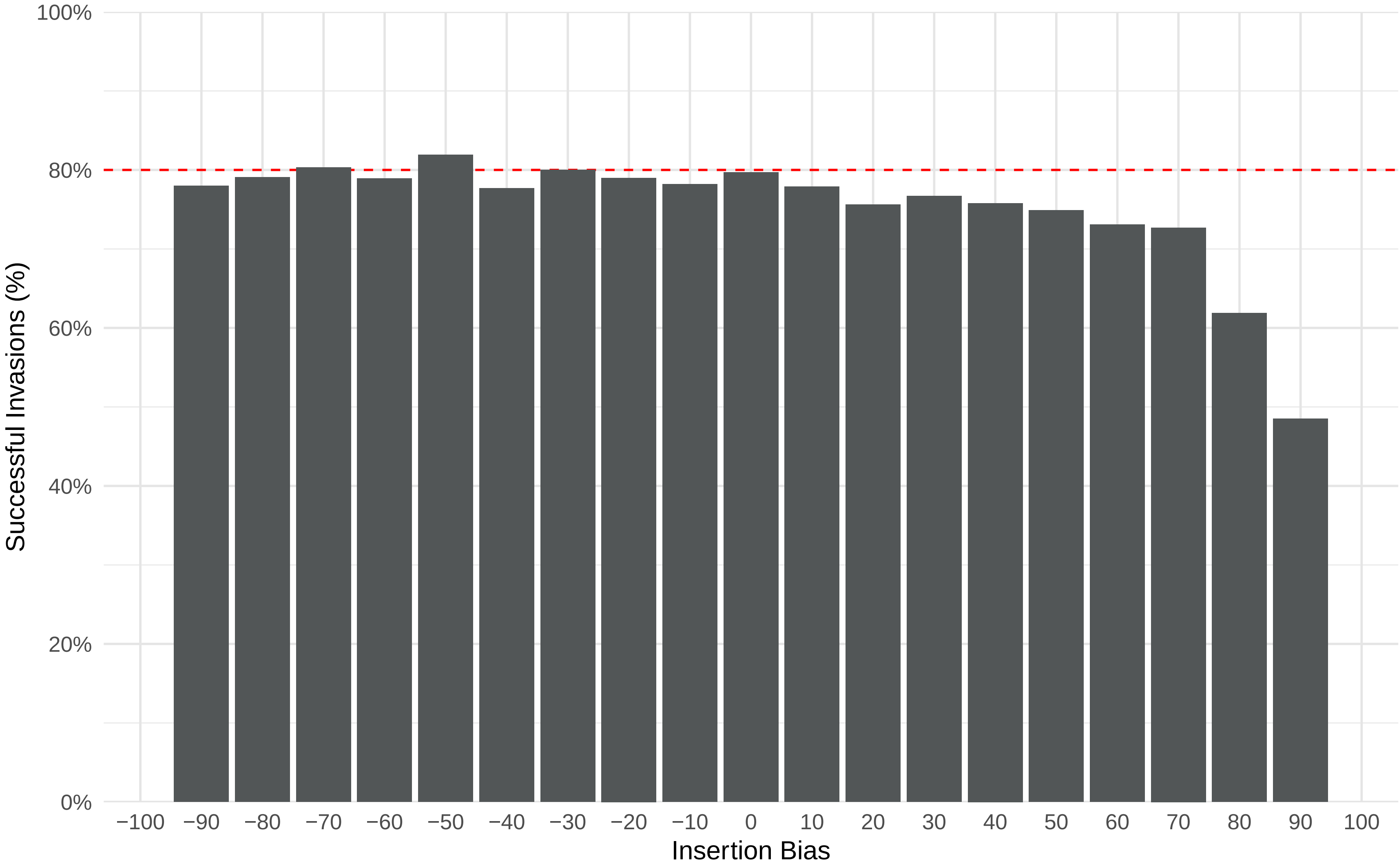

### 2023_04_18_Validation_5b_bias.pdf

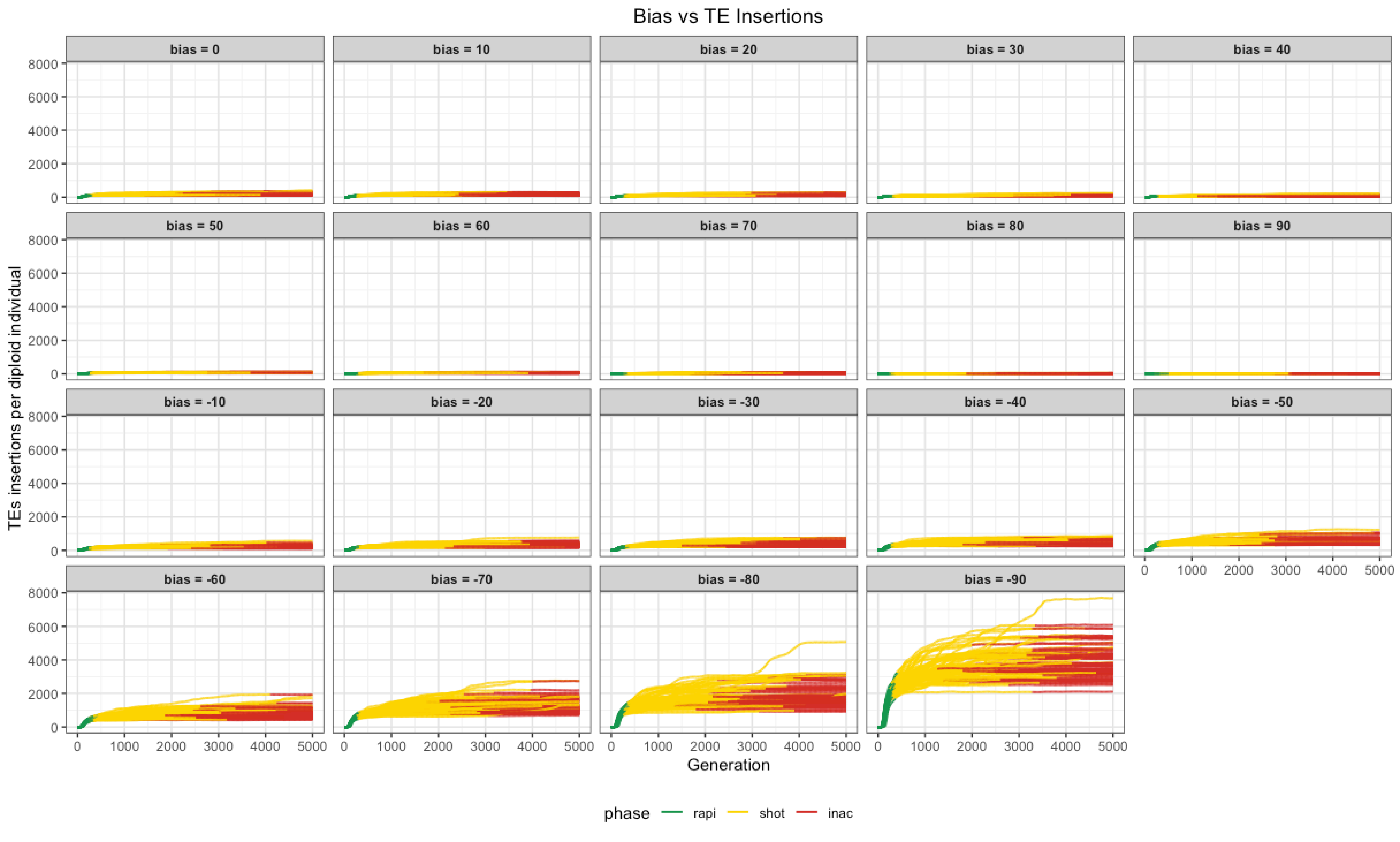

### 2023_04_18_Validation_5c_bias.pdf

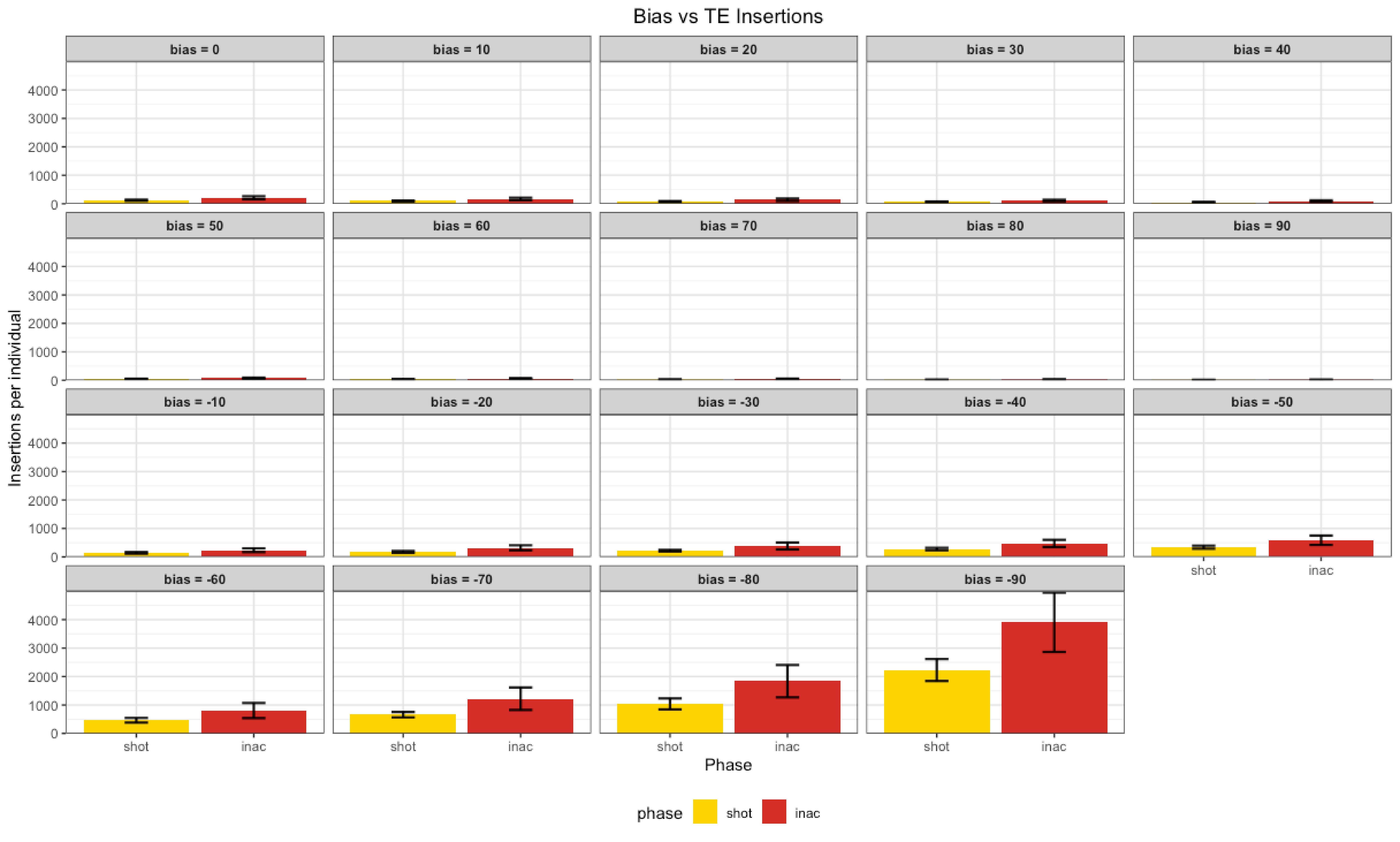

### 2023_04_18_Validation_5d_bias.pdf

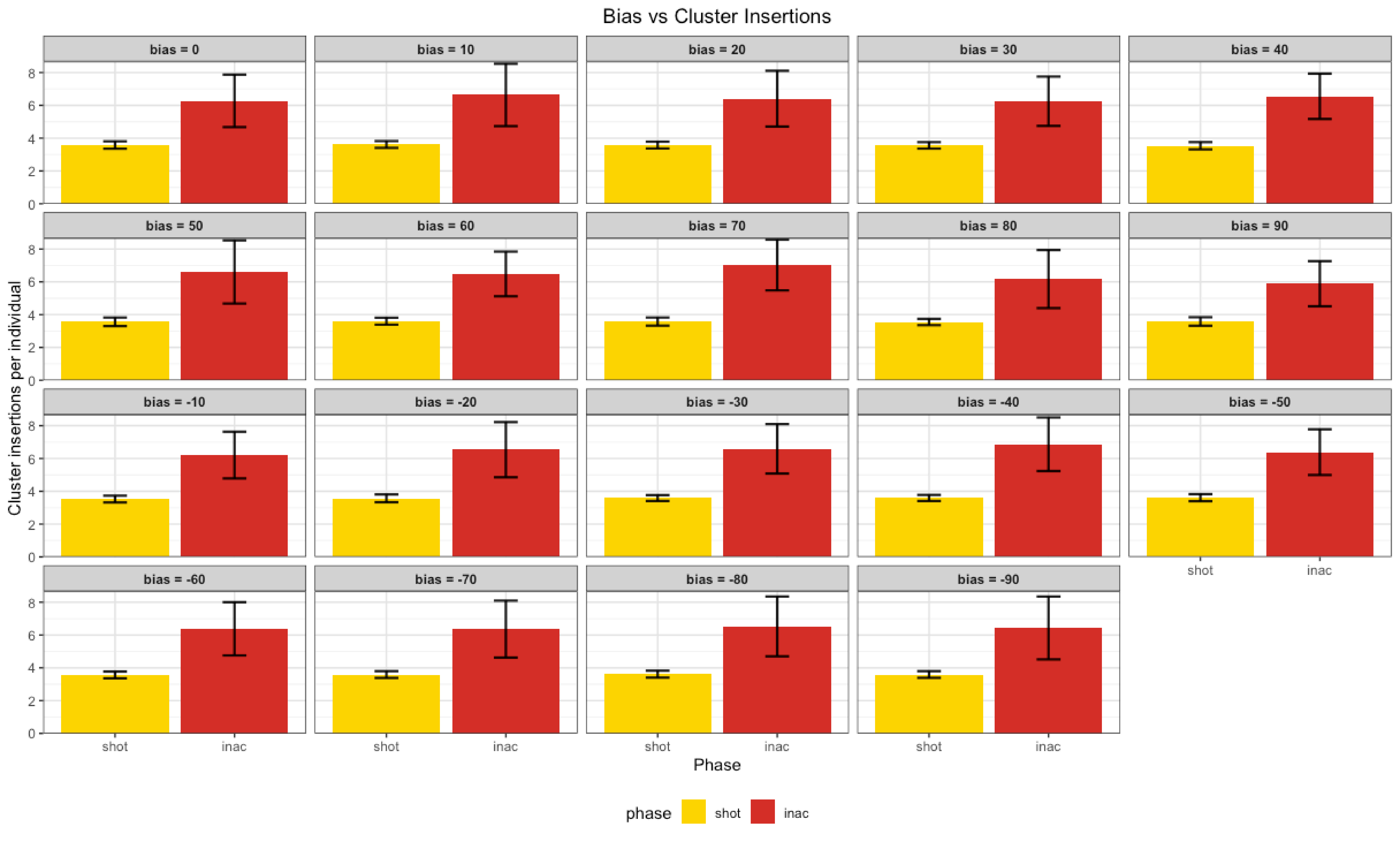

### 2023_04_18_Validation_5e_bias.pdf

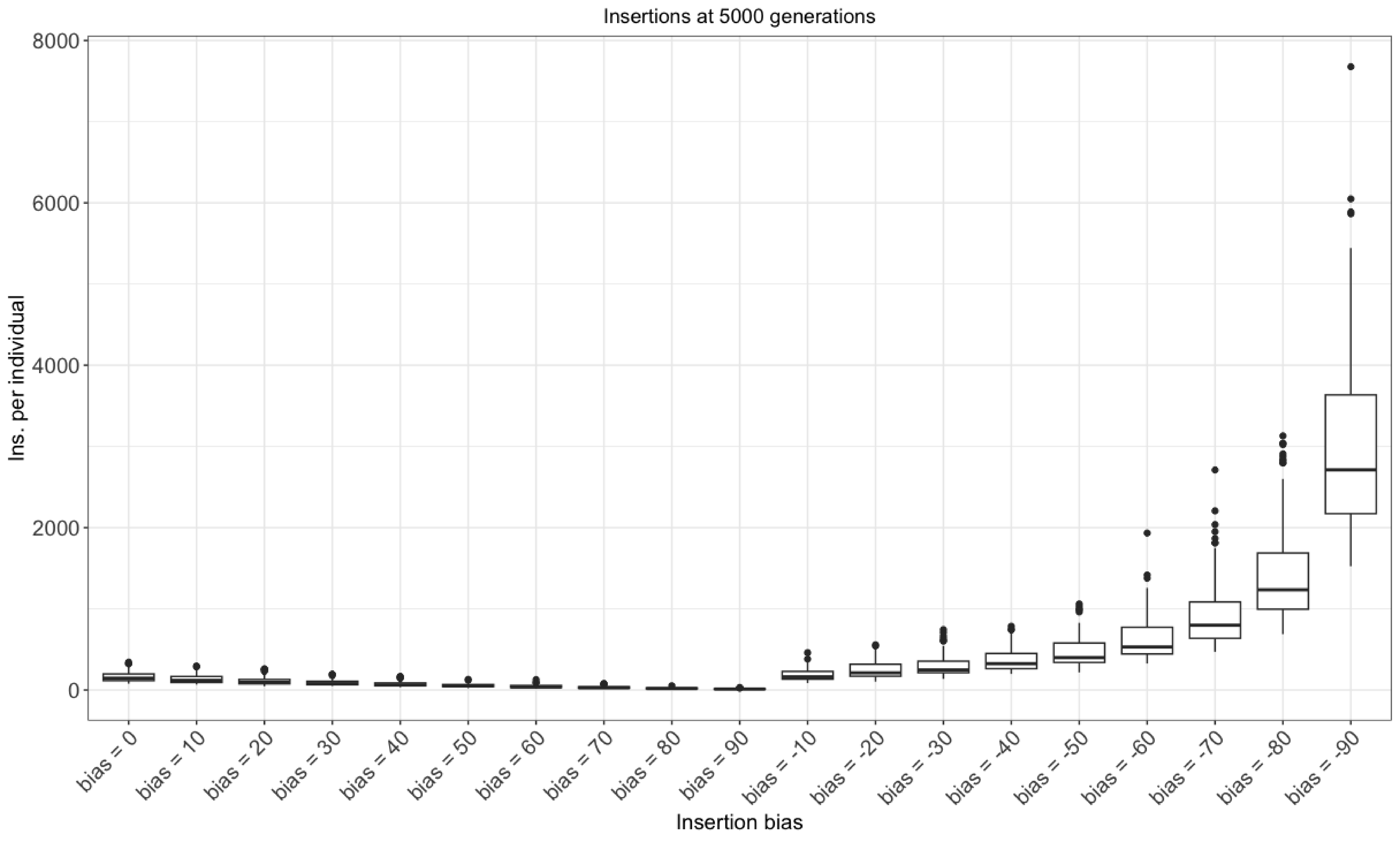

### 2023_04_18_Validation_5f_bias.pdf

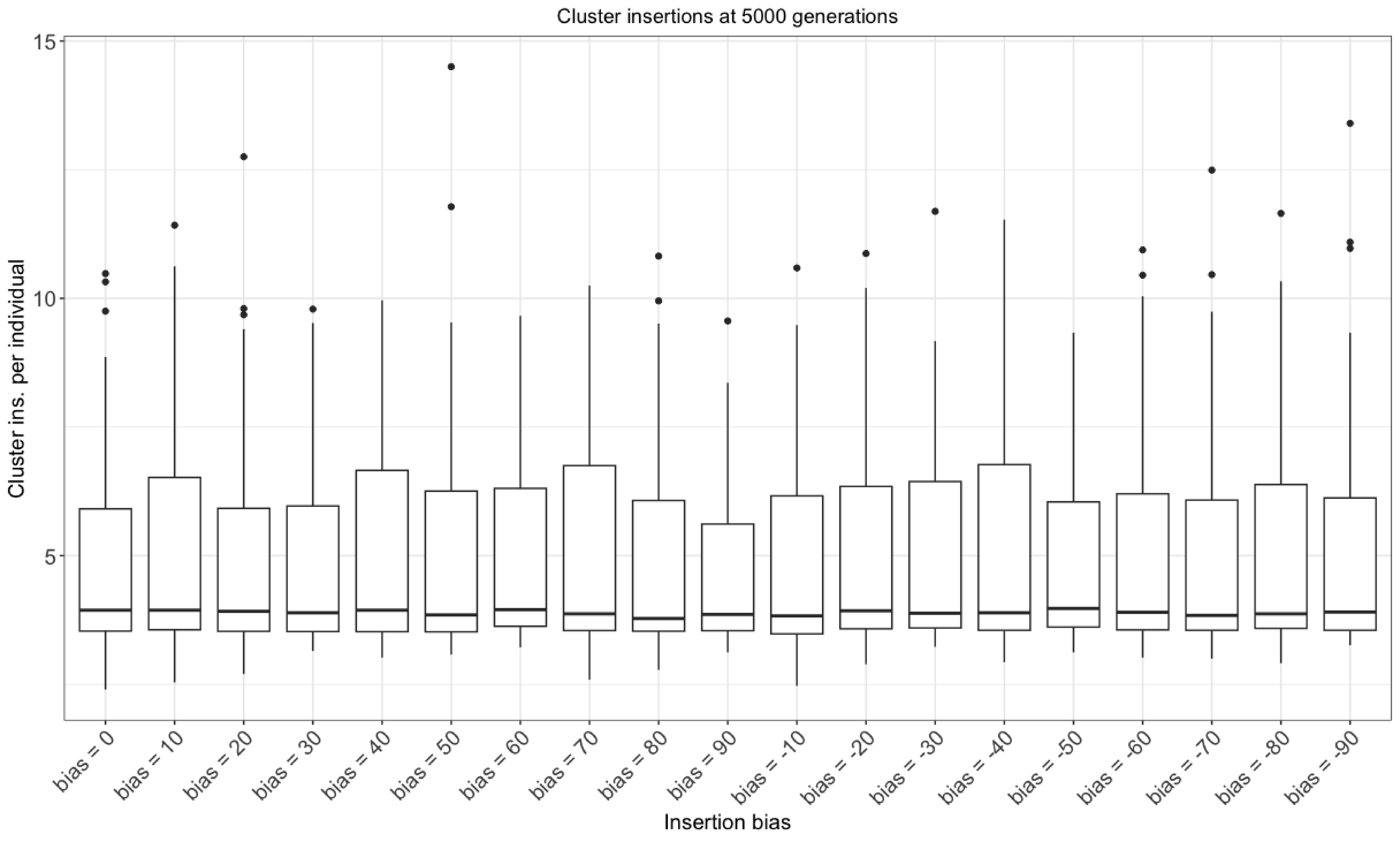

### 2023_06_05_Validation_4a_recombination.pdf

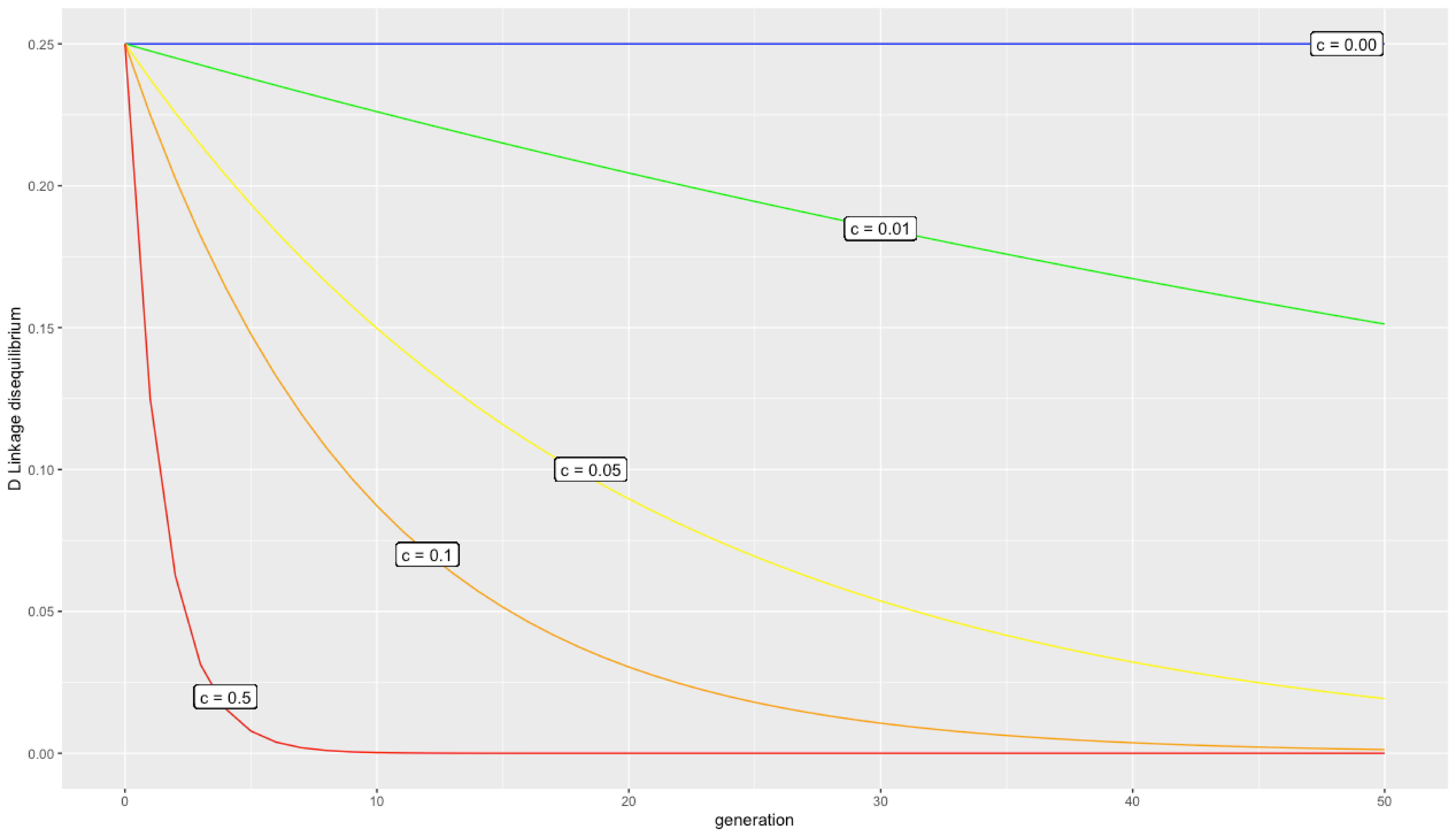

### 2023_06_05_Validation_4b_recombination.pdf

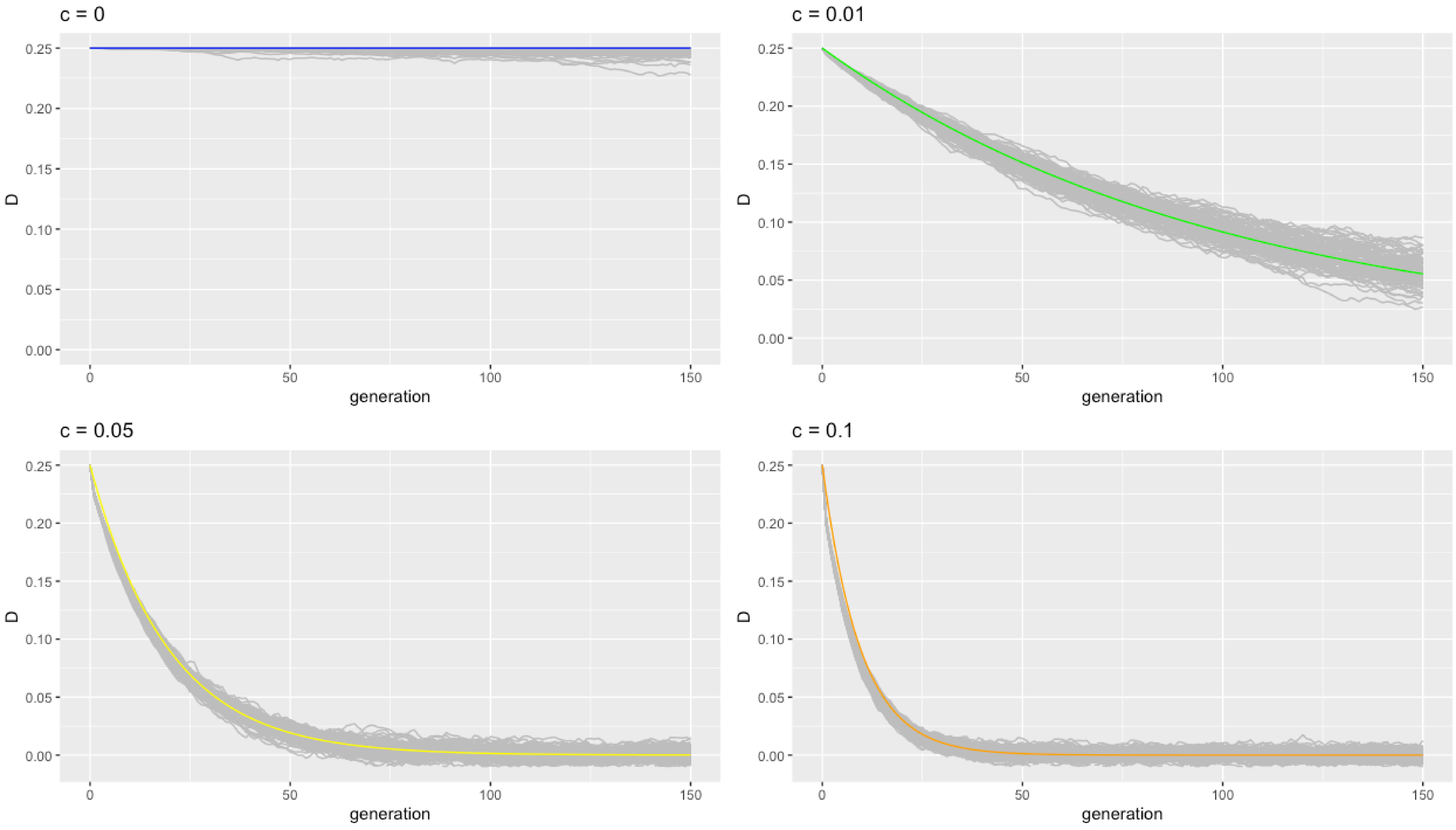

### Validation_6a_selection.pdf

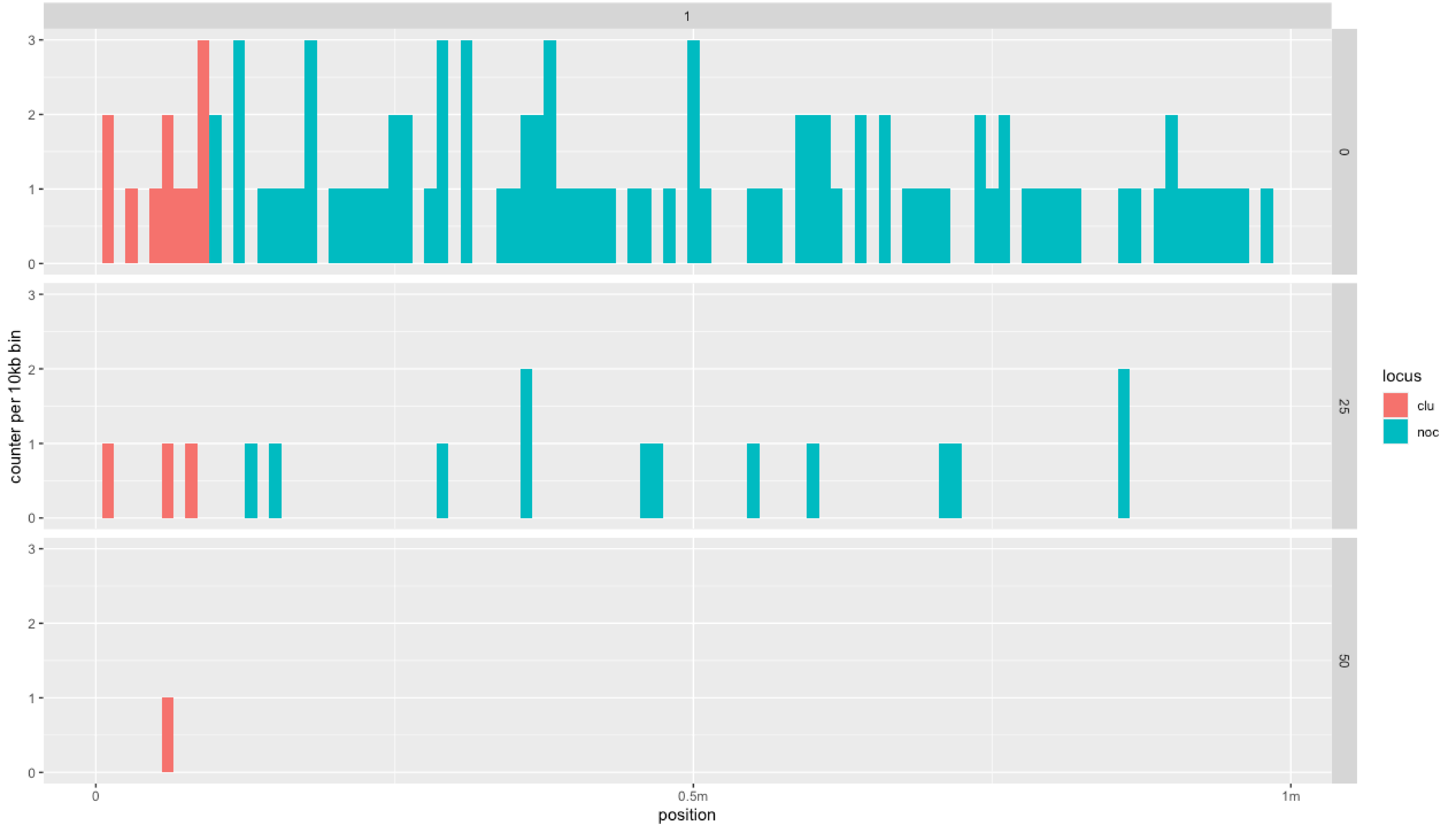

### Validation_6b_selection.pdf

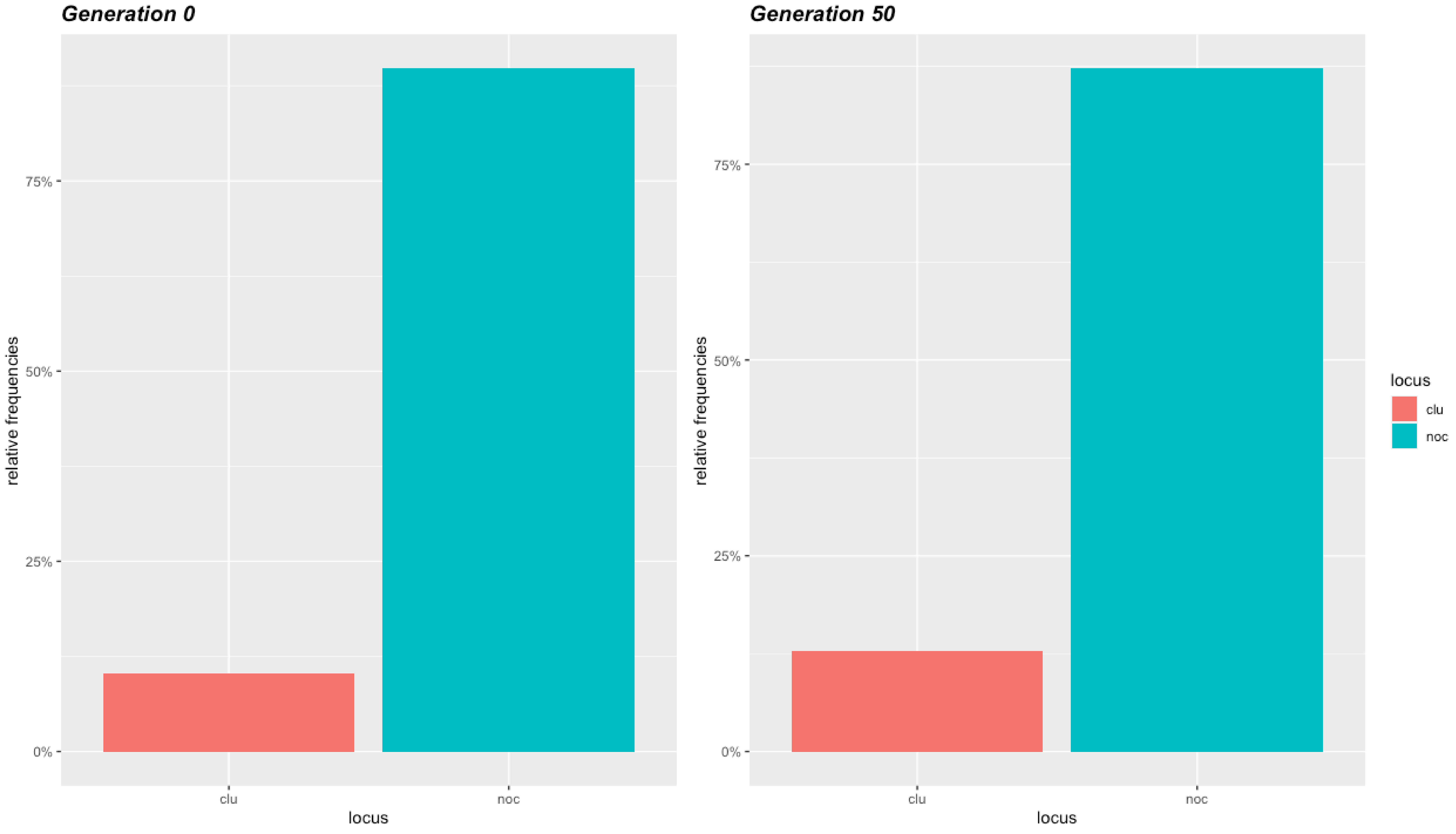

### Validation_6c_selection.pdf

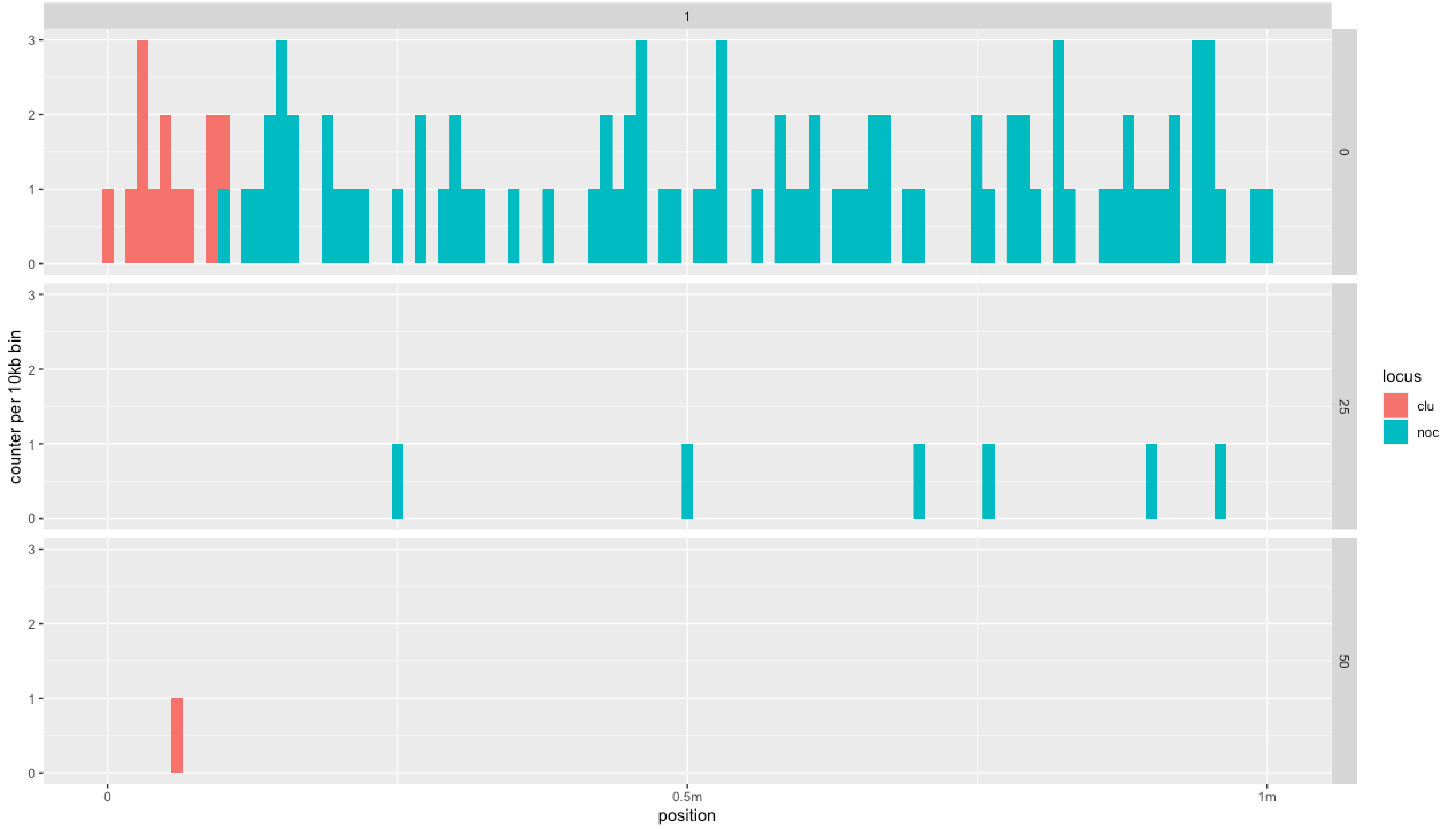

### Validation_6d_selection.pdf

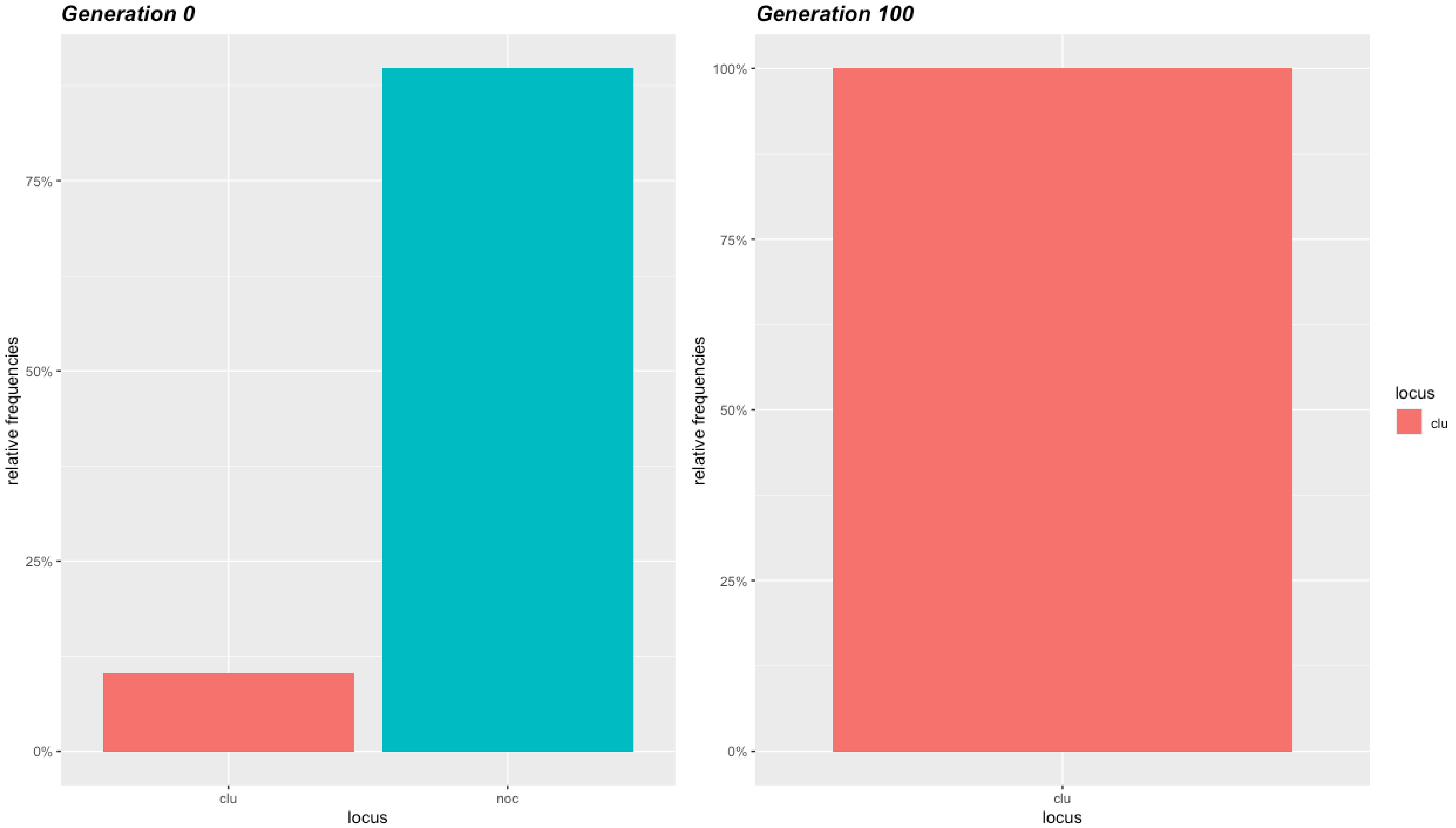

### Validation_6e_selection.pdf

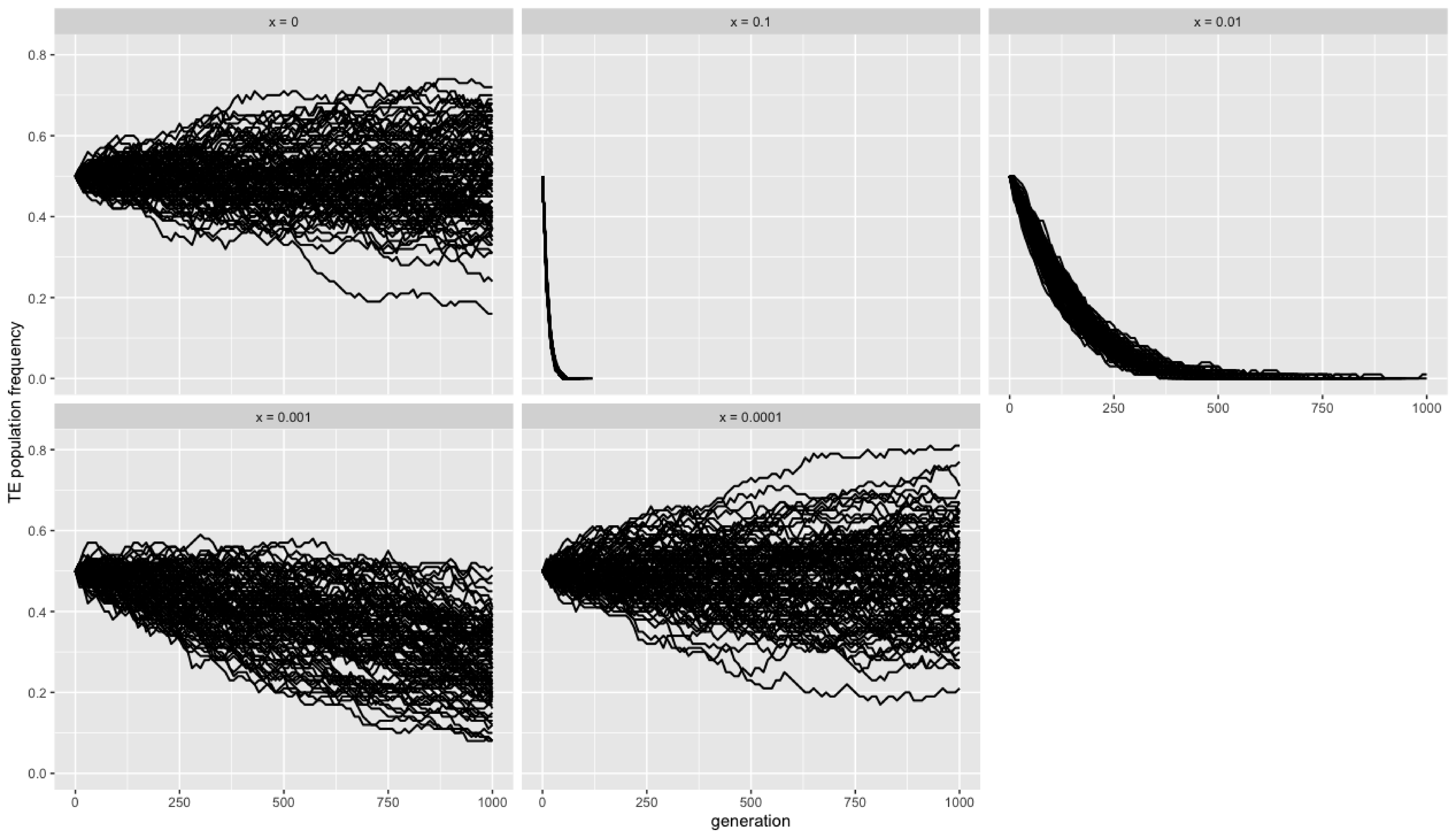

### Validation_6f_selection.pdf

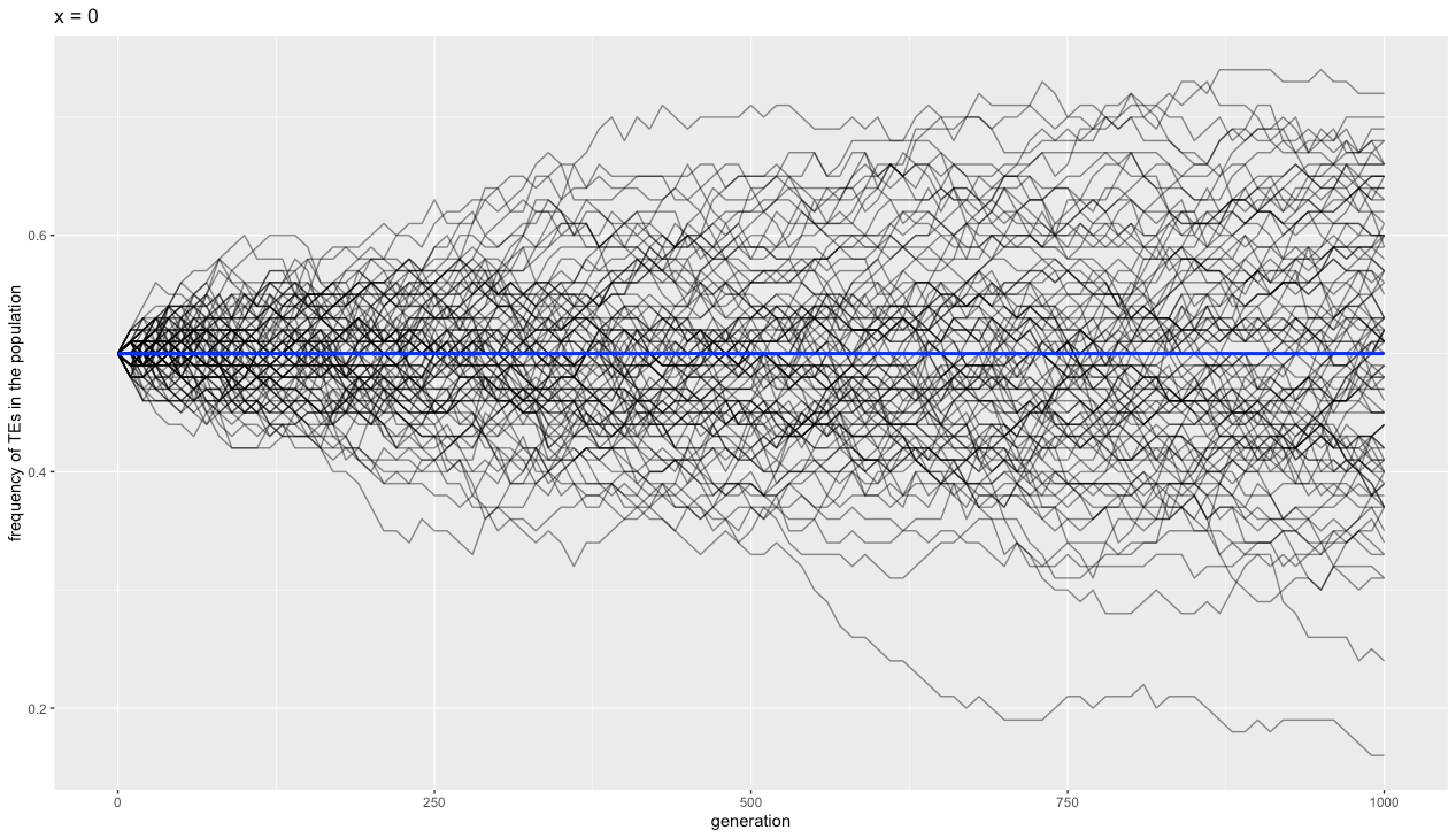

### Validation_6g_selection.pdf

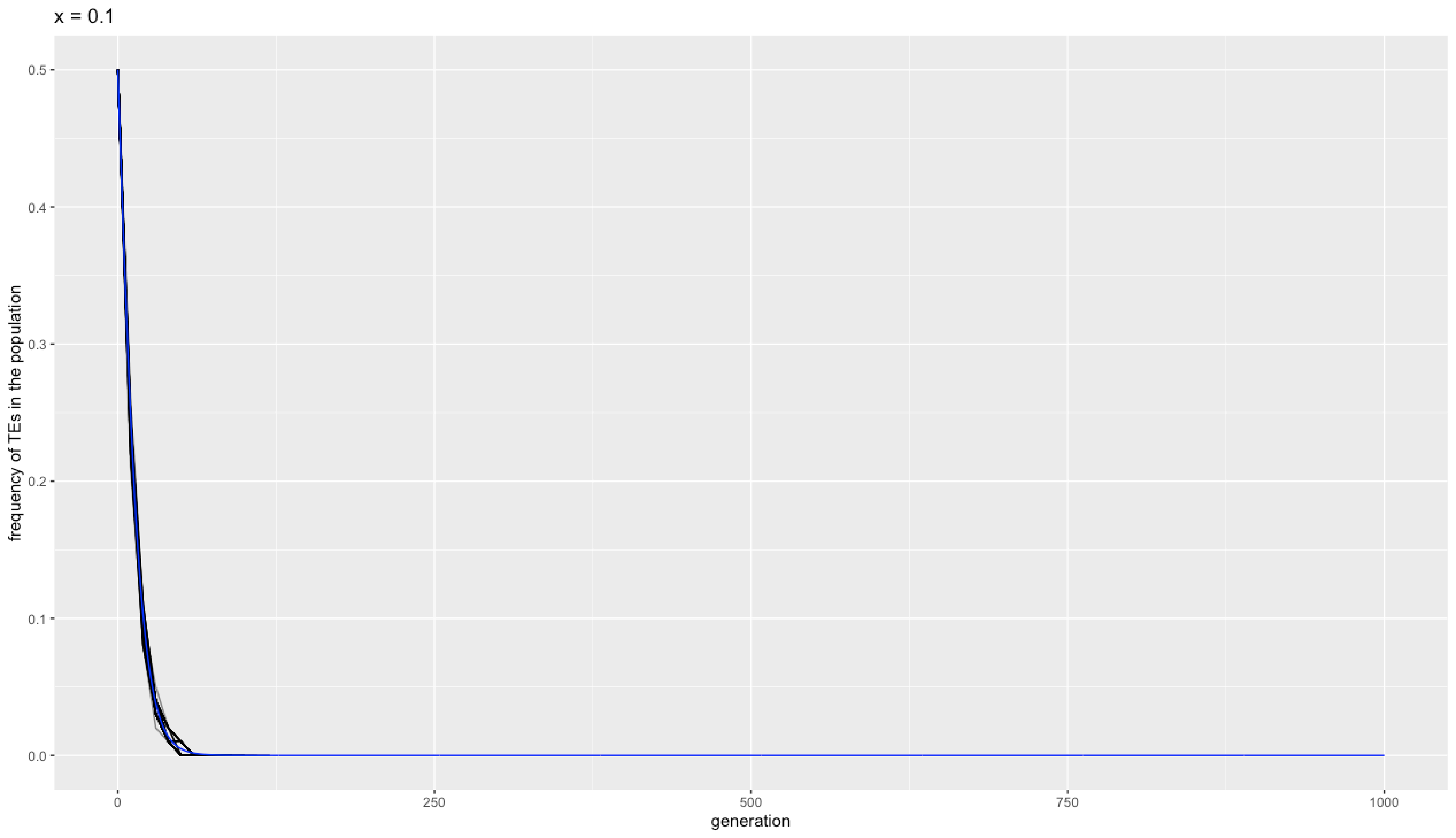

### Validation_6h_selection.pdf

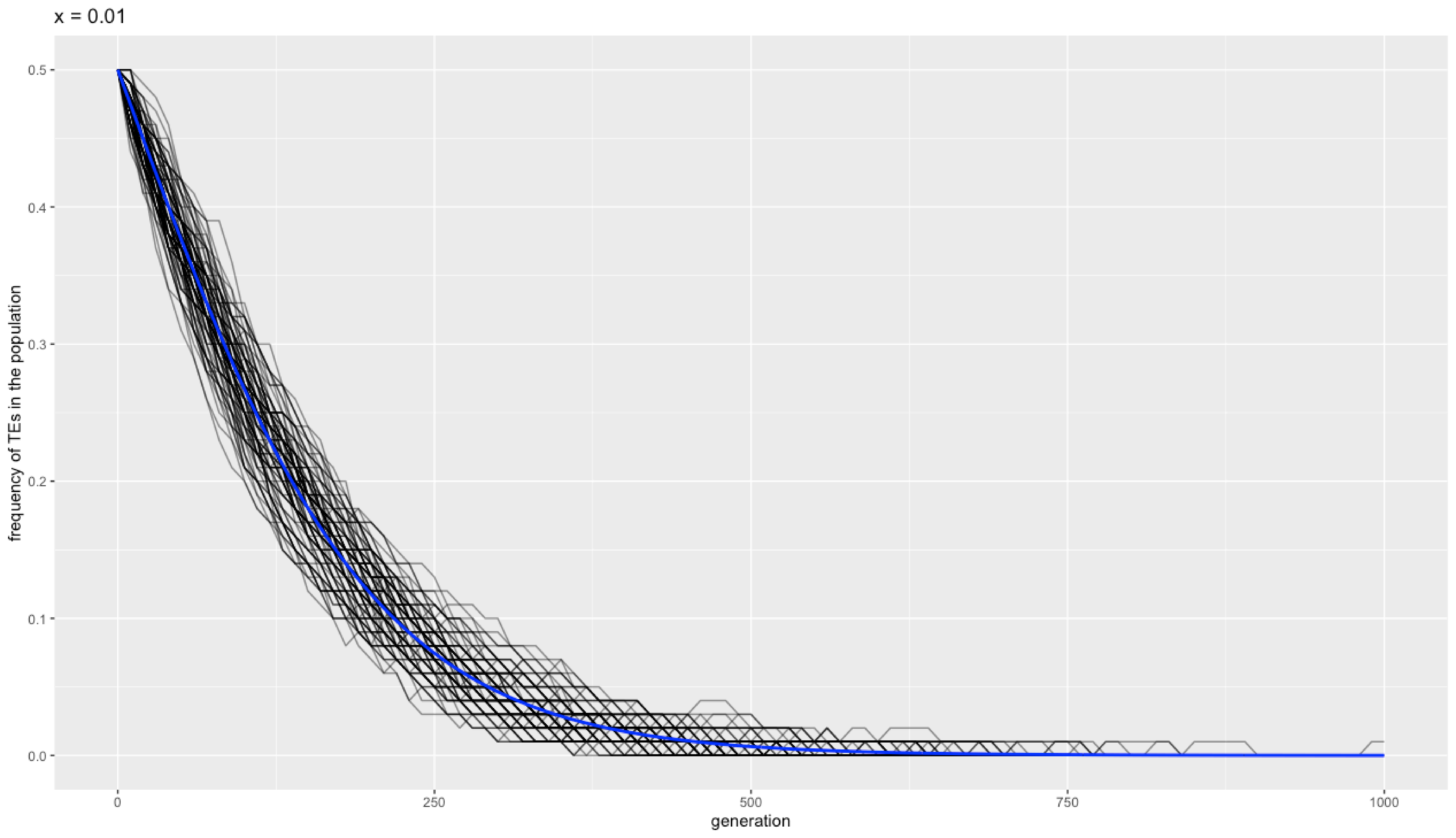

### Validation_6i_selection.pdf

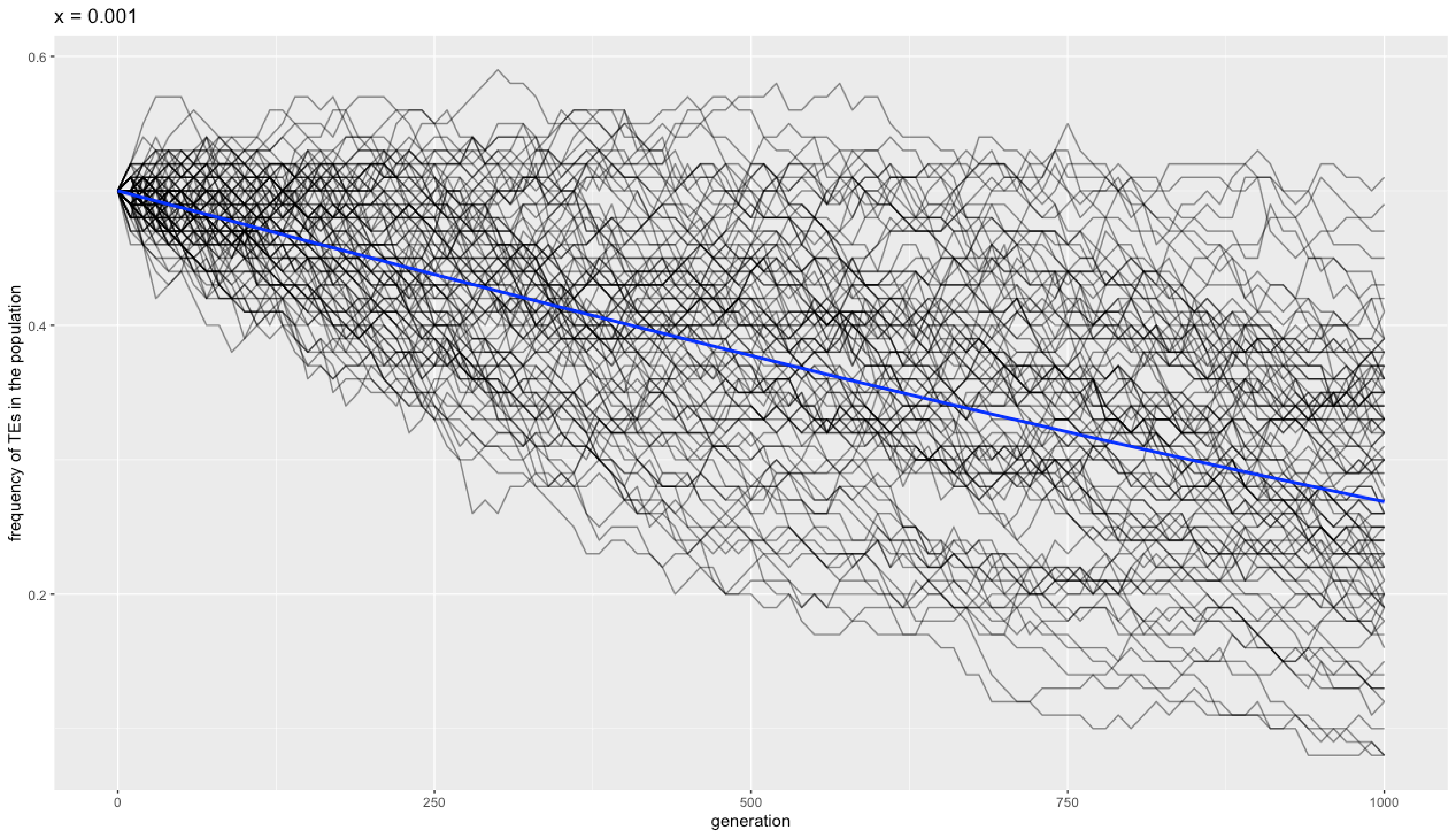

### Validation_6j_selection.pdf

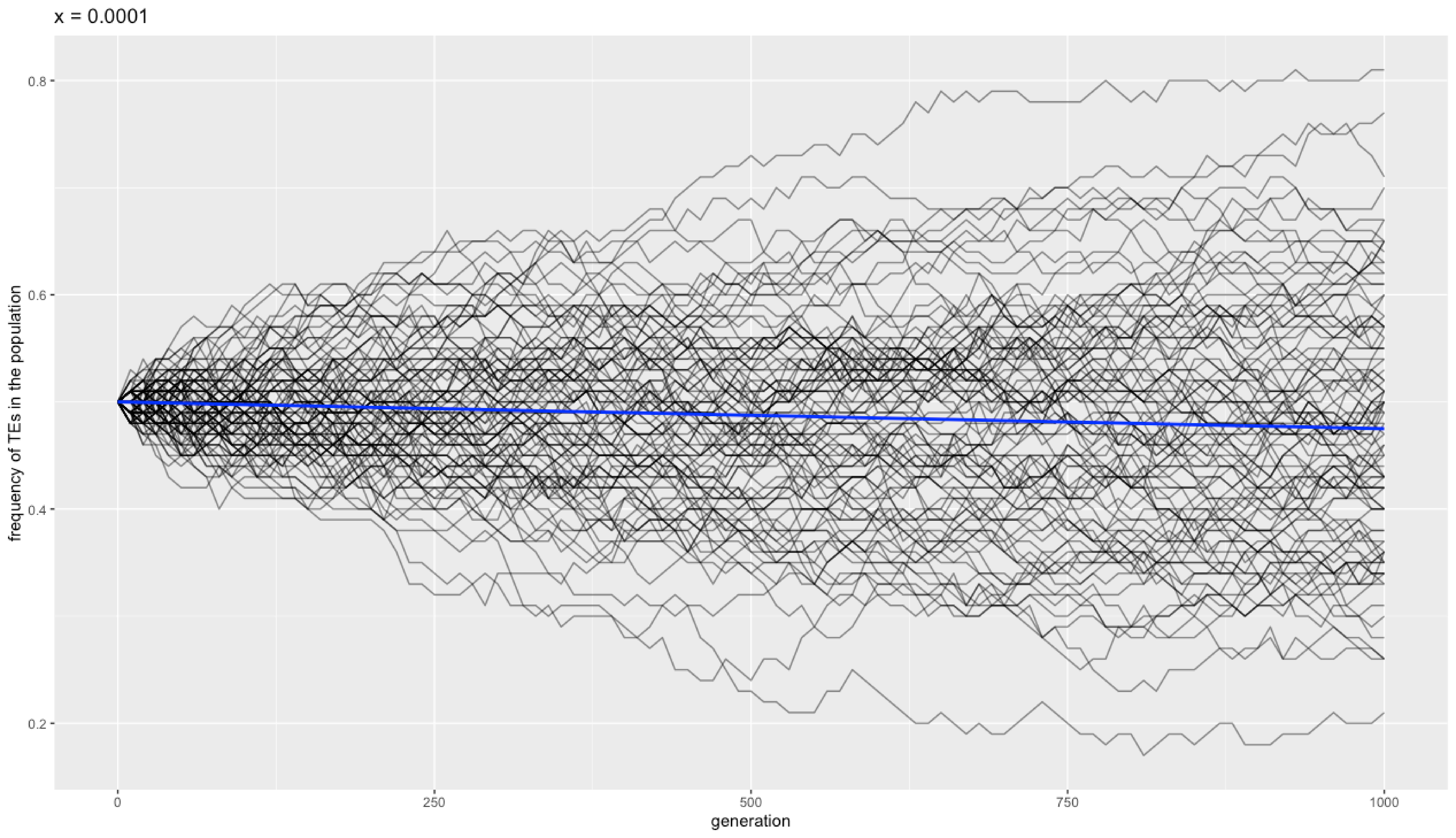

### Validation_6k_selection.pdf

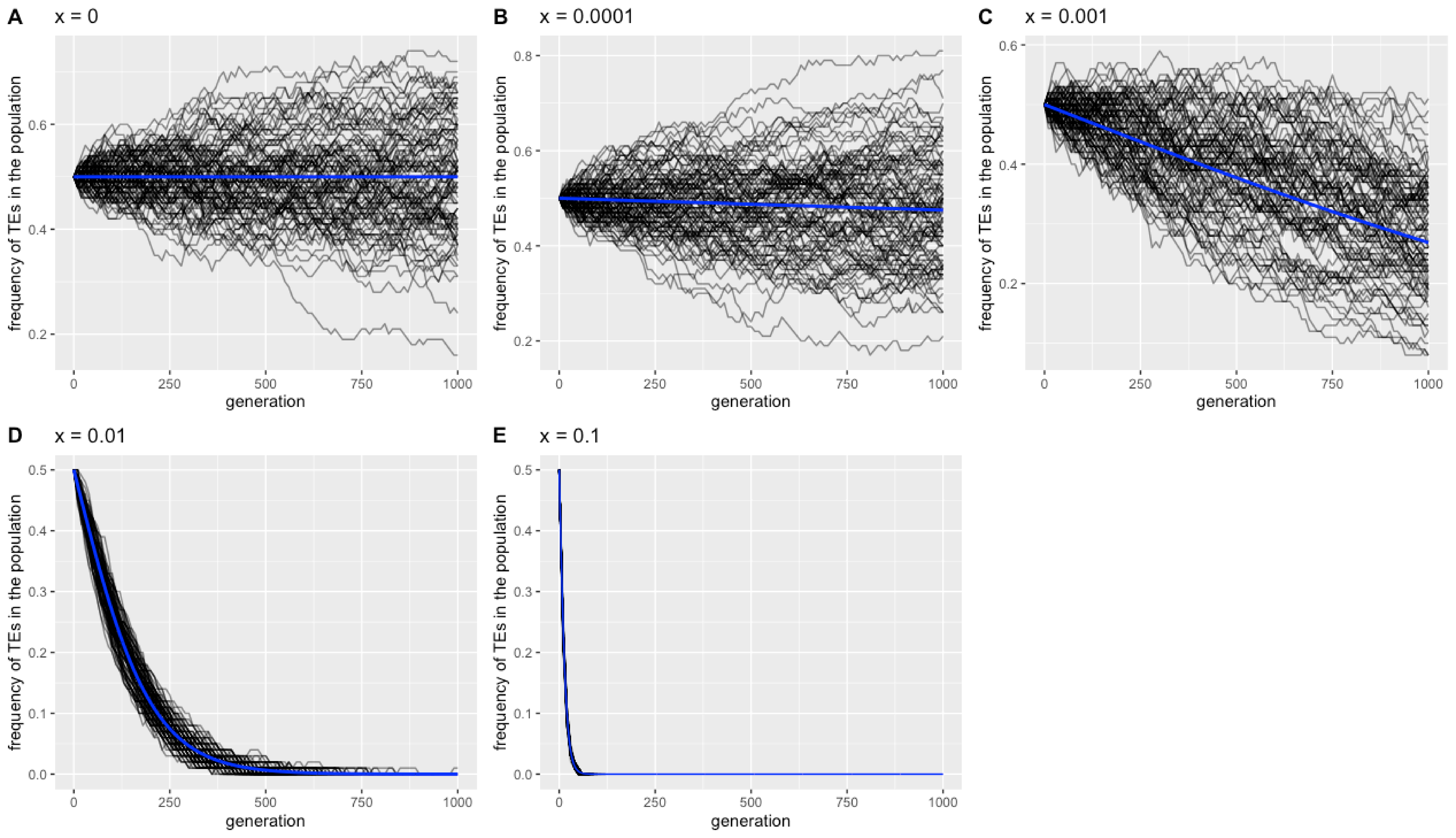

### Validation_7_1A.pdf

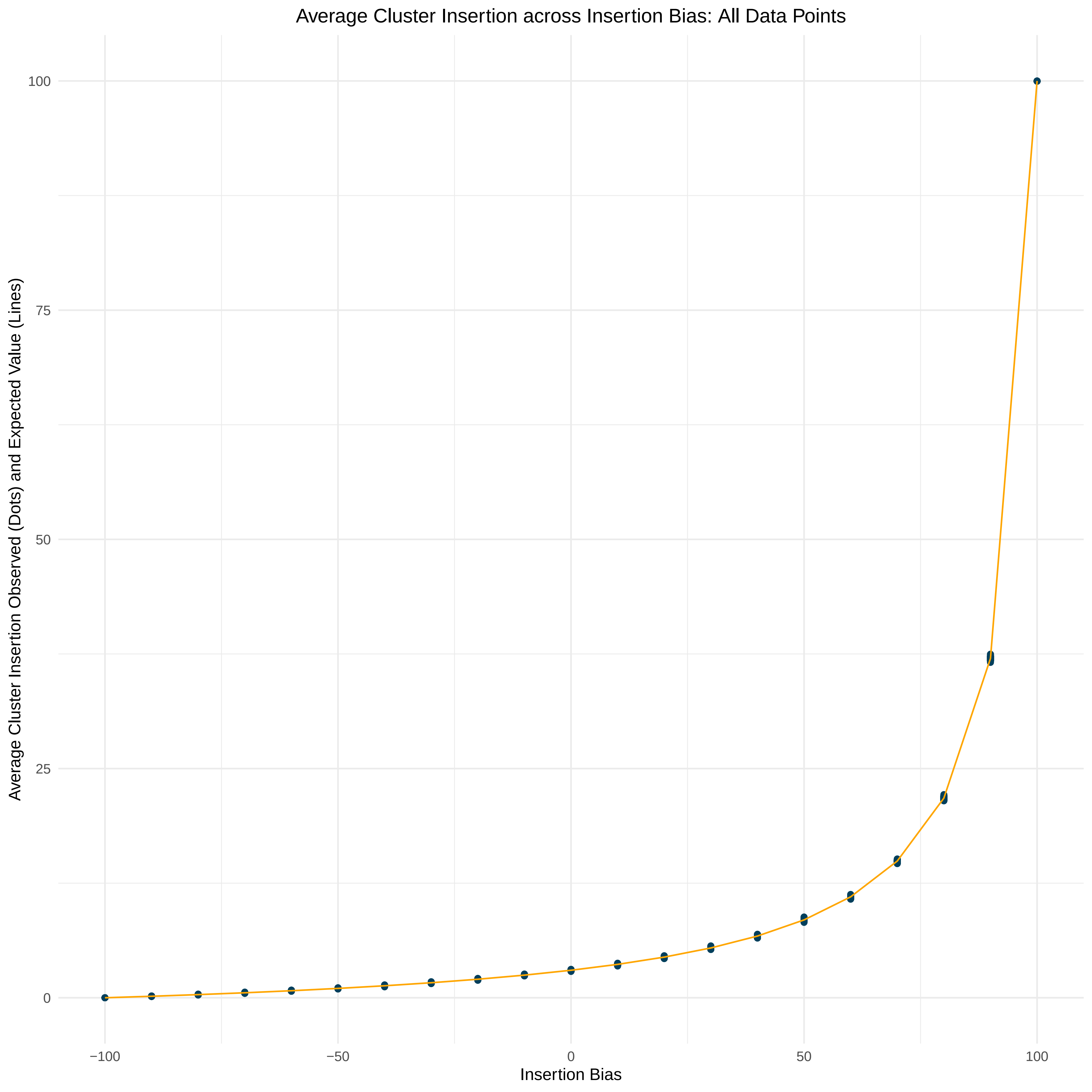

### Validation_7_1B.pdf

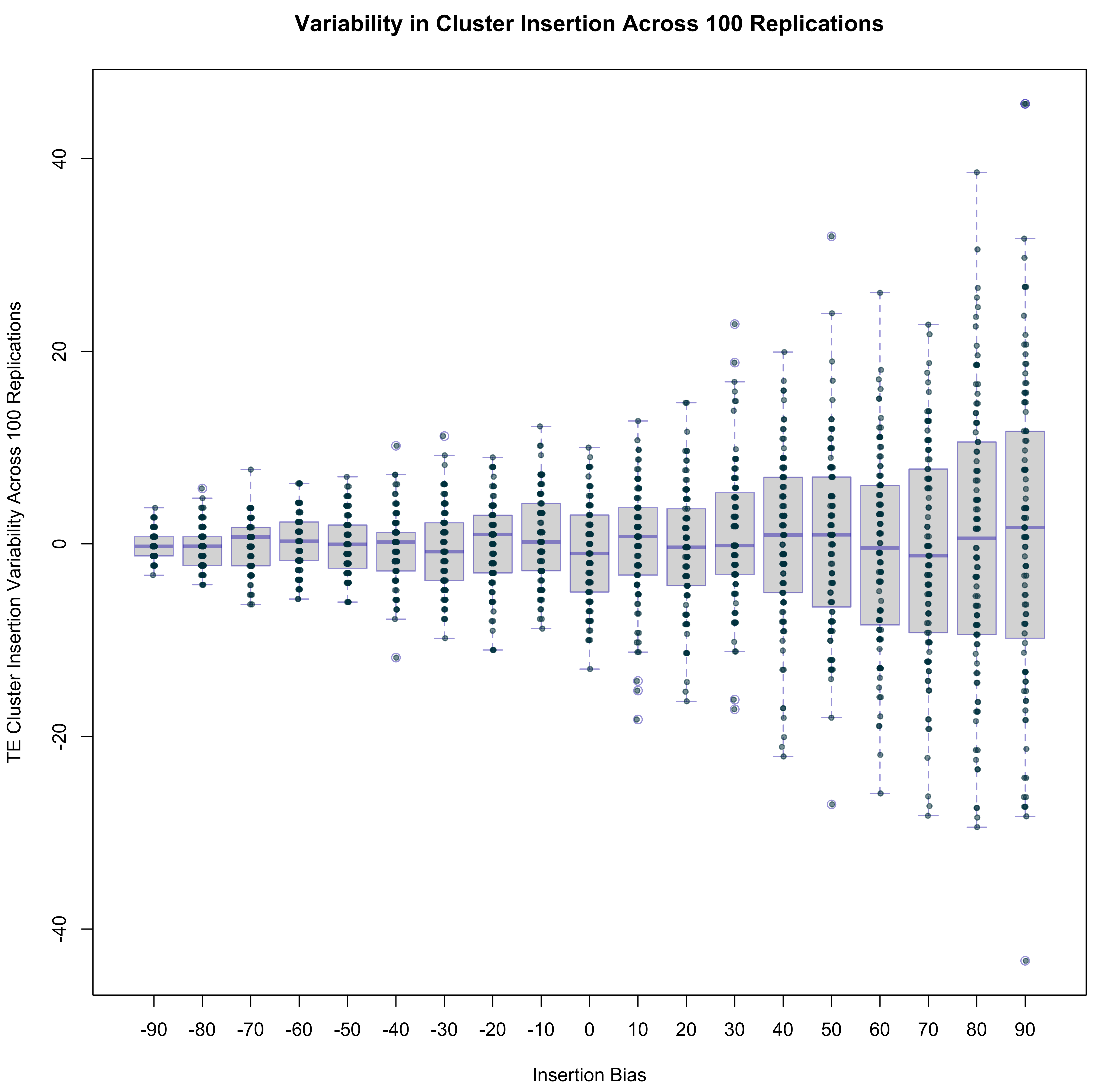

### Validation_7_1C.pdf

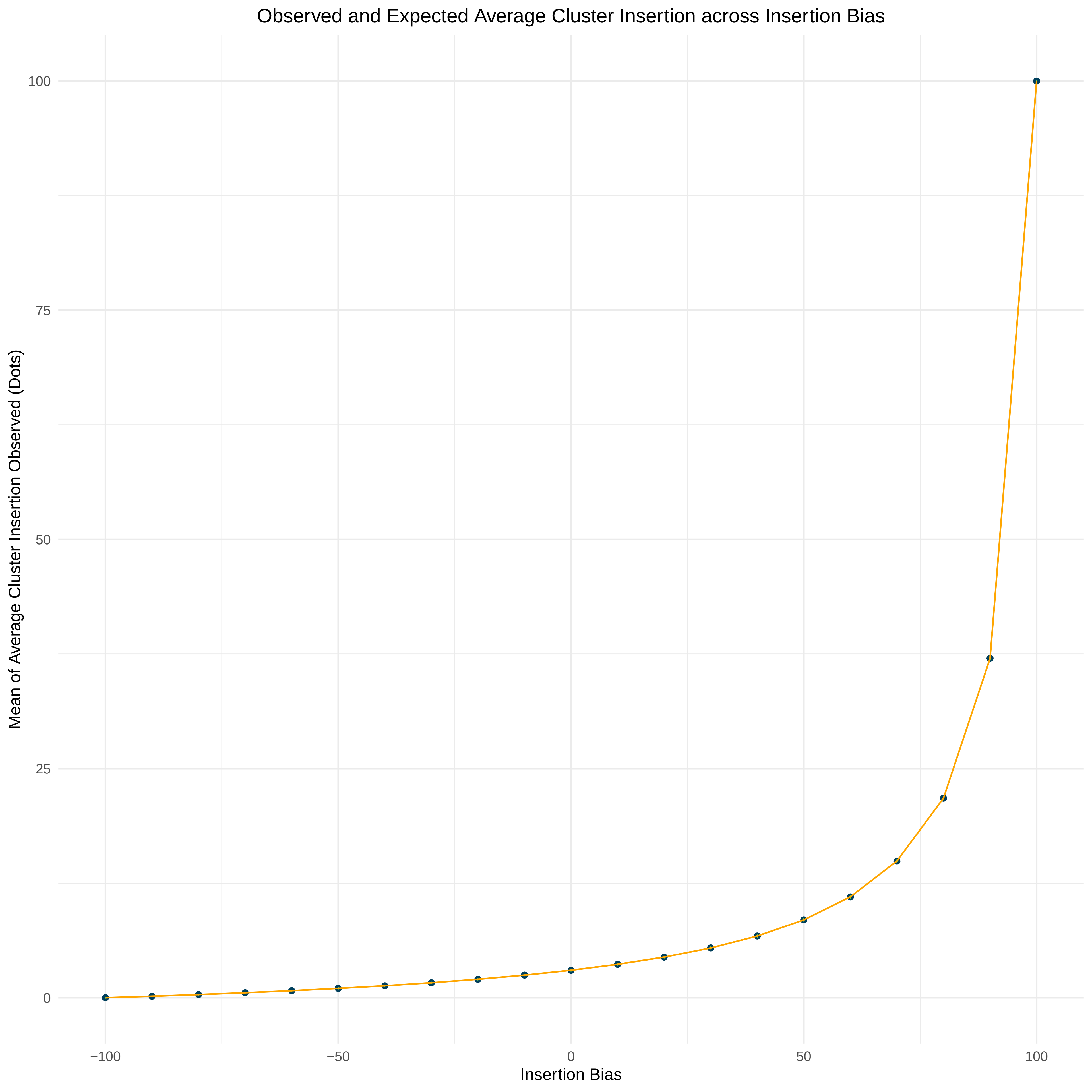

### Validation_7_1D.pdf

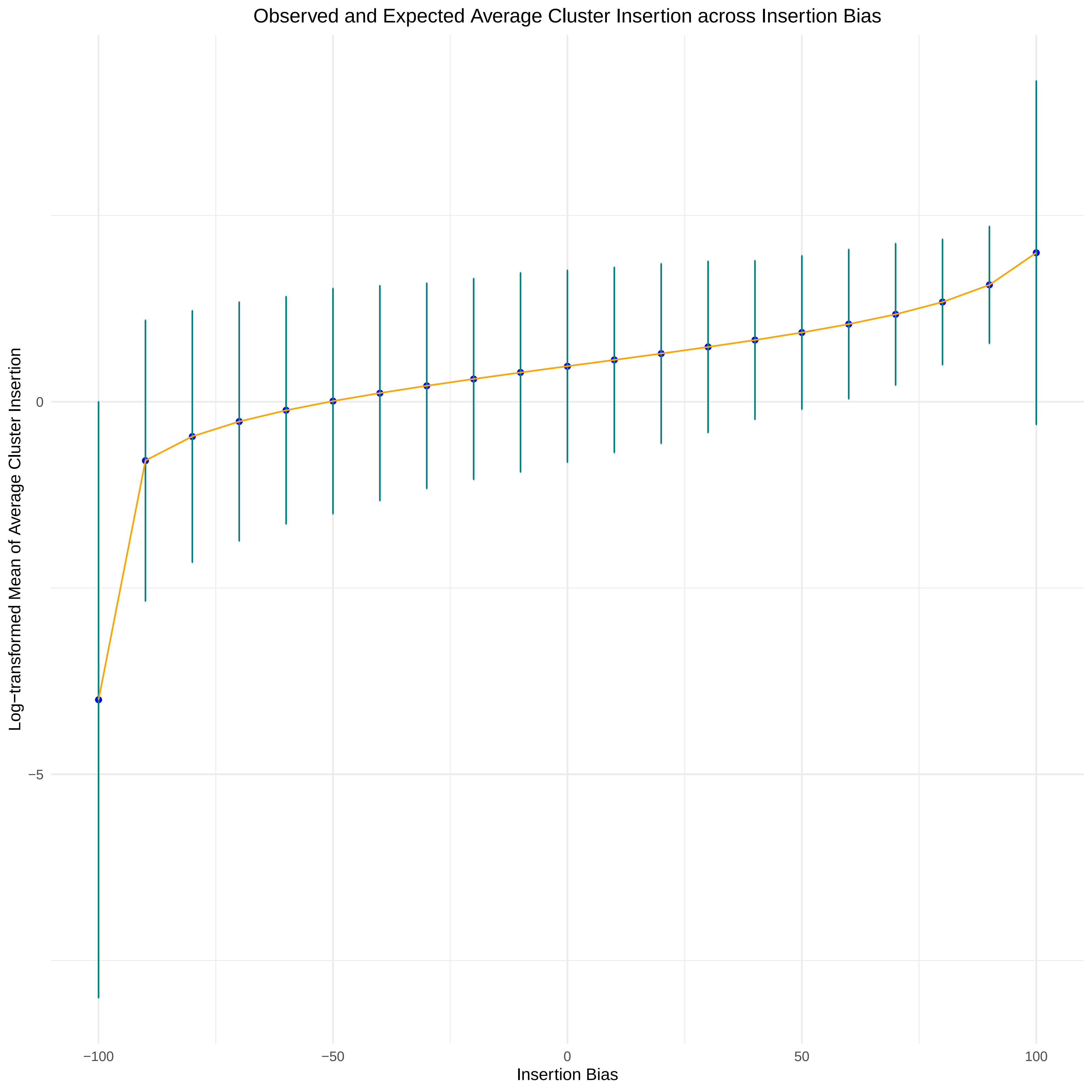

### Validation_7_1E.pdf

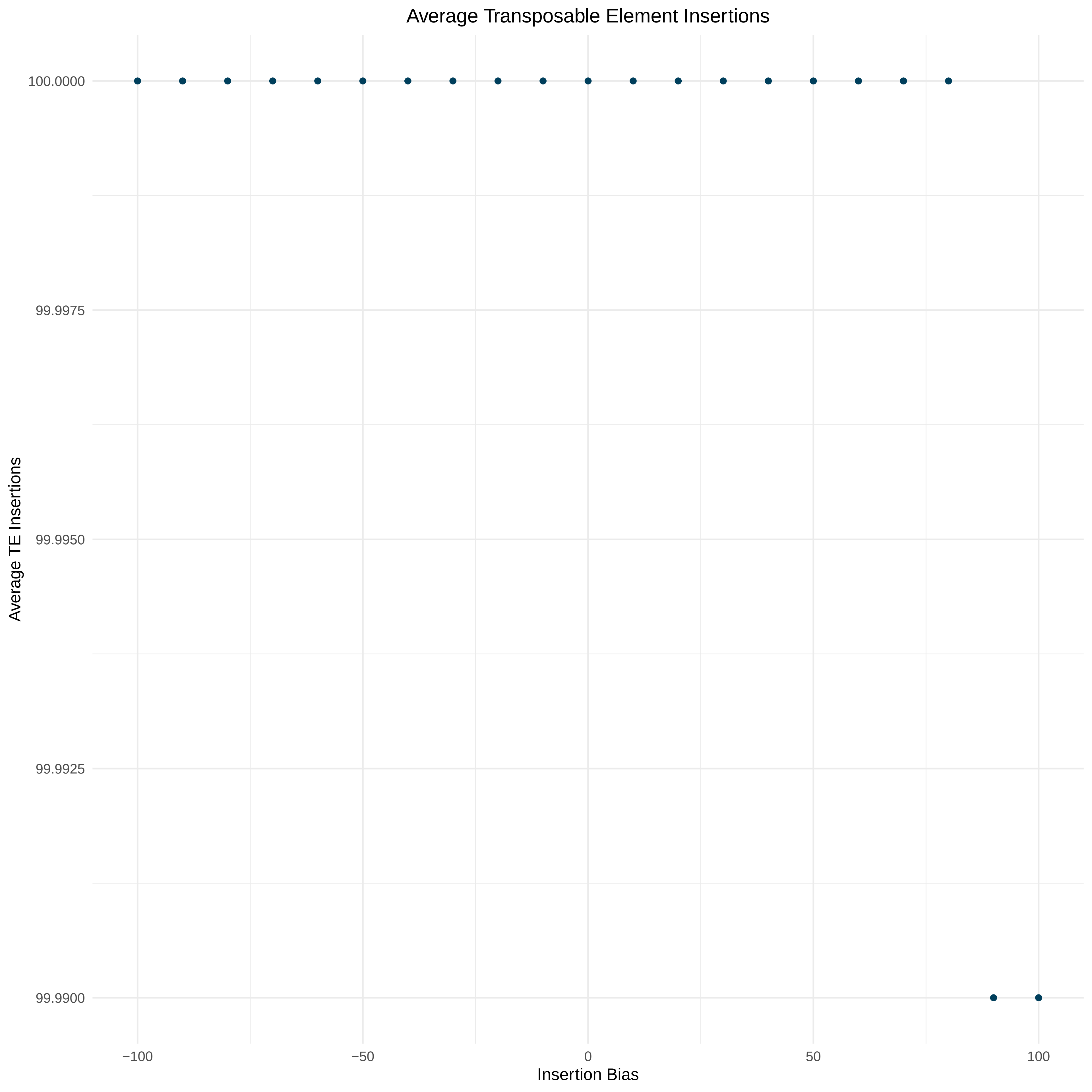

### Validation_7_1F.pdf

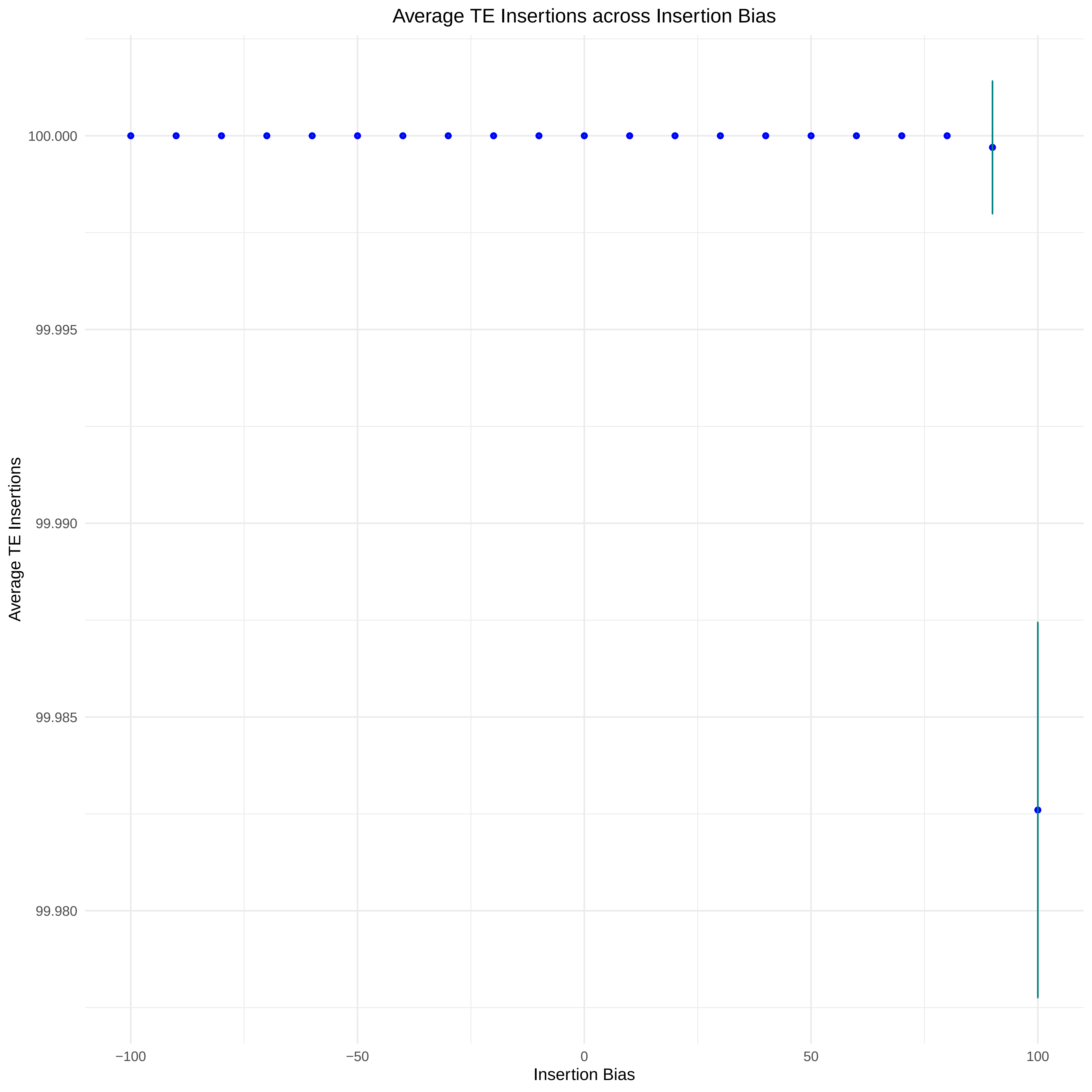
